# Long-read assembly of a Great Dane genome highlights the contribution of GC-rich sequence and mobile elements to canine genomes

**DOI:** 10.1101/2020.07.31.231761

**Authors:** Julia V. Halo, Amanda L. Pendleton, Feichen Shen, Aurélien J. Doucet, Thomas Derrien, Christophe Hitte, Laura E. Kirby, Bridget Myers, Elzbieta Sliwerska, Sarah Emery, John V. Moran, Adam R. Boyko, Jeffrey M. Kidd

## Abstract

Technological advances have allowed improvements in genome reference sequence assemblies. Here, we combined long- and short-read sequence resources to assemble the genome of a female Great Dane dog. This assembly has improved continuity compared to the existing Boxer-derived (CanFam3.1) reference genome. Annotation of the Great Dane assembly identified 22,182 protein-coding gene models and 7,049 long non-coding RNAs, including 49 protein-coding genes not present in the CanFam3.1 reference. The Great Dane assembly spans the majority of sequence gaps in the CanFam3.1 reference and illustrates that 2,151 gaps overlap the transcription start site of a predicted protein-coding gene. Moreover, a subset of the resolved gaps, which have an 80.95% median GC content, localize to transcription start sites and recombination hotspots more often than expected by chance, suggesting the stable canine recombinational landscape has shaped genome architecture. Alignment of the Great Dane and CanFam3.1 assemblies identified 16,834 deletions and 15,621 insertions, as well as 2,665 deletions and 3,493 insertions located on secondary contigs. These structural variants are dominated by retrotransposon insertion/deletion polymorphisms and include 16,221 dimorphic canine short interspersed elements (SINECs) and 1,121 dimorphic long interspersed element-1 sequences (LINE-1_Cfs). Analysis of sequences flanking the 3’ end of LINE-1_Cfs (*i*.*e*., LINE-1_Cf 3’-transductions) suggests multiple retrotransposition-competent LINE-1_Cfs segregate among dog populations. Consistent with this conclusion, we demonstrate that a canine LINE-1_Cf element with intact open reading frames can retrotranspose its own RNA and that of a SINEC_Cf consensus sequence in cultured human cells, implicating ongoing retrotransposon activity as a driver of canine genetic variation.

**Significance:** Advancements in long-read DNA sequencing technologies provide more comprehensive views of genomes. We used long-read sequences to assemble a Great Dane dog genome that provides several improvements over the existing reference derived from a Boxer dog. Assembly comparisons revealed that gaps in the Boxer assembly often occur at the beginning of protein-coding genes and have a high-GC content, which likely reflects limitations of previous technologies in resolving GC-rich sequences. Dimorphic LINE-1 and SINEC retrotransposon sequences represent the predominant differences between the Great Dane and Boxer assemblies. Proof-of-principle experiments demonstrated that expression of a canine LINE-1 could promote the retrotransposition of itself and a SINEC_Cf consensus sequence in cultured human cells. Thus, ongoing retrotransposon activity may contribute to canine genetic diversity.

## Introduction

The domestic dog (*Canis lupus familiaris*) is an established model system for studying the genetic basis of phenotype diversity, assessing the impact of natural and artificial selection on genome architecture, and identifying genes relevant to human disease. The unique genetic structure of dogs, formed as a result of trait selection and breed formation, has particularly aided genetic mapping of dog traits (1, 2).

Canine genetics research has taken advantage of a growing collection of genomics tools including high density single nucleotide polymorphism (SNP) arrays, thousands of genome sequences acquired with short-read technologies, the existence of rich phenotype information, and the availability of DNA obtained from ancient samples (3). This research has relied on the reference genome, CanFam, derived from a Boxer breed dog named Tasha and originally released in 2005 (4). The CanFam assembly was constructed at the end of the first phase of mammalian genome sequencing projects and used a whole-genome shotgun approach that included the end-sequencing of large genomic DNA inserts contained within bacterial artificial chromosome (BAC) and fosmid libraries in conjunction with a limited amount of finished BAC clone sequence (4). Subsequent analyses of CanFam and other genomes sequenced in this manner have suggested that there is an incomplete representation of duplicated and repetitive sequences in the resultant assemblies. Although multiple updates have improved the CanFam assembly, yielding the current CanFam3.1 reference assembly (5), numerous assembly errors, sequence gaps, and incomplete gene models remain. Thus, a more complete and comprehensive dog genome will aid the identification of mutations that cause phenotypic differences among dogs and enable continued advances in comparative genomics (6).

Genome analyses have revealed that canine genomes contain an elevated number of high GC-content segments relative to other mammalian species (7, 8). Genetic recombination may contribute to the evolution of these segments. Studies across a number of mammalian species have indicated that genetic recombination events cluster in specific regions known as hotspots (9). In many species, the PRDM9 zing-finger protein binds to specific nucleotide sequences to promote the initiation of recombination (10-12). In addition to cross-overs, the molecular resolution of recombination events involves gene conversion (13). Gene conversion shows a bias in favor of copying G/C sequences and makes an important contribution to the evolution of genome content (14, 15). Due to gene conversion events and changes in the DNA binding domains of PRDM9, the locations of recombination hotspots in many species are not stable over evolutionary time (16, 17). However, dogs lack a functional PRDM9 protein (18), and canine recombination maps indicate that recombination events are concentrated in GC-rich segments that reside near gene promotors (19, 20). Thus, the observed distribution of GC-rich sequence segments in the canine genome may be a consequence of the stable recombination landscape in canines.

Analysis of the CanFam3.1 reference has demonstrated that a large fraction of the dog genome has resulted from the expansion of transposable elements belonging to the short and long interspersed element (SINE and LINE) families. Fine mapping has implicated mobile element insertions, and associated events such as retrogene insertions, as the causal mutation underlying morphological differences, canine diseases, and selectively bred phenotypes (21-31). A comparison between the Boxer-derived CanFam reference and a low coverage (∼1.5x) draft genome from a Poodle identified several thousand dimorphic copies of a recently active lysine transfer RNA (tRNA)-derived canine SINE element, SINEC_Cf, implying a variation rate 10-100 fold higher than observed for still active SINE lineages in humans (32). Similarly, insertions derived from a young canine LINE-1 lineage, L1_Cf, were found to be >3-fold more prevalent than L1Hs, the active LINE-1 lineage found in humans (32, 33). However, the assembly of long repetitive sequences with a high nucleotide identity is technically challenging, leaving many LINE sequences incorrectly represented in existing reference genomes. Consequently, the biological impact of these elements has remained largely unexplored and the discovery of dimorphic canine LINE-1 sequences is limited to a few reports (32, 34, 35).

Following the era of capillary sequencing, genome reference construction shifted toward high coverage assemblies that utilized comparatively short-sequencing reads. These approaches offered a great reduction in cost and an increase in per-base accuracy, but still were largely unable to resolve duplicated and repetitive sequences, often yielding assemblies that contained tens of thousands of contigs (36). Methods based on linked-read or chromosome conformation sequencing are capable of linking the resulting contigs into larger scaffolds, including entire chromosome arms, but these scaffolds are typically littered with sequence gaps reflecting the poor representation of repetitive sequences (37-39). Here, we analyze the genome of a female Great Dane named Zoey that we sequenced using PacBio long-read technology. We integrated this long-read data with additional sequencing resources, including standard high coverage short-read sequence data, as well as sequence data derived from mate-pair and pooled fosmid libraries, to generate a high-quality assembly. Using this new assembly, we annotate novel gene structures and GC-rich sequences that are absent from CanFam3.1 and under-represented in existing Illumina canine short-read sequence datasets. We demonstrate that gaps in the CanFam3.1 assembly are enriched with sequences that have an extremely high GC content and that overlap with transcription start sites and recombination hotspots. We identify thousands of mobile element insertions, including intact LINE-1 copies, and make use of our fosmid library to subclone an intact L1_Cf element. We demonstrate that a cloned canine L1_Cf is capable of high levels of retrotransposition of its own mRNA (*in cis*) and can drive the retrotransposition of a consensus SINEC_Cf RNA (*in trans*) in cultured human cells. Our analysis provides a more complete view of the canine genome and demonstrates that the distribution of extremely GC-rich sequences and the activity of mobile elements are major factors affecting the content of canine genomes.

## Results

### Long-read assembly of a Great Dane genome

We performed a genome assembly of a female Great Dane, Zoey, using multiple genome sequencing resources that included a standard Illumina short-read sequencing library, a 3 kb Illumina mate-pair sequencing library, sequences from a pooled fosmid library, and ∼50X raw long-read coverage generated using the PacBio RSII system. PacBio long reads were assembled using the Falcon assembler (40), yielding 2,620 primary contigs longer than 1 kbp that encompassed 2.3 Gbp of sequence. In addition, 6,857 secondary contigs, with a total length of 178.5 Mbp, that represent the sequence of heterozygous alleles were assembled (see *Supplementary Information*, Section 1).

The assembly process is based on detecting overlaps among sequencing reads. As a result, reads that end in long stretches of sequence which map to multiple genomic locations and that have high sequence identity, can give rise to chimeric contigs that falsely conjoin discontinuous genomic segments. Using Illumina mate-pair and fosmid pool data from Zoey, clone end sequences from the Boxer Tasha, and alignments to the existing CanFam3.1 assembly, we identified 20 contigs that appeared to be chimeric. We split these contigs at the chimeric junctions, yielding a total of 2,640 contigs with an N_50_ length of 4.3 Mbp and a maximum contig length of 28.8 Mbp. As expected, alignment against the CanFam3.1 assembly indicated comprehensive chromosome coverage (Figure 1). Consistent with the problems in assembly caused by segmental duplications, we found that long contigs (> 3 Mbp) ended in duplicated sequence greater than 10 kbp more often than expected by chance (p<0.001 by permutation, see *Supplementary Information*, Section 1).

**Figure 1:**
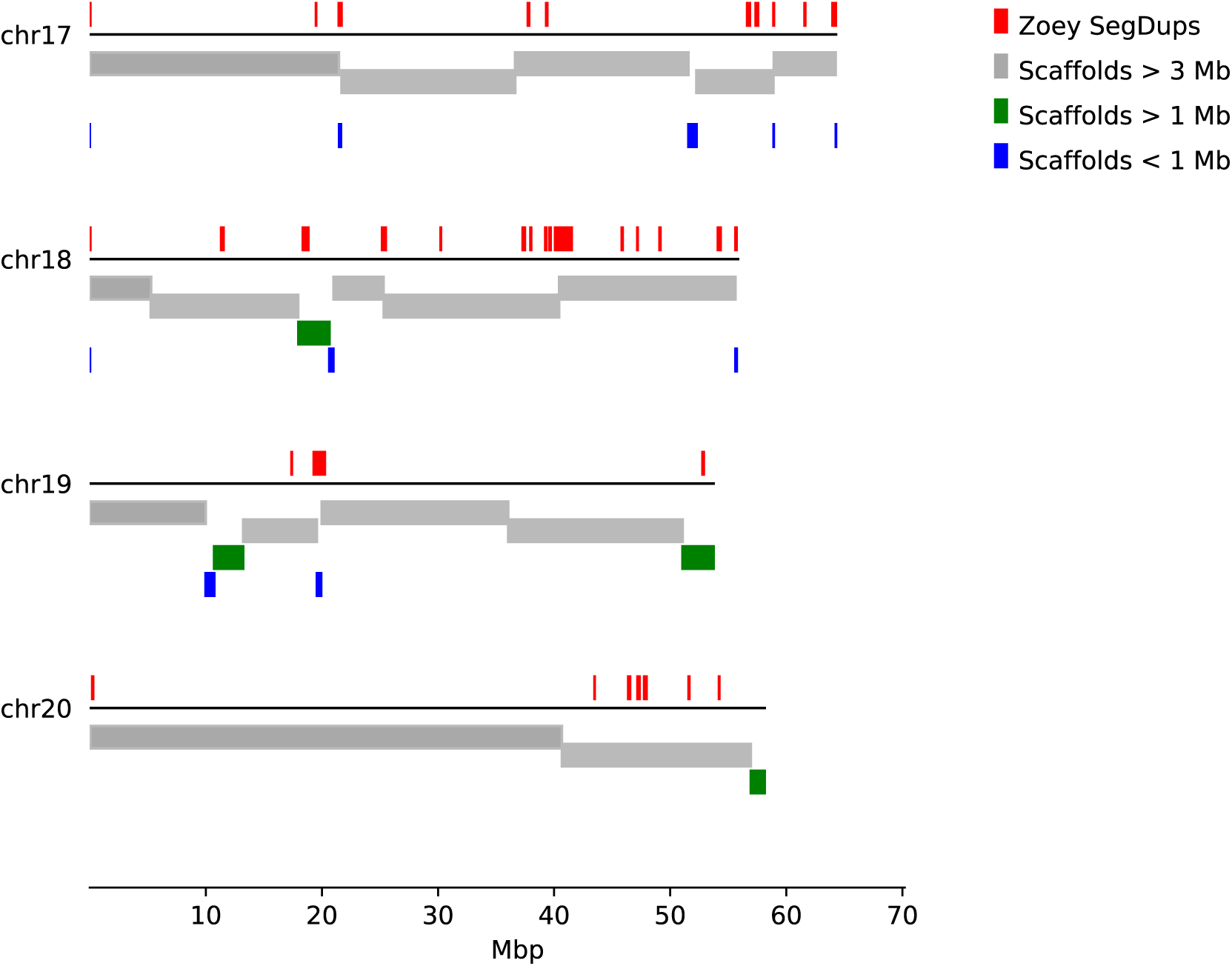
Alignment of assembled contigs to the CanFam3.1 genome. Each of the 2,640 primary contigs were aligned to the CanFam3.1 reference genome. Results are shown for four chromosomes. The colored bars below each line indicate the corresponding position of each contig, colored based on their indicated length. Above each line, regions of segmental duplications based on read depth in the Zoey Illumina data are indicated by red boxes. Permutation tests indicate that long contigs end at regions of segmental duplication more often than expected by chance. See *Supplementary Information*, Section 1 for additional details.

Alignment of the 2,640 contigs and the raw PacBio reads against the CanFam3.1 assembly revealed apparent gaps between contigs, many of which were spanned by PacBio reads. Reasoning that these reads may have been excluded from the assembly due to length cutoff parameters used in the Falcon pipeline, we performed a locus specific assembly utilizing the Canu assembler (v1.3) (41.). This process yielded 373 additional contigs with a total length of 10.5 Mbp and a N_50_ length of 30 kbp. Based on the mapping of the Zoey derived mate-pair sequences and end sequences from the Tasha-derived fosmid and BAC libraries, we linked the 2,640 primary contigs and 373 gap-filling contigs into scaffolds (42). Gap-filling contigs that were not linked using paired reads were excluded from further analysis, resulting in a total of 1,759 scaffolds with a N_50_ of 21 Mbp. Scaffolds were assigned to chromosomes and ordered based on alignment to CanFam3.1. Sequences that appeared to represent allelic variants based on sequence identity and read depth were removed, yielding a chromosomal representation that included 754 unlocalized sequences (see *Supplementary Information*, Section 1).

### Annotation of genome features

We identified segmental duplications in the Zoey and CanFam3.1 assemblies based on assembly self-alignment (43) and read-depth (44) approaches (see *Supplementary Information*, Section 3). Although the number of duplications is similar in each genome, the Zoey assembly contains a smaller total amount of sequence classified as “duplicated”, which likely reflects the continued challenges in properly resolving duplications longer than 10 kbp (Table 1) (45). To compare the large scale organization of the assemblies, we constructed reciprocal liftOver tracks that identify corresponding segments between the CanFam3.1 and Zoey assemblies (46). Based on this comparison, we identified 44 candidate inversions >5kb. Of these candidate large inversions, 68% (30 of 44) were associated with duplicated sequences. The X chromosome, which contributes 5.3% of the genome length, contains 41% (18/44) of the predicted inversions.

**Table 1:** Comparison of the Boxer and Great Dane assemblies. Presented are general assembly statistics for the primary autosomal and X chromosome sequence of the CanFam3.1 and Zoey assemblies. Contiguous segment refers to the length of sequence uninterrupted by an ‘N’ nucleotide. Segmental duplications were identified in each assembly based on an assembly self-alignment and by the depth of coverage of Illumina sequencing reads from Penelope, an Iberian Wolf. See *Supplementary Information*, Section 3 for additional details.

|  | **CanFam3.1**<br>**autosomes + X** | **Zoey**<br>**autosomes + X** |
| --- | --- | --- |
| Total length | 2,327,633,984 | 2,326,329,672 |
| *non-N* | 2,317,593,971 | 2,320,292,846 |
| Number of gaps | 19,553 | 997 |
| Longest contiguous segment | 2,428,071 | 28,813,894 |
| Mean contiguous segment length | 118,523 | 2,239,665 |
| Median contiguous segment length | 54,641 | 1,107,836 |
| N<sub>50</sub> segment length | 277,468 | 4,765,928 |
| Segmental Duplications<br>Genome Alignment, >1 kbp, >90% ID | 6,250 | 6,371 |
| *bp* | 49,339,683 | 45,425,166 |
| Segmental Duplications<br>Penelope Read Depth | 459 | 468 |
| *bp* | 47,757,534 | 40,836,807 |

We created a new gene annotation based on previously published RNA sequencing data using both genome-guided and genome free approaches (47-49) (see *Supplementary Information*, Section 2). Following filtration, this process resulted in a final set of 22,182 protein coding gene models; forty-nine of these gene models are absent from the CanFam3.1 assembly. Full-length matches were found for only 84.9% (18,834) of all protein-coding gene models, while near-full length alignments were found for 93% (20,670) of the models. We additionally annotated 7,049 long non-coding RNAs (50), including 84 with no or only partial alignment to CanFam3.1. Using existing RNA-Seq data (5), we estimated expression values for each protein-coding gene across eleven tissues and report the results as tracks on a custom UCSC Genome Browser assembly hub (51) (Figure 2). The assembly hub illustrates correspondence between the CanFam3.1 and Zoey assemblies and displays the annotation of additional features including structural variants, segmental duplications, common repeats, and BAC clone end-sequences (see *Supplementary Information*, Section 7).

**Figure 2:**
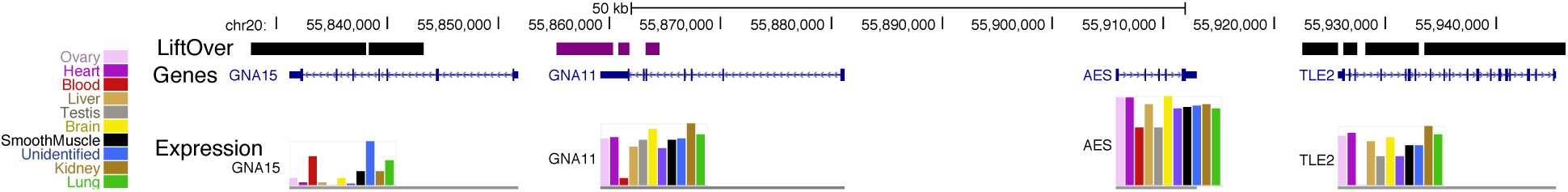
Annotation of genes missing from the CanFam3.1 assembly. Shown is a genome browser view of chr20 on the Zoey assembly is shown. The top track summarizes a comparison between the Zoey and CanFam3.1 assemblies using the UCSC liftOver tool. Black segments show alignment to the corresponding chromosome on the CanFam3.1 assembly. Purple segments match to an unlocalized contig (chrUn_JH374124) in the CanFam3.1 assembly. The large region in the middle between the purple and black segments is absent from the CanFam3.1 assembly. The track below shows the position of four genes in this region annotated using RNA-Seq data: *GNA15, GNA11, AES*, and *TLE2*. The colored bars below each gene model show the expression levels across different tissues, as indicated by the color key in the left of the figure. See *Supplementary Information*, Section 2 for additional details.

### Resolved assembly gaps include GC-rich segments underrepresented in Illumina libraries

Alignment indicates that 12,806 of the autosomal gaps in CanFam3.1 are confidently localized to a unique location in the Zoey genome assembly. In total, 16.8% (2,151) of the gap segments overlap with a transcription start site of a protein-coding gene, which makes it possible to better understand the importance of these previously missing sequences in canine biology (5, 52). Surprisingly, analysis of unique k-mer sequences that map to the CanFam3.1 gap sequences suggested that these DNA segments often are absent from existing Illumina short-read data sets, even though analysis of DNA from the same samples using a custom array comparative genomic hybridization platform indicates their presence. Interrogation of read-pair signatures also suggest that these sequences are systematically depleted in Illumina libraries, which is due to their extreme GC-rich sequence composition (see *Supplementary Information*, Section 4).

The sequences corresponding to gaps in CanFam3.1 have an extremely high GC content, with a median GC content of 67.3%, a value substantially higher than the genome-wide expectation of 39.6% (Figure 3). Given the relationship between GC content and recombination in dogs (19, 20), we examined the distance between CanFam3.1 gap sequences and recombination hotspots. We found that 11.8% of gap segments (1,457 of 12,304 on the autosomes) are located within 1 kbp of a hotspot, compared to only 2.9% of intervals expected by chance. These patterns are driven by a subset of segments that have the most extreme GC content. We identified 5,553 segments with a GC content greater than that obtained from 1,000 random permutations. These extreme-GC segments span a total of 4.03 Mbp in the Zoey assembly, have a median length of 531 bp, a median GC content of 80.95%, and are located much closer to transcription start sites (median distance of 290 bp) and recombination hotspots (median distance of 68.7 kbp) than expected by chance.

**Figure 3:**
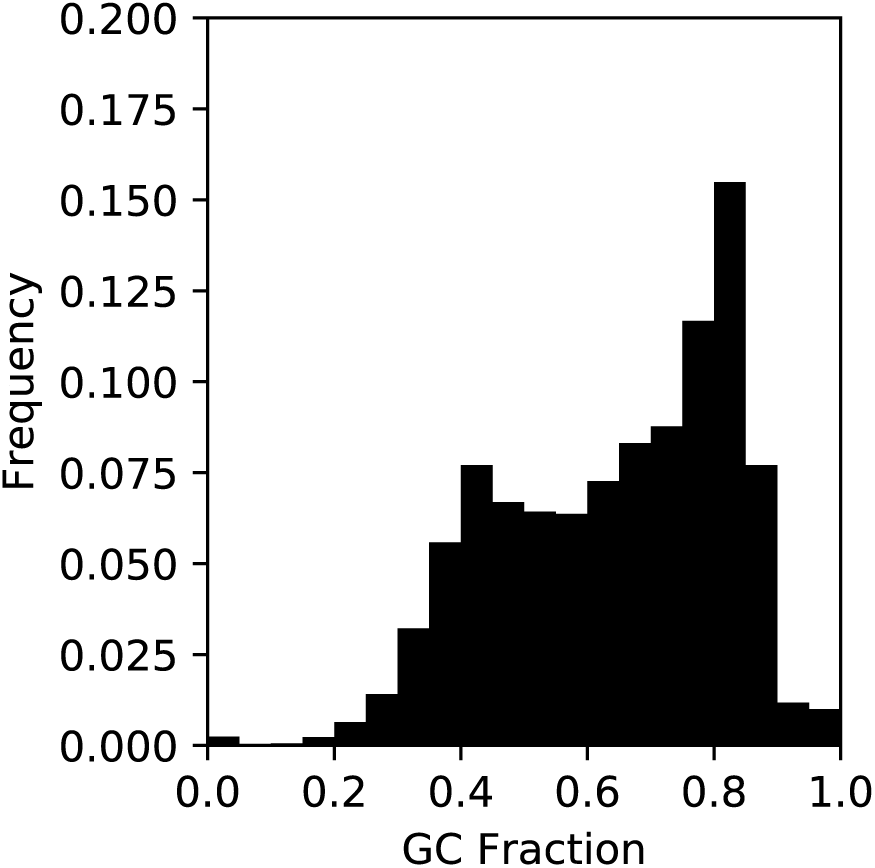
CanFam3.1 assembly gaps are enriched for sequence with extreme GC content. Depicted is the distribution of GC content for 12,806 resolved assembly gaps. A subset consisting of 5,553 of the 12,806 segments have a GC content greater than that found in 99% of randomly selected segments. See *Supplementary Information*, Section 4 for additional details.

### Mobile elements account for the majority of structural differences between canine genomes

We compared the CanFam3.1 and Zoey assemblies to identify insertion-deletion differences at least 50bp in length. After filtering variants that intersect with assembly gaps or segmental duplications, we identified 16,834 deletions (median size: 207 bp) and 15,621 insertions (median size of 204 bp) in the Zoey assembly relative to CanFam3.1 (see *Supplementary Information*, Section 5). In total, these structural variants represent 13.2 Mbp of sequence difference between the two assemblies. The length distribution of the detected variants shows a striking bimodal pattern with clear peaks at ∼200 bp and ∼6 kbp, consistent with the size of canine SINEC and LINE-1 sequences (Figure 4). We inspected the sequence of the events in the 150-250 bp size range and found that 7,298 deletions and 6,071 insertions were dimorphic SINEC sequences. Additionally, LINE-1 sequences accounted for 339 deletions and 581 insertions longer than 1 kbp.

**Figure 4:**
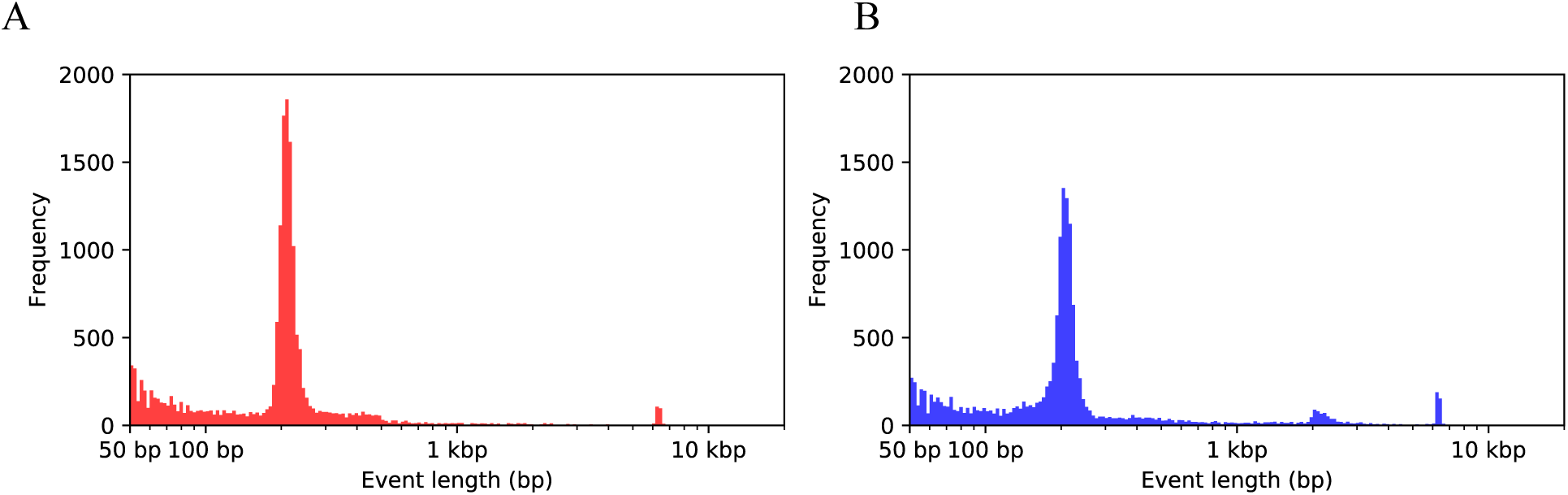
Size of structural variants identified between CanFam3.1 and Zoey assemblies. Shown are histograms depicting the size distribution of 16,834 deletions (panel A) and 15,621 insertions (panel B) between the Zoey and CanFam3.1 assemblies. Variant size is plotted on a logarithmic scale such that the bins in the histogram are of equal size in the log scale. Large increases at ∼200bp and ∼6kbp indicate the disproportionate contribution of dimorphic LINE1 and SINEC sequences to the genetic differences between the two assemblies. See *Supplementary Information*, Section 5 for additional details.

Our assembly also contains 6,857 secondary contigs, which represent alternative sequences at loci where Zoey is heterozygous for a structural variant. Alignment of these secondary contigs against the CanFam3.1 assembly yielded an additional 2,665 deletion and 3,493 insertion events, encompassing a total of 2.67 Mbp of sequence. We further inspected the sequence of these variants and found 1,259 deletions and 1,593 insertions consistent with dimorphic SINEC elements, and 75 deletions and 126 insertions consistent with dimorphic LINE-1 elements. Together, comparison of the Zoey and CanFam3.1 genomes identified at least 16,221 dimorphic SINEC and 1,121 dimorphic LINE-1 sequences (see *Supplementary Information*, Section 5).

LINE-1 transcription often bypasses the polyadenylation signal encoded within the element, resulting in the inclusion of flanking genomic sequence in the LINE-1 RNA (53-55). Thus, after retrotransposition, the resulting 3’-transductions can be used as sequence signatures to identify the progenitor source elements of individual LINE-1 insertions (56, 57). We identified 18 transduced sequences among the dimorphic LINE-1 sequences in our data set. Of these transduced sequences, 17 aligned elsewhere in the genome at a location that is not adjacent to an annotated LINE-1. This includes a pair of LINE-1 copies on chr25 and chrX which share the same transduced sequence, as well as a locus on chr19 that has the same transduction as a duplicated sequence present on chr2 and chr3. Such “parentless” 3’-transductions suggest the presence of additional dimorphic LINE-1 sequences that are capable of retrotransposition (see *Supplementary Information*, Section 5).

### Canine genomes contain LINE-1s and SINEs capable of retrotransposition

The high degree of dimorphic LINE-1 and SINEC sequences found between the two assemblies suggests that mobile element activity represents a mutational process that is ongoing in canines. The canine LINE-1 (L1_Cf) consensus sequence contains segments of GC-rich sequence and homopolymer runs, including a stretch of 7 ‘C’ nucleotides in the ORF1p coding sequence that likely are prone to errors incurred during DNA replication, PCR, and sequencing. Thus, a bioinformatic search for L1_Cf sequences with intact open reading frames is biased by uncorrected sequencing errors. We therefore searched the Zoey assembly for sequences that have long matches with low sequence divergence from the L1_Cf consensus. The Zoey assembly encodes 837 L1_Cf sequences that have less than 2% divergence and are greater than 99.4% of full-length; an additional 169 elements are present on the secondary contigs. This set includes 187 full length LINE-1s, of which, 31 were found in secondary contigs. For comparison, these values represent a 65% increase over the 113 elements present in CanFam3.1 that meet the same criteria (see *Supplementary Information*, Section 5).

To more thoroughly characterize canine LINE-1 copies that may remain active, we isolated and sequenced individual fosmids from Zoey predicted to contain full-length LINE-1s. We identified one sequence, from fosmid clone 104_5 on chr1 (L1_Cf-104_5), possessing intact open readings frames, which encode the ORF1p and ORF2p predicted proteins, that lack mutations expected to disrupt protein function (see *Supplementary Information*, Section 6). We subcloned this element for functional analysis in a cultured cell assay that uses an indicator cassette that is only expressed following a successful round of retrotransposition (58-60), yielding G418-resistnat foci. We found that the L1_Cf-104_5 element is capable of retrotransposition of its own mRNA *in cis* in human HeLa cells (Figure 5 and *Supplementary Information*, Section 6).

**Figure 5:**
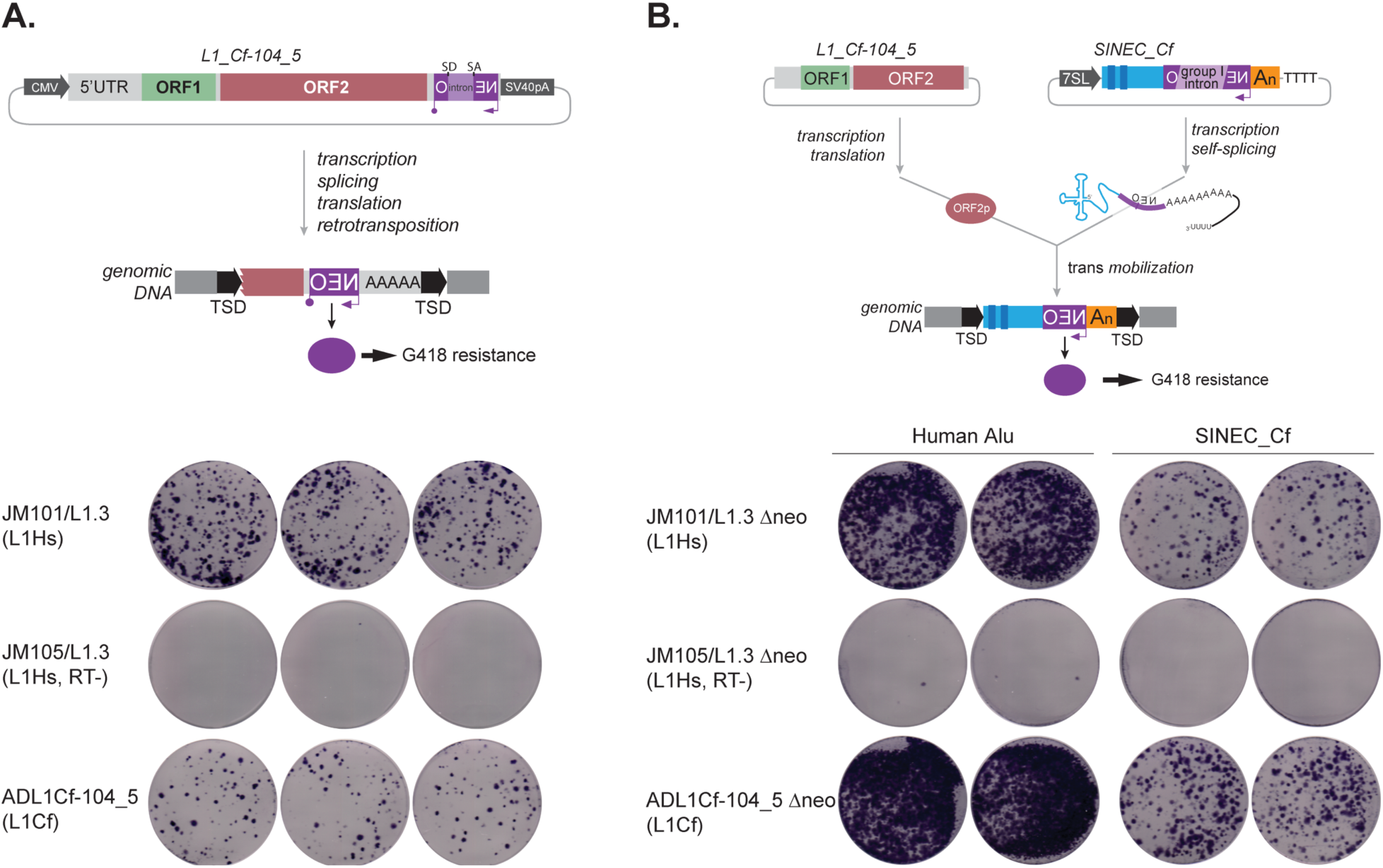
Identification of canine LINE-1 and SINEC elements capable of retrotransposition. (Panel A, top): A full length L1_Cf equipped with a retrotransposition indicator cassette (*mneoI*) was assayed for retrotransposition in human HeLa-HA cells. TSD, indicates a target site duplication generated upon retrotransposition. (Panel A, bottom): Results of the retrotransposition assay. JM101/L1.3 (positive control) contains an active human LINE-1. JM105/L1.3 (negative control) contains a human LINE-1 that harbors an inactivating missense mutation in the reverse transcriptase domain of ORF2p (85). ADL1Cf-104_5 contains the full-length canine LINE-1 identified in this study. (Panel B, top): A consensus SINEC_Cf element equipped with an indicator cassette to monitor the retrotransposition of RNA polymerase III transcripts (*neo*^tet^) (61) was assayed for retrotransposition in human HeLa-HA cells in the presence of either an active human LINE-1 or the newly cloned L1_Cf-104_5 sequence that lacks a retrotransposition indicator cassette (JM101/L1.3Δneo or ADL1Cf-104_5Δneo, respectively). (Panel B, bottom): Results of the retrotransposition assay. JM101/L1.3Δneo (positive control) contains an active human LINE-1. JM105/L1.3 Δneo (negative control) contains a human LINE-1 that harbors an inactivating missense mutation in the reverse transcriptase domain of ORF2p (85). ADL1Cf-104_5Δneo contains an active canine LINE-1 (see panel A). The expression of either JM101/L1.3Δneo or ADL1-Cf-105Δneo could drive human Alu and SINEC_Cf retrotransposition. In both assays, the blue stained foci represent G418-resistant foci containing a presumptive retrotransposition event. See *Supplementary Information*, Section 6 for additional details.

SINE sequences are non-autonomous elements that utilize the function of LINE-1 ORF2p to mediate their retrotransposition *in trans* (60, 61). To test the capability of L1_Cf-104_5 to mobilize SINE RNA in *trans*, we constructed a second reporter vector containing the SINEC_Cf consensus sequence marked by an appropriate indicator cassette (61). We found that expression of L1_Cf-104_was capable of mobilizing both canine SINEC and human Alu RNAs in *trans* (Figure 5 and *Supplementary Information*, Section 6).

## Discussion

Due to their unique breed structure, history of selection for disparate traits, and extensive phenotypic data, dogs are an essential model for dissecting the genetic basis of complex traits and understanding the impact of evolutionary forces on genome diversity. The era of long-read sequencing is revolutionizing genomics by enabling a more complete view genomic variation (62). Here, we describe the assembly and annotation of the genome of Great Dane dog and compare it with the Boxer-derived CanFam3.1 reference assembly. Comparison of our Great Dane genome to the CanFam3.1 reference revealed several key findings important to canine genome biology. Several other long-read assemblies of canines are planned or have been recently released (63, 64). The availability of these resources, along with other long-read canine that assembles that are planned or have been recently released (63, 64), will provide significant benefits to the canine genomics community.

Our Great Dane assembly has improved sequence continuity, resolves novel gene structures, and identifies several features important to canine genome biology. For example, we created a new gene annotation that includes 49 predicted protein coding genes that are absent from the CanFam3.1 reference genome. Our analysis also identified 2,151 protein-coding gene models whose transcription start position corresponds to a gap in the CanFam3.1 assembly. This finding largely resolves prior observations that many dog genes appear to have incomplete first exons and promoters (5, 6). Analysis of the Great Dane assembly further revealed that gaps in the CanFam3.1 assembly are enriched for sequence that has extremely high-GC content, providing a probable explanation of their absence from the CanFam3.1 assembly (65).

The presence of extremely GC-rich segments likely reflects a key aspect of canine genome biology. In contrast to humans and many other mammals, genetic recombination in canines is targeted towards gene promoter regions due to the absence of a functional *PRDM9* gene (20). In other species, the PRDM9 protein binds to specific nucleotide sequences and targets the initiation of recombination to distinct loci in the genome. It has been hypothesized that recombination in dogs is instead localized by general chromatin marks, which are associated with promoters, resulting in a fine-scale genetic map that is more stable over evolutionary time (19, 20). In addition to crossing-over, recombination events result in gene conversion, a process with a bias in favor of G/C alleles. Biased gene conversion can be modeled as positive selection in favor of G/C alleles at a locus (14, 15, 66) and has been previously proposed as an explanation for the unusual GC content of the dog genome (19, 20). Our analysis indicates that the GC rich segments associated with recombination hotspots are larger than expected previously. These expanded segments have an unknown effect on the expression of their associated gene, have been largely absent from previous genome assemblies, and are depleted from Illumina sequencing data. A more extensive examination of the long-term consequence of stable recombination hotspots on genome sequence structure will require assessment of genomes of other species which lack PRDM9 using long-read technologies.

Long-read sequencing offers a less biased view of structural variation between genomes, particularly for insertions (67). The profile of genomic structural variation between the Zoey and CanFam3.1 assemblies is dominated by dimorphic SINEC and LINE-1 sequences, with 16,221 dimorphic SINEC and 1,121 dimorphic LINE-1 sequences. Although analogies between humans and dogs can be problematic (68), a comparison with humans illustrates the magnitude of the mobile element diversity found between the Great Dane and Boxer genome assemblies. In terms of human mobile element diversity, the 1000 Genomes Project estimates that humans differ from the reference genome by an average of 915 Alu insertions and 128 LINE-1 insertions (69). A recent study collated these findings, along with other published data sets, and identified a total of 13,572 dimorphic Alu elements in humans (70), though we note that these estimates are based on Illumina sequencing data, which has limitations in mapping to repetitive regions and in fully capturing insertion alleles (67). Finally, an approach specifically designed to identify dimorphic human LINE-1 insertions utilizing long-read sequencing data identified 203 non-reference insertions in the benchmark sample NA12878, of which 123 which were greater than 1 kbp in length (71).

Illumina sequencing data indicate that Zoey differs from the CanFam3.1 reference at 3.57 million single nucleotide variants (SNVs). This number is lower than the number of differences typically found in a globally diverse collection of human genomes (4.1-5.0 million SNVs) (69), and is comparable to the number found in the NHLBI TOPMed data set (72) (median of 3.3 million SNVs among 53,831 humans sequenced as part of the National Heart, Lung, and Blood Institute’s Trans-Omics for Precision Medicine program). Relative to the number of SNVs, the level of LINE-1 and SINEC dimorphism we found between two dog genomes is disproportionately large. This total represents an approximately 17-fold increase in SINE differences (16,221/915) and an eight-fold increase in LINE differences (1,121/128) compared to the numbers found among humans. Remarkably, more dimorphic SINEs were found between these two breed dogs than have been found in studies of thousands of humans (70, 73). Our data will aid systematic studies of the potential contribution of these elements to canine phenotypes, including cancers (74).

Further study is required to determine the relative contribution of (i) new insertions in breeds or populations versus (ii) the assortment of segregating variants that were present in the progenitor populations. However, our study suggests that retrotransposition is an ongoing process that continues to affect the canine genome. We provide proof-of-principle evidence that dog genomes contain LINE-1 and SINEC elements that are capable of retrotransposition in a cultured cell assay. We also identified two LINE-1 lineages with the same 3’-transduced sequence associated with multiple elements, suggesting the presence of multiple canine LINE-1s that are capable of spawning new insertions. Additionally, analysis of 3’-transduction patterns suggests the presence of additional active LINE-1s in canines that have yet to be characterized. Thus, a full understanding of canine evolution and phenotypic differences requires consideration of these important drivers of genome diversity.

## Methods

Genome assembly and analysis utilized long-read and short data from a female Great Dane named Zoey, a pooled fosmid library (75) constructed from Zoey, sequence data generated from a female Boxer, named Tasha, as part of the CanFam genome assembly (4), and results from a custom comparative genomics hybridization array (array-CGH). Data accessions and detailed methods are available in the Supplementary Information.

Genome assembly of ∼50-fold whole-genome, single-molecule, real-time sequencing (SMRT) data from Zoey was performed on DNAnexus using the FALCON 1.7.7 pipeline (40) and the Damasker suite (76). Chimeric contigs were identified based on mapped reads from the Zoey mate-pair jumping library, the Zoey fosmid pools, the Tasha BAC end sequences, and Tasha fosmid end sequences. Regions that showed a lack of concordant paired end coverage were identified as potential chimeric junction sites and split apart prior to scaffolding. Primary contigs were supplemented with contigs obtained from a local assembly of reads aligning to gaps between contigs on CanFam3.1 using Canu v1.3 (41). Contigs were linked into scaffolds using mapping of the Zoey mate -data, Tasha BAC end sequence data and Tasha fosmid data using the BESST scaffolding algorithm (version 2.2.7)(42) and assigned to chromosomes based on alignment to CanFam3.1. Chain files for use with the UCSC liftOver tool were constructed based on blat (46) alignments. A UCSC TrackHub hosting the Zoey assembly, as well as relevant annotations of both the Zoey assembly and CanFam3.1 is available at https://github.com/KiddLab/zoey_genome_hub

Common repeats in both the CanFam3.1 and Zoey assemblies were identified using RepeatMasker version 4.0.7 with option ‘–species dog’, using the rmblastn (version 2.2.27+) search engine and a combined repeat database consisting of the Dfam_Consensus-20170127 and RepBase-20170127 releases. Self-alignment analysis of each assembly was performed using SEDEF (43) with default parameters. Results were filtered for alignments at least 1kb in length and at least 90% sequence identity. Read-depth analysis was performed using fastCN as described previously (44). Copy-number estimates were constructed in non-overlapping windows each containing 3 kbp of unmasked sequence. Segmental duplications were identified as runs of four windows in a row with an estimated copy-number >=2.5. To provide an unbiased assement of duplication content, read-depth analysis was performed based on Illumina data form Penelope, an Iberian Wolf, in addition to sequences from Zoey and Tasha.

Forty-two canine RNA-Seq runs representing eleven tissue types were used to annotate genes in the Zoey genome (5). *De novo* gene models were created based on alignment of RNA-Seq reads using Cufflinks (v2.2.1) (47, 48) and in a non-reference guided fashion using Trinity (v2.3.2) (49). Gene models were merged and annotated using PASA-Lite (77) and the transdecoder pipeline (version 5.0.1) (78). Gene names and functional annotations were determined using BLAST2GO (79). Expression levels for each of the 22,182 protein-coding gene models were estimated using Kallisto (version 0.46.0) (80). Long non-coding RNAs in the Zoey genome were identified using the FEELnc program (50).

To identify large insertion and deletion variants, the Zoey assembly and 6,857 secondary contigs, were aligned to CanFam3.1 using minimap2 (version 2.9-r720) with the -asm5 option (81). The output from the alignment was parsed using the paftools.js program released as part of minimap2 to identify candidate variants. Breakpoint coordinates were refined by performing targeted alignment of the flanking and variant sequence for each candidate using AGE (82).

Individual fosmids containing potentially full-length L1_Cf elements were isolated from pools using a lifting procedure coupled with hybridization of a probe containing digoxygenin (DIG) labeled dUTP. Isolated fosmids were sequenced in small pools via RS II PacBio sequencing and assembled using the HGAP2 software (83). An intact L1_Cf was subcloned from fosmid 104_5, equipped with an *mneoI* retrotransposition indicator cassette, and tested for retrotransposition in HeLa-HA cells (58, 59). The construction of the L1_Cf expression vector and the conditions used to assay for retrotransposition are detailed in *Supplemental Information*, Section 6.

To monitor SINEC_Cf mobilization, we modified the *Alu neo*^tet^ vector, which contains an active human AluYa5 element equipped with a reporter cassette engineered to monitor the retrotransposition of RNA polymerase III (pol III) transcripts (61). Briefly, the *Alu neo*^tet^ vector consists of a 7SL RNA Pol III enhancer sequence upstream of AluYa5 that is equipped with a ‘backward’ *neo*^R^ gene under the control of an SV40 promoter. The *neo*^R^ gene is disrupted by a tetrahymena self-splicing group I intron that is in the same transcriptional orientation as the Alu element. This arrangement only allows the expression of the *neo*^R^ gene (yielding G418-resistant foci) upon a successful round of retrotransposition in HeLa-HA cells, yielding G418-resistant foci (61). We replaced the AluYa5 sequence with the SINEC_Cf consensus sequence, obtained from Repbase (84). The resultant construct was used to assay SINEC_Cf mobilization, in *trans*, in the presence of either an active human LINE-1 or the newly cloned L1_Cf-104_5 expression plasmid that lacks the retrotransposition indicator cassette (JM101/L1.3Δneo or ADL1Cf-104_5Δneo, respectively). The construction of the SINEC_Cf expression vector and the conditions used to assay for retrotransposition are detailed in *Supplemental Information*, Section 6.

## Competing interests

J.V.M. is an inventor on patent US6150160, is a paid consultant for Gilead Sciences, serves on the scientific advisory board of Tessera Therapeutics Inc. (where he is paid as a consultant and has equity options), and currently serves on the American Society of Human Genetics Board of Directors. A.R.B. is the co-founder and Chief Science Officer of Embark Veterinary.

## Acknowledgments

This work was supported in part by National Institutes of Health (NIH) grant R01GM103961 to J.M.K. and A.R.B., NIH Academic Research Enhancement Award R15GM122028 to J.V.H., and NIH Training Fellowship T32HG00040 to A.L.P. DNA samples were provided by the Cornell Veterinary Biobank, a resource built with the support of NIH Grant R24GM082910, and the Cornell University College of Veterinary Medicine. Additional DNA samples were kindly provided by Brian Davis, Elaine Ostrander, and Linda Gates. We thank Dorina Twigg, Chai Fungtammasan, Brett Hannigan, Mark Mooney, Dylan Pollard, and DNAnexus for assistance with sequence data processing and the University of Michigan Advanced Genomics Core for assistance with data production. We especially thank Linda Gates for her continued devotion to all Great Danes and her assistance with this project.

## Notes

https://github.com/KiddLab/zoey_genome_hub

