## Supplemental Appendix for "Long-read assembly of a Great Dane genome highlights the contribution of GC-rich sequence and mobile elements to canine genomes"

### Supplementary Information

#### Table of Contents

**1. Data Generation and Assembly** **2**
*Data summary and accessions* *2*
*Isolation of individual fosmids* *6*
*Array-CGH analysis* *6*
*Genome assembly* *7*

**2. Gene Annotation** **11**
*RNA-Seq read processing* *11*
*Trinity de novo gene models* *11*
*PASA annotation pipeline* *11*
*Transdecoder* *13*
*Transcript redundancy and filtering* *13*
*Functional annotation of predicted gene models* *14*
*Alignment of gene models to CanFam3.1* *14*
*Quantification of gene expression* *14*
*Long non-coding RNA analysis* *15*

**3. Assembly Annotation** **16**
*Repetitive sequences* *16*
*Segmental duplications* *16*
*QuicK-mer2 analysis* *16*
*Alignment of Tasha clone end-sequences* *17*
*liftOver track generation* *17*
*Inversion analysis* *17*
*Identification of single nucleotide variants in Zoey* *18*

**4. Analysis of CanFam3.1 Sequence Gaps** **19**
*CanFam3.1 gaps resolved in Zoey* *19*
*Copy-number analysis of sequence in CanFam3.1 assembly gaps* *20*
*CanFam3.1 gaps are enriched for sequences with extreme GC content* *26*
*Extremely GC-rich segments are located near transcription start sites and recombination hotspots* *28*

**5. Analysis of Large Insertions and Deletions** **31**
*Filtering and breakpoint refinement* *31*
*Dimorphic mobile element sequences* *32*
*LINE-1 3' transduction analysis* *34*
*Annotation of transduced sequences* *35*
*Comparison of full-length L1_Cf elements in Zoey and CanFam3.1* *45*

**6. Mobile Element Analysis** **47**
*Identification of candidate intact LINE-1 insertions* *47*
*LINE-1 subcloning strategy* *47*
*SINEC_Cf subcloning strategy* *49*
*Cultured cell retrotransposition assay* *51*

**7. Genome Browser TrackHub Resources** **53**
*Accessing the TrackHub* *53*
*Summary of available data* *53*

**8. References** **57**

### 1. Data Generation and Assembly

*Data summary and accessions*

Genome assembly and analysis utilized long-read and short-read data from a female Great Dane, Zoey, a pooled fosmid library from Zoey, sequence data generated from a female boxer, Tasha, used as part of the CanFam assembly, and results from a custom comparative genomics hybridization array (array-CGH).

Small supplementary tables are included in this document. The larger supplementary tables, including Tables 1.1, 2.1, 2.3, 3.1 A-B, 3.2, 3.3, 4.1, 5.2 A-D, 5.3 A-D, 5.6 A-D, 5.7 A-D, and 5.8, are available as separate files in the online appendix.

Zoey Genomic Data

*Illumina sequence data*

Standard Illumina short-read data from Zoey have been published previously (1). Read files are available from the Sequence Read Archive (SRA) under accession SRX1705166 (Figure 1.1).

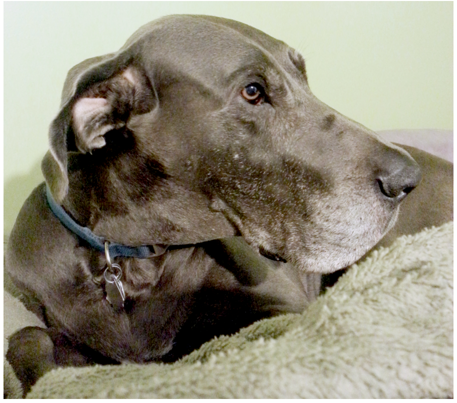

**Figure 1.1: Genomic DNA from a female Great Dane named Zoey was utilized for this study.** Photo provided by Linda Gates.

*Illumina mate-pair jumping library*

A ~3 kbp Illumina mate-pair library was constructed from Zoey genomic DNA following the procedure outlined in the following references (2, 3). Read files are available from the SRA under accessions SRX2677896 and SRX2677885. This procedure results in the sequencing of 27bp long sequences at opposite ends of ~3 kbp molecules that are separated by a defined linker sequence. Scripts for preprocessing the raw fastq files are available at https://github.com/KiddLab/jumps-map.

*Illumina fosmid pool data*

A pooled fosmid library was generated from genomic DNA from Zoey and processed as described previously (4). Alignment of reads from each pool to the CanFam3.1 reference genome identified a total of 333,328 clones across 192 pools with a median clone size of 34,010 bp (Figure 1.2). Read pool data is available from the SRA under accession SRP073312.

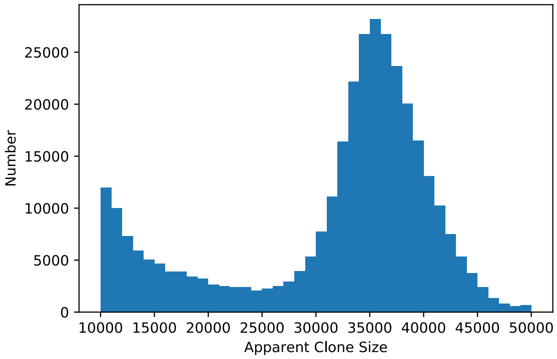


**Figure 1.2: Distribution of fosmid clone sizes used in this study.** The distribution of apparent clone sizes is plotted for 333,328 fosmids identified by mapping reads from each of 192 pools to the CanFam3.1 reference genome.

*PacBio WGS data*

Whole genome sequence data was generated at the University of Michigan Advanced Genomics Core using the PacBio RS II platform with the P6C4 chemistry (6th generation polymerase and 4th generation chemistry). A total of 91 SMRT cells were run from two separate DNA libraries. Read data is available from the SRA under accession SRP150834.

As an initial assessment, raw PacBio reads were aligned to the CanFam3.1 reference genome using BLASR (5). Alignments were filtered to require a primary mapping quality score of at least 20. Due to the presence of circular consensus reads, we assigned the longest mapping read obtained from the same SMRT Cell ZMW as the size of the DNA insert. In total, we achieved nearly 50X sequence depth when all reads are considered (assuming a haploid genome size of 2.4 Gbp). When considering the insert or fragment size, we obtained 28x coverage of unique DNA molecules. The median fragment size is 7.5 kbp with an $L_{50}$ length of approximately 10 kb (Figure 1.3). These data reveal that 50% of the total length sequenced is represented by fragments with a size larger than 10 kb.

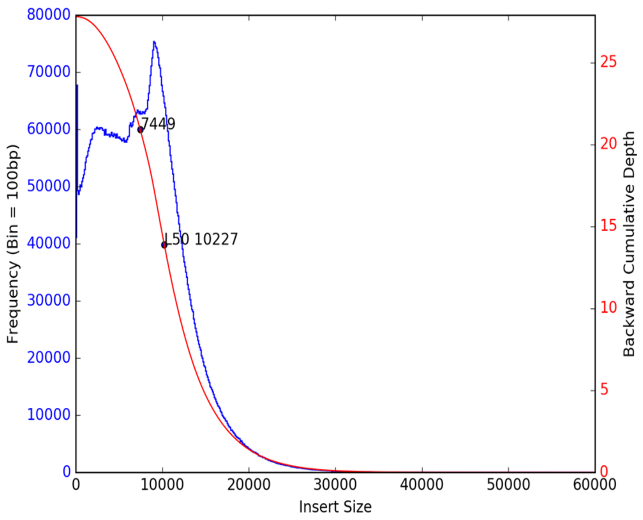


**Figure 1.3: PacBio sequencing fragment insert and coverage statistics.** The blue curve (left axis) shows the distribution of unique fragment inserts aligned to the CanFam3.1 reference genome. The red curve (right axis) shows cumulative depth of individual fragments obtained by adding the longest reads in the pool first. The $L_{50}$ and median fragment sizes are indicated along the red curve.

*Array-CGH data*

Results from 55 array-CGH experiments, including hybridizations of 54 samples against genomic DNA from Zoey, as well as a 'dye swap' experiment, where we hybridized genomic DNA from Zoey against DNA from an Iberian Wolf, Penelope, has been deposited in the Gene Expression Omnibus (GEO) database under accession GSE153608.

Tasha Genomic Data

We utilized Sanger sequencing reads generated from the boxer, Tasha, as part of the original public canine genome assembly effort (6). Reads were downloaded from the NCBI Trace Archive and converted to fastq format using scripts available at: https://github.com/jmkidd/sanger_dl.

*BAC end sequences*

End sequences from the CH-82 canine BAC library (6) were downloaded from the NCBI Trace Archive based on the information listed in: ftp://ftp.ncbi.nih.gov/repository/clone/reports/Canis_familiaris/CH82.endinfo_9615.out
A total of 392,240 paired sequences from 196,120 BAC clones were downloaded and analyzed in this study.

*Fosmid end sequences*

End sequences from a fosmid clone library were downloaded from the NCBI Trace Archive based on the following query:

```
species_code='CANIS LUPUS FAMILIARIS'and INSERT_SIZE =
40000 and CENTER_NAME = 'BI'
```

A total of 3,337,928 paired sequences from 1,668,964 clones were analyzed in this study.

*10 kb plasmid library*

End sequences from a 10 kb plasmid clone library were downloaded from the NCBI Trace Archive based on the following query:

```
species_code='CANIS LUPUS FAMILIARIS' and strain='BOXER'
and CENTER_NAME = 'BI' and trace_type_code='WGS' and
insert_size = 10000
```

A total of 5,299,806 paired sequences from 2,649,903 clones were analyzed in this study.

*4 kb plasmid library*

End sequences from a 4 kb plasmid clone library were downloaded from the NCBI Trace Archive based on the following query:

```
species_code='CANIS LUPUS FAMILIARIS' and strain='BOXER'
and CENTER_NAME = 'BI' and trace_type_code='WGS' and
insert_size != 10000 and insert_size != 40000
```

A total of 25,386,900 paired sequences from 12,693,450 clones were analyzed in this study.

Accessions of assembled data

*Primary assembly*

The primary Zoey assembly is available from GenBank under accession GCA_005444595.1. This assembly can also be explored using a UCSC Assembly Hub which is hosted at: https://github.com/KiddLab/zoey_genome_hub and can be loaded using the following link: https://raw.githubusercontent.com/KiddLab/zoey_genome_hub/master/zoey-genome.hub.txt. Internally, we referred to this sequence as the "Zoey 2.3" assembly, reflecting several iterations of filtering and scaffolding analyses. This sequence is equivalent to the sequence at GCA_005444595.1 which is named UMICH_Zoey_3.1. As described below, the layout of the scaffolds in the Zoey assembly is based on the CanFam3.1 assembly. Fasta sequence records of the primary assembly and secondary contigs are also available at: https://kiddlabshare.med.umich.edu/zoey2.3/.

*Secondary/alternative contigs*

The assembly procedure identified secondary or alternative contigs, which represent the alternative allele at heterozygous structural variant sites. These contigs are available from GenBank under accession SWLD00000000.1 and are also available at https://kiddlabshare.med.umich.edu/zoey2.3/.

*Zoey fosmid clones*

Individual fosmid clones from Zoey were isolated and sequenced using the PacBio RS II sequencer. Clones were sequenced in small pools (~4 fosmid clones per pool) per PacBio run and assembled using the HGAP2 assembler (7). Clone assemblies have been deposited to GenBank under accessions MK829534-MK829589.

*LINE-1 tested in retrotransposition assay*

An L1_Cf element was isolated from fosmid clone 104_5 and was and found to be active in a *cis*-based cultured cell retrotransposition assay (8). The proteins encoded by this L1_Cf element also were capable of mobilizing an active human Alu and the canine SINEC_Cf consensus sequence, obtained from Repbase (9), in a *trans*-based cultured cell retrotransposition assay (10, 11). The sequence of this L1_Cf retrotransposon has been deposited in GenBank under accession MT811810.

*Isolation of individual fosmids*

Individual fosmid clones were isolated from glycerol stocks of clone pools using a lifting procedure followed by hybridization using a clone-specific probe containing digoxygenin (DIG) labeled-dUTP. Hybridization probes were made by amplifying 250-400bp of unique sequence present on the targeted fosmid with the PCR DIG Probe Synthesis Kit, Cat. No. 11 636 090 910 (Sigma-Aldrich, St. Louis, MO). An overnight culture of a fosmid pool containing the clone of interest was diluted 1/10,000. Then 1 ml was plated on a 150 x 20 mm petri dish of LB solid medium supplemented with 25 µg/ml chloramphenicol and cultured overnight at 37˚C. Colony lifts and hybridizations were performed according to the recommended protocols and reagents provided by the manufacturer (https://www.sigmaaldrich.com/content/dam/sigma-aldrich/docs/Roche/General_Information/1/dig-application-manual-for-filter-hybridisation-iris.pdf) using GE Healthcare Amersham™ Hybond™ -N+ membranes (Fisher Scientific, Pittsburgh, PA). A UV Stratalinker (Stratagene, La Jolla. CA) was used to crosslink DNA to the filters.

Growth on petri dishes from diluted overnight cultures was dense, so a plug ~ 0.5cm in diameter centered over a positive clone was selected and rinsed in LB. Then, ~500 cells from this plug were plated on a second petri dish containing solid LB medium supplemented with 25 µg/ml chloramphenicol. The colony hybridization was repeated on this second dish and we isolated single, positive clones. If isolated clones could not be selected, plugs were collected and the colony hybridization was repeated a third time. Isolated clones were streaked and a PCR screen with the primers used to amplify the probe was performed to confirm clone identity and to demonstrate that the resultant clones were pure. Finally, standard capillary end-sequencing was performed on the isolated clones. Clones of interest were sequenced in small pools (4 clones per pool) using a PacBio RSII sequencer and assembled using the HGAP2 software (7).

*Array-CGH analysis*

A custom array-CGH platform (4x170,000 probe format) was designed using the Agilent platform. The array design primarily targeted to regions of predicted copy-number variation in the CanFam3.1 reference genome. Regions targeted on the array included: (i) control genomic regions not detected as copy-number variable; (ii) deletions and duplications predicted by analysis with fastCN, QuicKmer, and Delly (12, 13); (iii) the breakpoints of mobile element insertions predicted using a previously described pipeline (14); (iv) regions predicted to be on the Y chromosome; and (v) novel sequence contigs identified from Illumina short-read data in Zoey and the Iberian Wolf Penelope using the sga assembler (15).

Genomic DNA labeling and array hybridizations were performed using protocols and reagents recommended by the manufacturer: https://www.agilent.com/cs/library/usermanuals/public/GEN-MAN-G4410-90010.pdf. One microgram of genomic DNA was labeled using enzymatic labelling without restriction digest. Arrays were scanned using a NimbleGen MS 200

Microarray Scanner with multi TIFF option and 2 µm resolution. Images were quantified using Agilent Feature Extraction Software version 12.0.2.2. The raw values determined by the Agilent Feature Extraction Software and were normalized to ensure that the mean intensity of the control probes was the same for each hybridization. The array data has been deposited to the Gene Expression Omnibus (GEO) database under accession GSE153608. Illumina WGS data is also available for each of the analyzed samples; accessions are given in Table 1.1.

*Genome assembly*

Genome assembly was performed on DNAnexus using the FALCON 1.7.7 pipeline (16) and the Damasker suite (17). First, approximately 50-fold whole-genome, single-molecule, real-time sequencing (SMRT) data from a female Great Dane, Zoey, were passed through the TANmask and REPmask modules from the Damasker suite. This data then was used as input to the traditional FALCON pipeline, using a length cut-off of 1,775 bp during the initial error-correcting stage. This procedure resulted in 15 million error corrected reads with an $N_{50}$ read length equal to 8.7 kbp and ~38x coverage of the dog genome. Second, the error-corrected reads were again passed through the TANmask and REPmask modules, followed by the overlap portion of the FALCON pipeline. For the overlap portion, we used a length cut-off of 5,087 bp. The aligned reads were assembled in the third stage of FALCON into 2,688 primary contigs containing 2.3 Gbp with an $N_{50}$ contig length of 4.4 Mbp. Finally, the primary contigs were polished using the PacBio Quiver algorithm (7) from SMRT Link 3.1 using the original raw-reads. This procedure yielded an additional 6,857 secondary contigs (total length 178.5 Mb, median 23 kb, $N_{50}$ 28.6 kb) representing alternative alleles.

Next, we searched for potentially chimeric contigs among the 2,688 primary contigs. First, we removed from further analysis any contig less than 1 kbp in length, resulting in 2,620 primary contigs for further consideration. Chimeric contigs can occur during assembly due to the existence of large duplicated or repeated sequence that are not spanned by individual reads. Thus, reads corresponding to distinct copies of the repeated sequence can be erroneously conjoined, resulting in a chimeric contig that erroneously links sequences that actually are dispersed in the genome. We utilized several existing genomic datasets to identify potential chimeric contigs. We mapped reads from the Zoey mate pair jumping library, Zoey fosmid pools, Tasha BAC end sequences, and Tasha fosmid end sequences to the primary contigs and determined the span of coverage of concordant paired-end sequences. Regions that showed a lack of concordant paired end coverage were identified as potential chimeric junction sites. Candidates were further assessed for their total coverage in Zoey Illumina short-read sequencing data and based on alignment of the contigs to the existing CanFam3.1 reference assembly. In total, we split 20 primary contigs into separate pieces prior to scaffolding, resulting in 2,640 contigs. The coordinates for splitting contigs were selected based on the boundaries of contiguous mate-pair and fosmid end-sequence alignment.

We aligned the resulting contigs to the CanFam3.1 assembly using MUMMER (18), BLAT (19), and QuicK-mer2 (20) and aligned the raw PacBio reads to CanFam3.1 using blasr (5) (Figure 1.4). The ends of large contigs tended to correspond to regions of segmental duplication in Zoey. We examined the 514 end points of the 257 contigs longer than 3 Mbp and found that 53 contig ends were placed within 5 kbp of a region identified as a segmental duplication based on Zoey Illumina read depth. To assess how unusual this pattern is, we randomly permutated the position of the duplications 1,000 times and counted the distance to contig endpoints (Figure 1.5). The maximum value observed was 29 endpoints within 5 kbp of permutated duplication positions. Thus, the edges of long contigs correspond to segmental duplications more often than expected by chance, indicating the continued challenge of accurately constructing assemblies that encompass duplicated sequence.

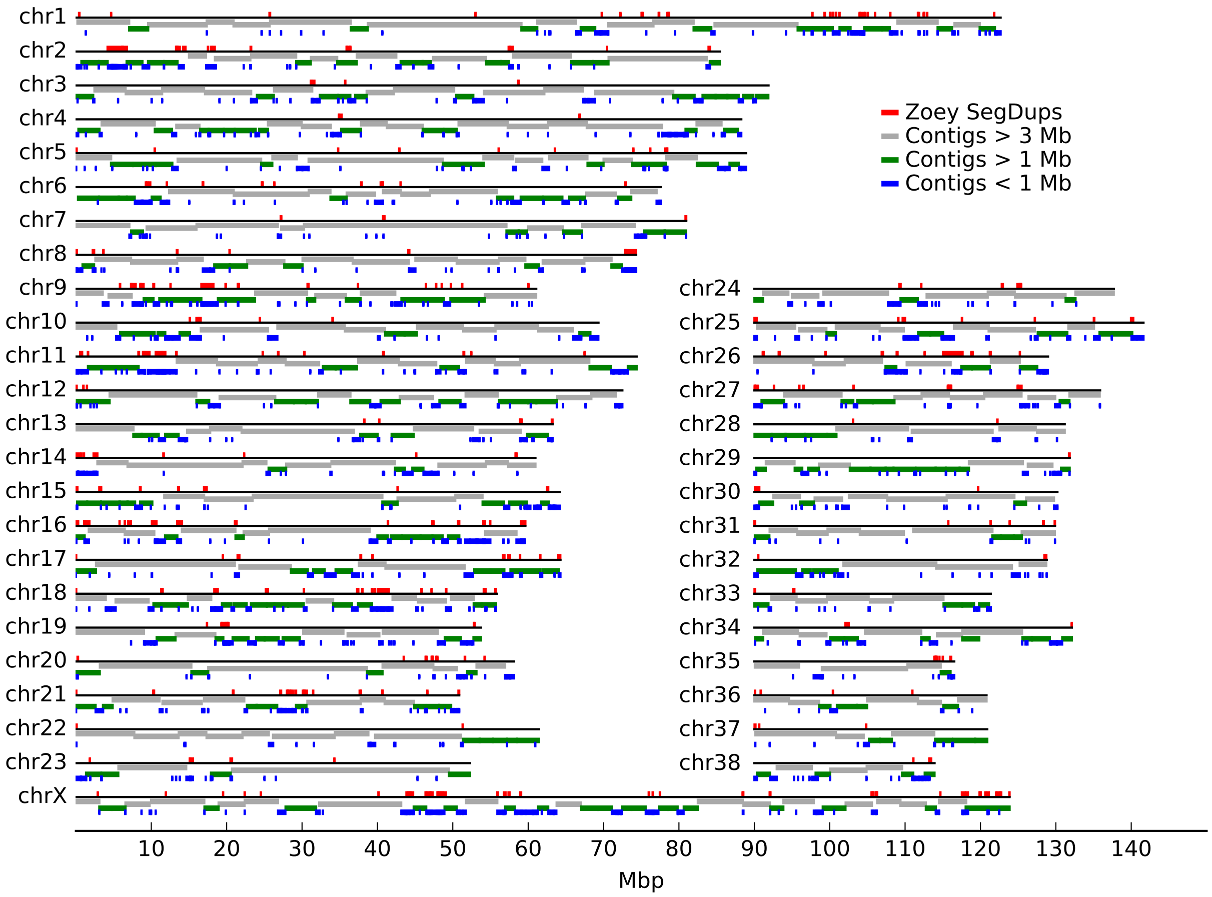


**Figure 1.4: Placement of assembled contigs on CanFam3.1 reference assembly.** The position of each contig is plotted on the CanFam3.1 genome. The colored bars below each line indicate the corresponding position of each contig, colored based on their length as indicated in the color key at the top right of the figure. Above each line, regions of segmental duplications based on read depth in the Zoey Illumina data are indicated by red boxes.

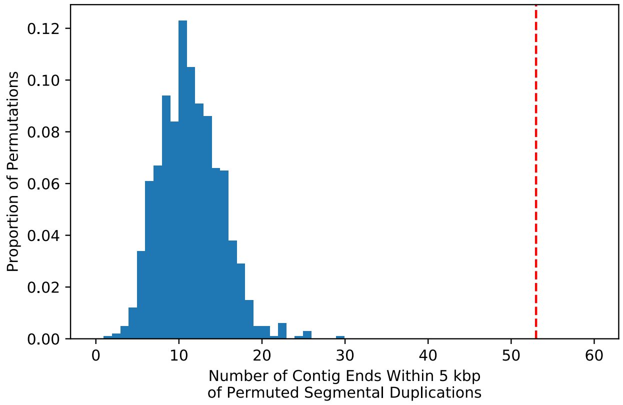


**Figure 1.5: Large contigs end at duplications more often than expected by chance.** We randomly permuted the position of segmental duplications identified by Zoey read depth and determined how many times a large contig (>3 Mb) ended within 5 kbp of a permuted position. The distribution of counts obtained from 1,000 random permutations is plotted. The observed value, of 53 contig ends within 5 kbp of a duplication, is indicated by the red dashed line.

We noted many genomic locations where regions not covered by the primary contigs were nonetheless spanned by raw PacBio reads. We reasoned that these reads may have been excluded from the assembly due to their length. Therefore, we performed a gap filling local assembly. For each region, raw PacBio reads were extracted and assembled using Canu v1.3 (21). The resultant contigs were compared with the flanking primary contigs using BLAT and retained if they showed >90% identity to flanking contigs. This process resulted in 373 candidate gap-filling contigs.

Next, we attempted to link the 2,640 filtered primary contigs along with the 373 candidate gap-filling contigs into scaffolds. This procedure was performed based on mapping of the Zoey mate pair data, Tasha BAC end sequence data, and Tasha fosmid data, using the BESST scaffolding algorithm (version 2.2.7) (22). Only 95 of the gap-filling contigs were linked to a primary contig by BESST, the remaining unlinked gap-filling contigs were omitted from subsequent analysis. This procedure resulted in a total of 1,759 scaffolds with a $N_{50}$ of 21 Mbp. The resulting scaffolds were aligned to the CanFam3.1 reference assembly using BLAT and MUMMER and organized into a chromosomal layout based on the mapping data (Figure 1.6). Additional analysis identified 691 contigs that were not linked to any other sequence using BESST, aligned to another scaffold with high sequence identity (>99% identity and > 15% fraction aligned), and did not correspond to a predicted duplication (copy number < 3 based on depth calculated using mosdepth) (23). These sequences likely represent allelic variants that were retained as primary contigs; they were removed in order to avoid contaminating the final assembly with allelic sequence variants. Finally, we utilized Zoey Illumina WGS short-read sequences to improve the base pair accuracy of the assembly. Reads were aligned using bwa-MEM (24)and processed using Picard to mark PCR duplicates. Variants then were identified using the GATK HaplotypeCaller (25). The sequence in the Zoey assembly was corrected for sites with a MAPQ value of at least 20 and where Zoey appeared homozygous (>90% alternative allele frequency).

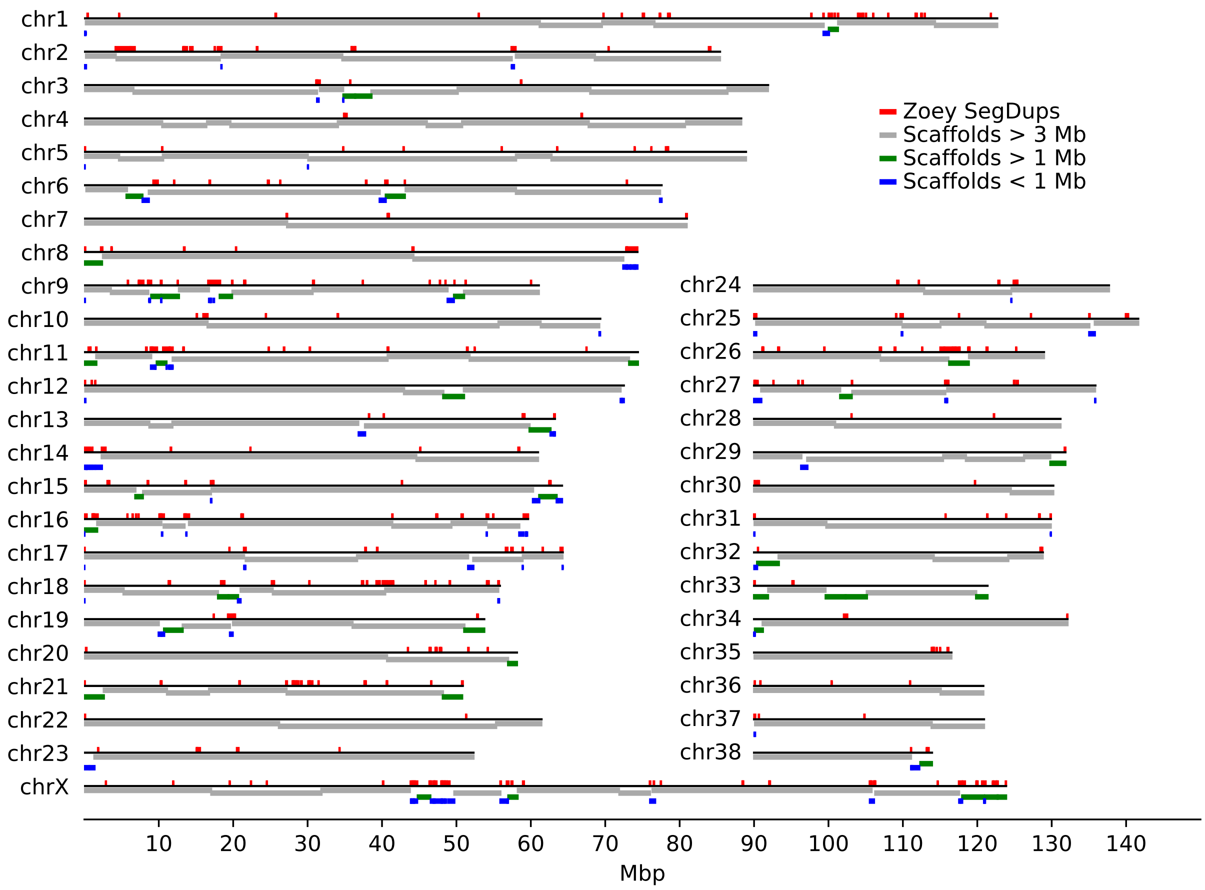


**Figure 1.6: Alignment of assembled scaffolds to CanFam3.1 reference assembly.** The position of each scaffold is plotted on the CanFam3.1 reference assembly. The colored bars below each line indicate the corresponding position of each scaffold. Scaffolds are colored based on their length as indicated in the color key at the top right of the figure. Above each line, regions of segmental duplications based on read depth in the Zoey Illumina data are indicated by red boxes.

### 2. Gene Annotation

*RNA-Seq read processing*

Forty-two canine RNA-Seq runs representing eleven tissue types were downloaded from the NCBI Sequence Read Archive (BioProject ID PRJNA78827) (26) (Table 2.1). Using trimmomatic (version 0.36) (27), we removed the first 10 bp of each read, trimmed any read at the point where a 4 bp sliding window did not achieve an average base quality score of at least 20, and eliminated any reads less than 50 bp in length. Additionally, rare k-mers were flagged and corrected using Rcorrector (28), and a script was used to remove reads that did not pass these filters (https://github.com/ harvardinformatics/TranscriptomeAssemblyTools/blob/master/ FilterUncorrectabledPEfastq.py). Read orientation information (e.g. /1 and /2) was appended to each read identifier using fastool (version 0.1.4; --append /1 or /2; https://github.com/fstrozzi/ Fastool/).

Reads from each library were aligned to both the Zoey and CanFam3.1 assemblies using GSNAP with a k-mer size of 15 (29). For both reference genomes, all autosomes, chromosome X, and unplaced sequences were treated as individual chromosomes. For alignment, we required that no read map to more than one genomic position (-n = 1). Alignments reported to extend past the end of chromosomes were corrected and the resulting alignment files were converted to sorted BAM format files using SAMtools (30). Using the alignment files, we assembled transcripts using Cufflinks (v2.2.1) (31, 32). Output files were merged either by tissue type or across all tissues using Cuffmerge (v2.2.1).

*Trinity de novo gene models*

Trinity can be run with and without a genome reference sequence as a guide. The cleaned and processed FASTQ files from the 42 RNA-Seq libraries were merged into left and right read files. Using Trinity (v2.3.2) (33, 34), these merged read files then were used as inputs for both genome guided and non-genome guided transcriptome assembly using a minimum contig length of 150bp (min_contig_length) and a normalized max read of 50 (normalize_max_read_cov). For the genome guided analysis, the sorted bam containing data merged across all tissue alignment files was used, with the maximum intron length (genome_guided_max_intron) set to 100,000 bp. A script provided with the Trinity package (TrinityStats.pl) calculated transcript count, length, and composition statistics (Table 2.2).

*PASA annotation pipeline*

We annotated the transcripts using PASA-Lite (https://github.com/PASApipeline/ PASA_Lite), which is a SQL-independent annotation pipeline that is built off of the original PASA annotation pipeline (35). Three inputs were used for this step: (i) the Trinity *de novo* (no genome) transcripts; (ii) the Trinity genome-guided transcripts based off of the Zoey transcript alignment data; and (iii) the merged GTF file generated by Cufflinks and Cuffmerge of the 42 RNA-Seq libraries aligned to the Zoey2.3 assembly. We first generated a merged Trinity transcript file by concatenating the fasta sequences from the *de novo* and genome-guided Trinity transcripts, yielding a total of 1,790,748 sequences (note, some could be redundant transcripts). We next used the Seqclean utility (http://www.tigr.org/tdb/tgi/software) to pre-process these merged transcripts to remove or trim low quality, vector, and poly-A sequences.

| | Reference Free | | Genome Guided | |
|---|---|---|---|---|
| | All Transcripts | Longest isoform per 'gene' | All Transcripts | Longest isoform per 'gene' |
| Total 'genes' | 746,294 | -- | 792,100 | -- |
| Total transcripts | 886,898 | -- | 903,850 | -- |
| Percent GC | 47.8 | -- | 46.83 | -- |
| Contig $N_{10}$ | 7,763 | 5,171 | 8,075 | 5,113 |
| Contig $N_{20}$ | 5,718 | 2,748 | 5,807 | 2,544 |
| Contig $N_{30}$ | 4,362 | 1,325 | 4,234 | 1,215 |
| Contig $N_{40}$ | 3,266 | 759 | 2,978 | 722 |
| Contig $N_{50}$ | 2,258 | 505 | 1,799 | 493 |
| Median contig length (bp) | 258 | 233 | 252 | 235 |
| Average contig length (bp) | 716.2 | 412.65 | 638.38 | 408.36 |
| Total assembled bases (bp) | 635,200,674 | 307,959,676 | 577,002,309 | 323,458,884 |

**Table 2.2 Statistics for transcripts assembled using Trinity.** Length and count statistics are shown for both the reference free and genome guided annotation pipelines. Results are shown using all transcripts as well as only considering the longest transcript present at each locus.

The next steps are based on alignment of the cleaned transcripts to the reference genome. First, we used BLAT (19) to align each transcript to the Zoey assembly. Second, alignments were processed using a script provided with the PASA software (process_BLAT_alignments.pl), which generated a GFF file based on the BLAT results with a maximum intron length of 500,000 bp (-I) and a maximum number of top hits equal to 10 (-N). Third, the alignments were processed using the PASA Alignment Validator script (PASA.AlignmentValidator). This script identifies transcript alignments that have a high percent identity across a large proportion of the transcript length, as well as the presence of minor or major consensus splice sites. This step also requires the GTF file generated by Cufflinks. For this step we required a minimum percent identity threshold of 95% (--min_per_id 95) with a 90% transcript length alignment threshold (default parameter). The outputs from the Alignment Validator script are two GTF files, one for valid and the other for invalid transcript alignments. Finally, in the fourth step, the Alignment Assembler (PASA.AlignmentAssembler) script was executed. This step identifies the best alignments, groups isoforms that likely result from alternative splicing, and performs open reading frame (ORF) identification. We ran the Alignment Assembler with all default parameters except the minimum threshold for the percent of shared sequence among isoforms, which was set to 20 (--min_pct_iso_shared_sequence 20). This last step generates a final PASA GTF file that contains passing transcripts and their possible isoforms.

*Transdecoder*

We used the transdecoder pipeline (version 5.0.1) (34) to annotate putative coding structure within the transcripts processed using PASA. First, the GTF file from PASA is converted to transcript FASTA sequence using the transdecoder script: cufflinks_gtf_genome_to_cdna_fasta.pl. A GFF3 file is generated using the script cufflinks_gtf_to_alignment_gff3.pl, through conversion of the same PASA GTF file. Open reading frames (ORFs) are annotated based on the newly generated transcript sequence files using the script: TransDecoder.LongOrfs. Finally, GFF3 and BED files from the annotated transcripts are generated in Zoey assembly coordinates using the scripts: cdna_alignment_orf_to_genome_orf.pl and gff3_file_to_bed.pl. In total, this resulted in a set of 1,301,960 transcript models.

*Transcript redundancy and filtering*

Examination indicated that the transdecoder output contained a considerable level of redundancy among transcript models as well as the presence of single exon gene models that overlapped fully with annotated repetitive elements. We therefore performed a series of additional filtration steps. First, for each locus, we selected the transcript with the highest score provided by transdecoder as the representative gene model. Second, we intersected single exon gene models with the genome RepeatMasker tracks annotated in the Zoey assembly and removed all single exon gene models that had greater than 10% of their total length annotated as LINE, SINE, or LTR retroelements. Third, we observed the presence of many nearly identical single exon gene models at a given locus. Therefore, we performed a reciprocal BLAT search and any single exon gene model that was at least 95% encompassed by another larger gene was removed from the analysis. These filters resulted in a set of 42,911 gene models, including 23,703 single exon models and 19,208 multi-exon gene models. Examination indicated that this set still included many likely spurious single-exon models which partially overlapped with an exon from a multi-gene model (Figure 2.1). Thus, we performed a final filtration step that included only those single exon models with a match to a previously described protein (using BLAST2GO, see below) and that did not intersect with a multi-exon model. This final set of protein-coding genes included a total of 22,182 gene models (2,974 single exon models and 19,208 multi-exon models, see Table 2.3).

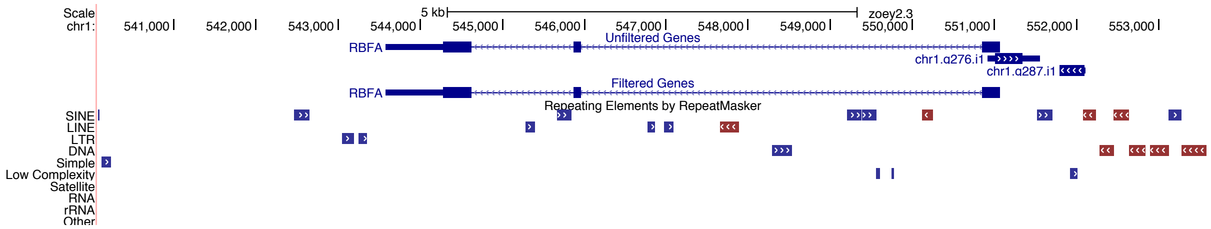


**Figure 2.1: Gene model filtration: A UCSC TrackHub view is shown to illustrate the gene filtration steps.** The top track contains 42,911 genes models prior to filtration and includes a gene annotated as *RBFA* as well as two single exon gene models. These single exon models are removed following filtration (Filtered Genes track). The RepeatMasker annotation for this region is shown at the bottom of the figure. The orientation of the identified repeat elements is indicated by color: blue (forward) and red (reverse).

*Functional annotation of predicted gene models*

We utilized homology-based searches to determine the predicted functional role and possible gene symbol for each gene model. The peptide fasta sequences, provided by transdecoder, for the final 22,182 genes were searched against the non-redundant protein database (blastp -db nr -outfmt 5 -evalue 1e-3 -word_size 3 -show_gis -max_hsps_per_subject 20 -num_threads 5 -max_target_seqs 20). The resultant XML output files were loaded into BLAST2GO (36), with the following parameters: minimum annotation cut-off of 55, GO weight equal to 5, BLASTp cut-off equal to $1e^{-6}$, HSP-hit cut-off of 0, and a hit filter equal to 55 (see Table 2.3).

*Alignment of gene models to CanFam3.1*

We aligned the gene models against CanFam3.1 reference assembly using BLAT and found that 18,834 of the 22,182 final genes (84.9%) had an alignment to CanFam3.1 that spanned the entire gene model length. An additional 1,836 (8.3%) of the gene models had an alignment that spanned more than 95% of the gene model. Of the 1,512 (6.8%) of gene models with an alignment that spanned less than 95% of the gene model length, 356 (1.6%) had hits that spanned less than 75% of the gene model length. This includes 49 (0.2%) gene models that had no hits against the CanFam3.1 assembly. This information is given in Table 2.3. We note that despite the filters described above, many suspect gene models remain. In fact, 310 of the final gene models have at least 50% of their length annotated by RepeatMasker. Thus, we performed an additional analysis that removed any BLAT hit that had a 50% overlap with segments of the gene model annotated by RepeatMasker. This process removed all of the hits for 286 of the genes and identified 15,853 with a remaining alignment that spanned at least 75% of their length.

*Quantification of gene expression*

We estimated expression levels for each of the 22,182 protein-coding gene models using Kallisto (version 0.46.0) (37). Gene transcript fasta sequences were indexed with 31 bp kmers and quantified using the –bias option. Of the 22,182, genes, 97% (21,497) have an estimated expression level greater than one transcript per million (TPM) in at least one of the 42 SRA read datasets. Next, the expression level was estimated for each of the 11 tissues (including one tissue of unknown origin) using all of the reads from each tissue (Figure 2.2). This analysis identified 988 transcripts that do not have an estimated expression level >1 TPM in any of the tissues and 1,599 transcripts above that threshold in only a single tissue. We additionally find that 2,005 transcripts models have an expression level >10 TPM in each of the 11 analyzed tissues. We created a TrackHub barChart track that shows the expression level of each gene (Figure 2.3).

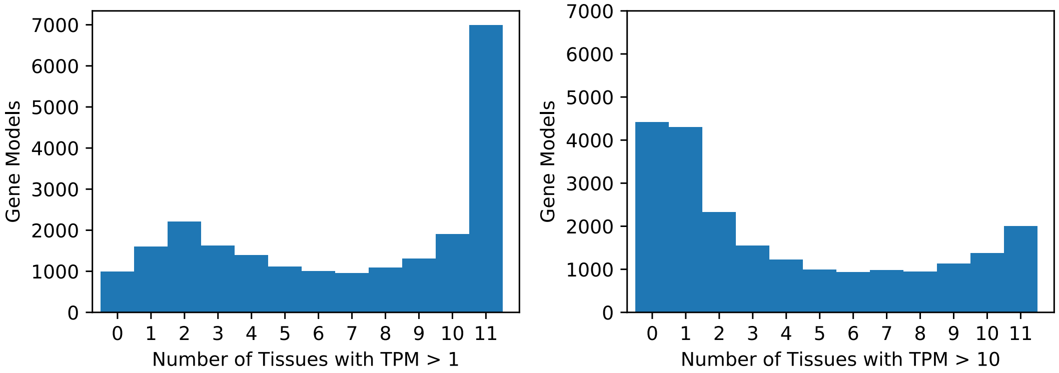


**Figure 2.2: Gene abundance across tissues.** Histograms are shown for the abundance of 22,182 protein-coding genes across 11 tissues (including one tissue of unknown origin). Distributions are shown using a cutoff of 1 TPM and 10 TPM.

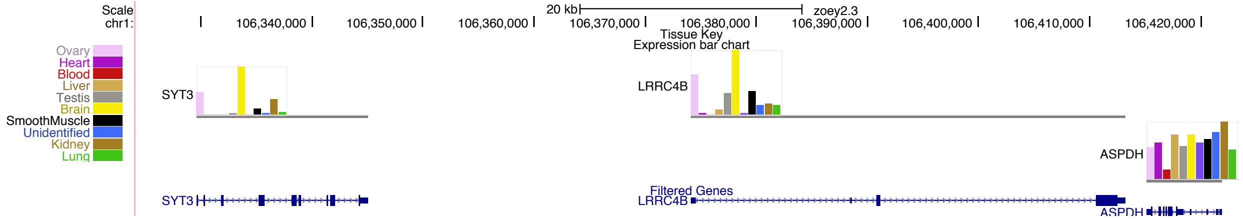


**Figure 2.3: Gene expression levels across tissues.** The estimated expression level (in units of $log_{10}$(TPM)) is shown for 11 tissues for each gene. Each bar corresponds to a different tissue, as indicated by the color key in the left of the figure.

*Long non-coding RNA analysis*

We annotated long non-coding RNAs in the Zoey genome using the FEELnc program (38). Analysis began with the set of transcripts assembled using the Trinity and PASA-Lite pipelines described above. Multi-exon transcripts > 200 nt in length that did not overlap with protein-coding loci were extracted and 7,888 of these were consistently annotated as lncRNAs using FEELnc. These 7,888 candidate lncRNAs were aligned to the CanFam3.1 assembly using minimap2 and BLAT. We observed that 749 candidate lncRNAs had a partial overlap with an mRNA transcript annotated by Ensembl. These transcripts were removed from the analysis, resulting in a final set of 7,049 lncRNAs. Of these, 18 could not be mapped to CanFam3.1 while 66 had partial alignments with low mapping identity (<95%). A total of 7,055 transcripts had a full CanFam3.1 alignment with high identity (>95%). Of these, only 1,591 overlapped with an existing annotated non-coding RNA.

### 3. Assembly Annotation

*Repetitive sequences*

Common repeats in both the CanFam3.1 and Zoey assemblies were masked using RepeatMasker (39) version 4.0.7 with option '–species dog' using the rmblastn (version 2.2.27+) search engine and a combined repeat database of Dfam_Consensus-20170127 and RepBase-20170127. In both assemblies, 40.5% of the genome was annotated as a repeat by RepeatMasker. An additional search against both assemblies was made using only the Repbase L1_Cf consensus sequence as a custom library. Tandem repeats were identified using the program trf (version 4.04) (40) with options '2 7 7 80 10 50 500'. Windowmasker/DUST annotations (41) were created using the NCBI-blast toolkit version 2.5.0. Tracks corresponding to these annotations are available in the Zoey assembly track hub.

*Segmental duplications*

Segmental duplications were identified using two approaches: an assembly self-alignment and read-depth analysis. Self-alignment analysis of each assembly was performed using SEDEF (42) with default parameters. Results were filtered for alignments at least 1 kbp in length and at least 90% sequence identity (Table 3.1). After merging redundant alignments, a total of 57,568,649 bp on the Zoey assembly and 85,880,296 on the CanFam3.1 assembly (including the unplaced contigs) were identified as being in segmental duplications based on genome self-alignment.

Read-depth analysis was performed using fastCN as described previously (12). Reads from three samples were analyzed independently against both the CanFam3.1 and Zoey assemblies, which include (i) Illumina WGS short-read data from Zoey; (ii) Illumina WGS short-read data from the Iberian Wolf, Penelope; and (iii) Sanger reads from the 4 kbp plasmid library generated from the boxer, Tasha. As part of the fastCN pipeline, reads were divided into non-overlapping 36 bp segments. Copy-number estimates were constructed in non-overlapping windows that each contain 3 kbp of unmasked sequence. Segmental duplications were identified as runs of four windows in a row where each window has an estimated copy-number >=2.5. Considering only the primary chromosomes, alignment of the Zoey read data to the Zoey assembly identified a total of 467 segments with a total of 40,077,510 bp. Alignment of the Tasha plasmid data to the CanFam3.1 assembly identified 1,842 segments and a total of 73,421,120 bp. The plasmid data has a lower coverage than the Illumina short-read data sets, resulting in noisier copy-number estimates. To limit the confounding effects of differences in the source data, we additionally estimated duplications using read-depth information from Illumina snort-read data derived from the Iberian Wolf, Penelope. On the primary chromosomes of the CanFam3.1 assembly, the wolf data identified 459 segments with a total size of 47,757,534 bp, whereas the wolf data aligned to Zoey identified 468 segments with a total size of 40,836,807 bp.

In the Zoey genome, the union of the assembly comparison and Zoey read-depth approaches identified 2.7% of the primary chromosomes as segmentally duplicated. Individual chromosomes showed a wide range of duplication content, ranging from 0.9% for chr22 to 7.8% for chr26 (Table 3.2).

*QuicK-mer2 analysis*

An index for the Zoey assembly was constructed using QuicK-mer2 with default arguments (20), identifying a total of 2,050,414,720 30-mers for analysis. We supplemented the control regions used for the fastCN analysis by also excluding segmental duplications defined by assembly self-alignment and Zoey read depth.

*Alignment of Tasha clone end-sequences*

Clone end sequences from Tasha were aligned against the Zoey and CanFam3.1 genomes using bwa mem (24). Read-pair signatures identify clones that may span structural variants between the sample genome and the reference assembly. This analysis is based on the distance between the mapped position of the read pairs and the orientation of the aligned reads (concordant placements have reads on opposite ends aligned to different strands of the same chromosome in an inward orientation). Concordant pairs were defined as having proper read orientation and an apparent insert size within five median absolute deviations from the observed median size. For the CHORI-82 BAC library, 142,006 BACs showed a concordant alignment to CanFam3.1 while 141,367 had a concordant alignment to the Zoey assembly. Results were converted into bed format for display as tracks in the UCSC assembly hub browser. Scripts for processing the alignments of clone end-sequence reads are available at: https://github.com/jmkidd/sanger_dl.

*liftOver track generation*

We created chain files for use with the UCSC liftOver tool based on BLAT alignments. The procedure was based on the UCSC documentation available at: http://genomewiki.ucsc.edu/index.php Minimal_Steps_For_LiftOver. Chain files were generated in both directions, allowing for coordinate conversion from CanFam3.1 to Zoey and from Zoey to CanFam3.1 coordinates.

We created a track to visually illustrate the comparison between the two genomes. Using each assembly in turn as a reference, we created a bed file of overlapping 1 kbp windows with a step size of 250 bp. The window coordinates were converted to the other assembly using liftOver, and windows with consistent overlapping placements on both assemblies were merged together. Going from CanFam3.1 to Zoey, 94.91% of windows had a successful liftOver conversion. When going from Zoey to CanFam3.1, 96.97% of windows had a successful liftOver conversion.

A BED formatted track for display was created from the resultant merged window coordinates. Segments that were assigned to the same chromosome on the other assembly are colored in black, segments with a lift over in an inverted orientation are colored in green, and segments that correspond to a different chromosome are colored in purple. Regions without a corresponding segment on the other assembly are not included, resulting in a gap in the displayed track.

*Inversion analysis*

We mined the liftOver tracks to identify candidate inversions between the two assemblies. Because some calls appeared to be fragmented due to individual windows that did not liftOver between assemblies, candidates within 10kb were merged together. Candidate intervals were required to be at least 2.5 kbp in length, have less than 75% of their bases annotated by RepeatMasker (39), and less than 50% of their length annotated as a segmental duplication. Sequence interval coordinates were refined by mapping the corresponding sequence using minimap2 (43). This process resulted in 44 candidate inversions between the two assemblies (Table 3.3). The majority of these candidate inversions (30/44) have segmental duplications present in at least one of their breakpoints. Non-allelic homologous recombination

(NAHR) among inverted duplications may explain the origin of these inversions. However, assemblies across duplicated regions may not be accurate; thus, the candidate inversions we describe should be interpreted with caution.

Although the X chromosome only accounts for 5.3% of the Zoey assembly length, 41% of the candidate inversions (18/44) are located on the X chromosome. The X chromosome has a unique relationship with inversions. Due to the presence of only a single homolog in the heterogametic sex, the sex chromosomes show a reduced recombination rate. This reduced recombination rate has been associated with chromosomal inversions (reviewed in (44)). In some species, the X chromosome is also enriched for high-identity segmental duplications known as amplicons (45). These amplicons typically contain genes that are expressed in the testis and may be linked to complex evolutionary forces related to fertility (45). Large segmental duplications may be a substrate for inversions through NAHR. A region on the X has been found to be recurrently inverted across eutherians (46) and an enrichment of inversions on the X chromosome has also been reported in Great Apes (47).

We note that our chromosomal layout relied upon the CanFam3.1 assembly; thus we did not correct likely orientation errors that have been suggested for chromosomes 27 and 32 (48, 49).

*Identification of single nucleotide variants in Zoey*

Illumina reads from Zoey were aligned to CanFam3.1 using bwa mem (24) and processed to perform indel realignment, base quality score recalibration, and marking PCR duplicates using Picard (http://broadinstitute.github.io/picard/) and GATK v3.7. SNP calls were generated using the haplotype caller algorithm in GATK v3.7 (25). The VQSR procedure was used to select a high-quality set of variants using the sites on canine Illumina SNP array as a training set. We set cutoffs such that 99% of the sites on the Illumina array were retained, resulting in a filtered call set of 3,573,985 single nucleotide variants (SNVs). Zoey is predicted to be heterozygous at 1,928,497 of these positions.

### 4. Analysis of CanFam3.1 Sequence Gaps

*CanFam3.1 gaps resolved in Zoey*

The CanFam3.1 genome assembly contains a total of 23,876 gaps, of which 19,553 are located on a primary chromosome sequence (autosomes + chrX). Because the Zoey assembly only contains 997 gaps, we expect that the majority of the gaps in CanFam3.1 are filled. Using liftOver, we determined the approximate coordinates in the Zoey assembly that correspond to each of the chromosomal CanFam3.1 gaps. These results were further refined by extracting the corresponding sequence from the Zoey assembly, plus 5 kbp of flanking sequence, and aligning back to CanFam3.1 using BLAT (19). This analysis resulted in a confident, unique alignment match with base-pair resolution for 12,806 gaps in the CanFam3.1 assembly (Table 4.1, Figure 4.1). There is a strong correlation (Pearson R = 0.414) between the size of the sequence in the Zoey assembly and the corresponding gap segment in CanFam3.1 (Figure 4.2).

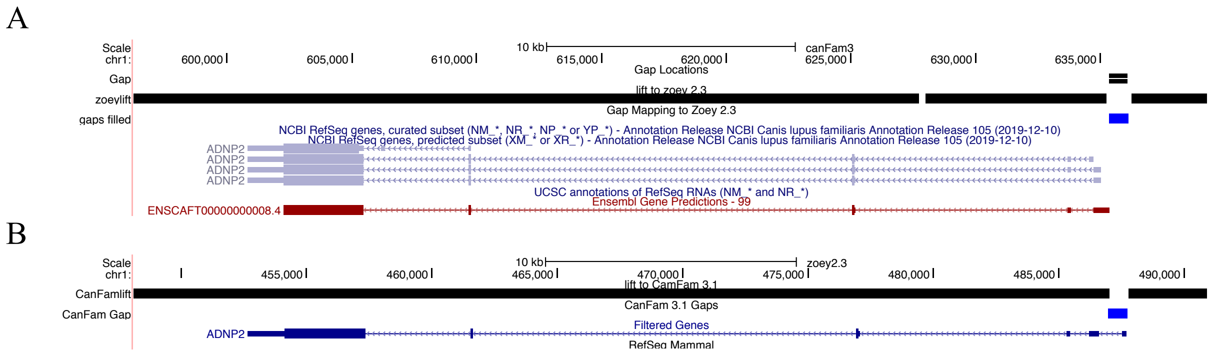


**Figure 4.1: Resolving gaps in the CanFam3.1 assembly.** Genome browser plots are shown for a corresponding region in the CanFam3.1 (A, chr1:596255-639326) and Zoey (B, chr1:448123-490934) genome assemblies. In the CanFam3.1 genome, an assembly gap is present adjacent to the *ADNP2* gene model. This assembly gap is filled in the Zoey genome (indicated by blue box). The corresponding view in the Zoey genome (B) shows that the first exon of the predicted *ADNP2* gene model resides within the sequence that is absent from CanFam3.1. In each plot the solid black track at the top indicates contiguous portions of the genome that liftOver to the same chromosome between the assemblies.

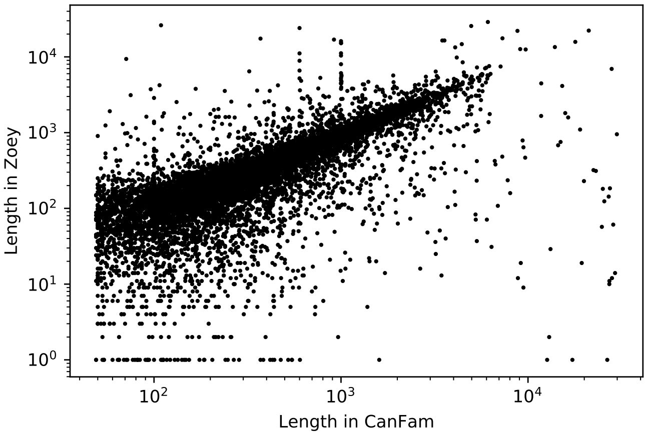


**Figure 4.2: Comparisons of the gap sequence length.** Depicted are the sizes of sequences corresponding to 12,806 gaps in the CanFam3.1 assembly. The X axis gives the size of the gap in CamFam3.1 while the Y axis shows the size of the corresponding segment in the Zoey assembly. Note that each axis is displayed on a logarithmic scale.

*Copy-number analysis of sequence in CanFam3.1 assembly gaps*

We estimated genomic copy-number across 55 canine samples (Table 1.1) based on the Zoey assembly using QuicK-mer2 (20). QuicK-mer2 utilizes a k-mer matching strategy to rapidly estimate genomic copy-number in a paralog-specific manner from Illumina short-read sequencing data sets. This estimate is achieved by tabulating the observed read depth at defined sets of k-mers (k=30 bp) that are unique in the genome reference assembly. Preliminary analysis identified regions of the Zoey assembly that were absent from many canines and that corresponded to regions that were gaps in CanFam3.1. We explored this pattern systematically. To avoid the confounding effect of sample sex on copy-number, we focused on 11,267 intervals in Zoey that: (i) correspond to CanFam3.1 gaps, (ii) are present on the autosomes, and (iii) contain at least 10 unique k-mers. We found that 50% of the intervals (5,739 of 11,267) had an estimated copy-number of 0 (raw value less than 0.5) in at least 28 of the 55 samples. If true, these data would suggest that nearly half of the sequences found in gaps in the CanFam3.1 assembly are absent across most other individual dogs. We further looked into the patterns of presence and absence and found a clear peak corresponding to sequences that are absent in 46 of the 55 analyzed canines (Figure 4.3). Examination of sample level data showed that a substantial fraction of these sequences appears to be present in only the same nine samples (Figure 4.4).

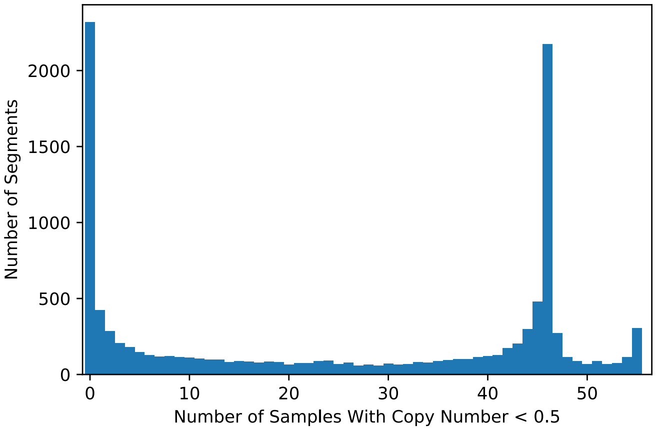


**Figure 4.3: CanFam3.1 Gap Sequences which are absent from a majority of analyzed samples.** Depicted is a histogram showing the number of samples for each segment that have a copy-number estimated by Quic-Kmer2 that is less than 0.5.

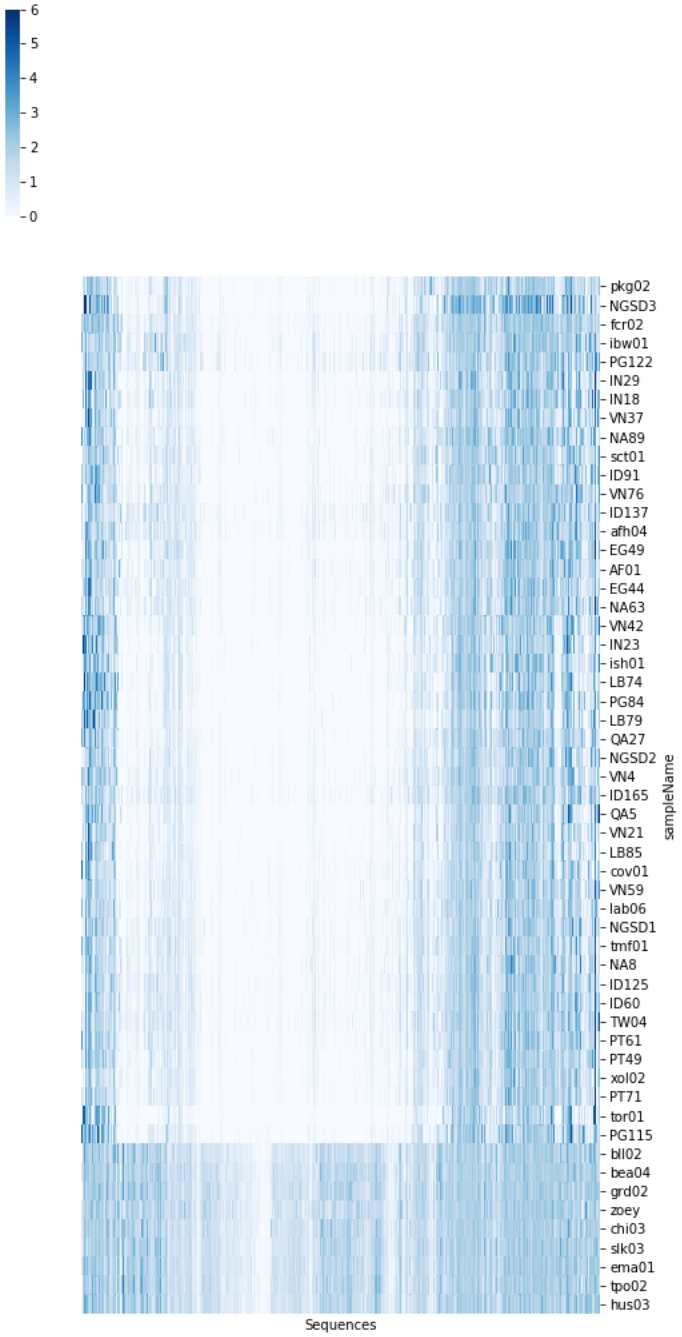


**Figure 4.4: Estimated copy-number of gap sequences across dog samples.** The heatmap depicts the estimated copy-number for sequences corresponding to autosomal CanFam3.1 sequence gaps. Copy number is estimated using QuicK-mer2 and is indicated by the color bar in the top left. The X axis represents the individual gap-sequences. The examined canine samples are shown along the Y axis. The order of the rows and columns was determined based on median Euclidean distance clustering.

The QuicK-mer2 data suggest that sequences corresponding to CanFam3.1 sequence gaps are absent across a collection of sequenced canines. To explore this result further, we interrogated a subset of these sequences using a custom Agilent array comparative genomic hybridization (array-CGH) platform. This array included probes designed against a preliminary set of assembled canine sequences that are absent from the CanFam3.1 reference genome. The array contained at least one CGH probe for 1,735 of the 11,267 autosomal regions analyzed by Quick-mer2. We analyzed the hybridization intensity data in two ways. First, we considered the distribution of single channel ('red' or test sample) intensities across the relevant probes and found that probes corresponding to CanFam3.1 gap sequences exhibited a hybridization intensity well above the background level (Figure 4.5). Next, we compared the QuicK-mer2 depth, single channel sample CGH intensity, and $\log_2$ intensity (test/Zoey) for the 1,735 segments. Although some samples appear to have a systematically increased or decreased intensity, the striking pattern of sample-specific absence indicated by the QuicK-mer2 analysis was not supported by the array CGH data (Figure 4.6 and Figure 4.7).

These copy number data represent contradictory information: analysis of Illumina short-read sequence data suggest that the sequences are often absent from canine genomes, while hybridization analysis indicates that the segments are in fact present in genomic DNA. To resolve this contradiction, we examined the individual aligned read data in greater detail. Deletions in a sample relative to a refence genome assembly manifest with distinct read alignment characteristics, including read-pairs that appear to map far apart and the presence of split or soft-clipped alignments across the deletion junction (50). Examination of Illumina reads aligned to the Zoey reference assembly failed to show any of the standard indicators of a deletion at these regions (Figure 4.8).

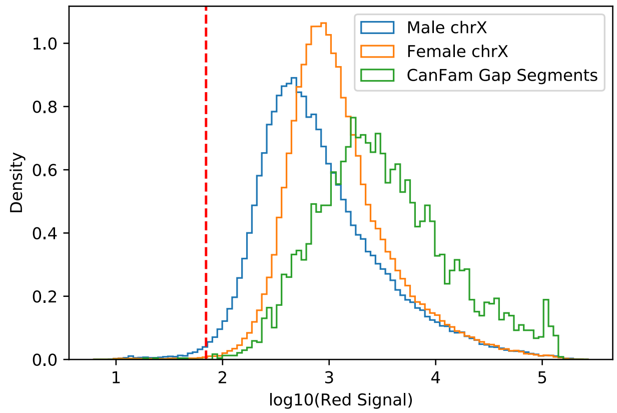


**Figure 4.5: Single channel probe intensity distribution.** A histogram of the single channel (red) intensity is shown across samples. The blue line gives the distribution of intensity values for all probes on the X chromosome in males, representing typical intensity for single copy sequence. The orange line shows the intensity distribution of X chromosome probes in female samples. The green line shows the intensity distribution for probes from the 1,753 autosomal gap segments across all samples. The vertical red line corresponds to the 1% tail of intensity values observed for X chromosome probes in males. The cutoff value is $\log_{10}(69.964)=1.84487$.

**A** **B**

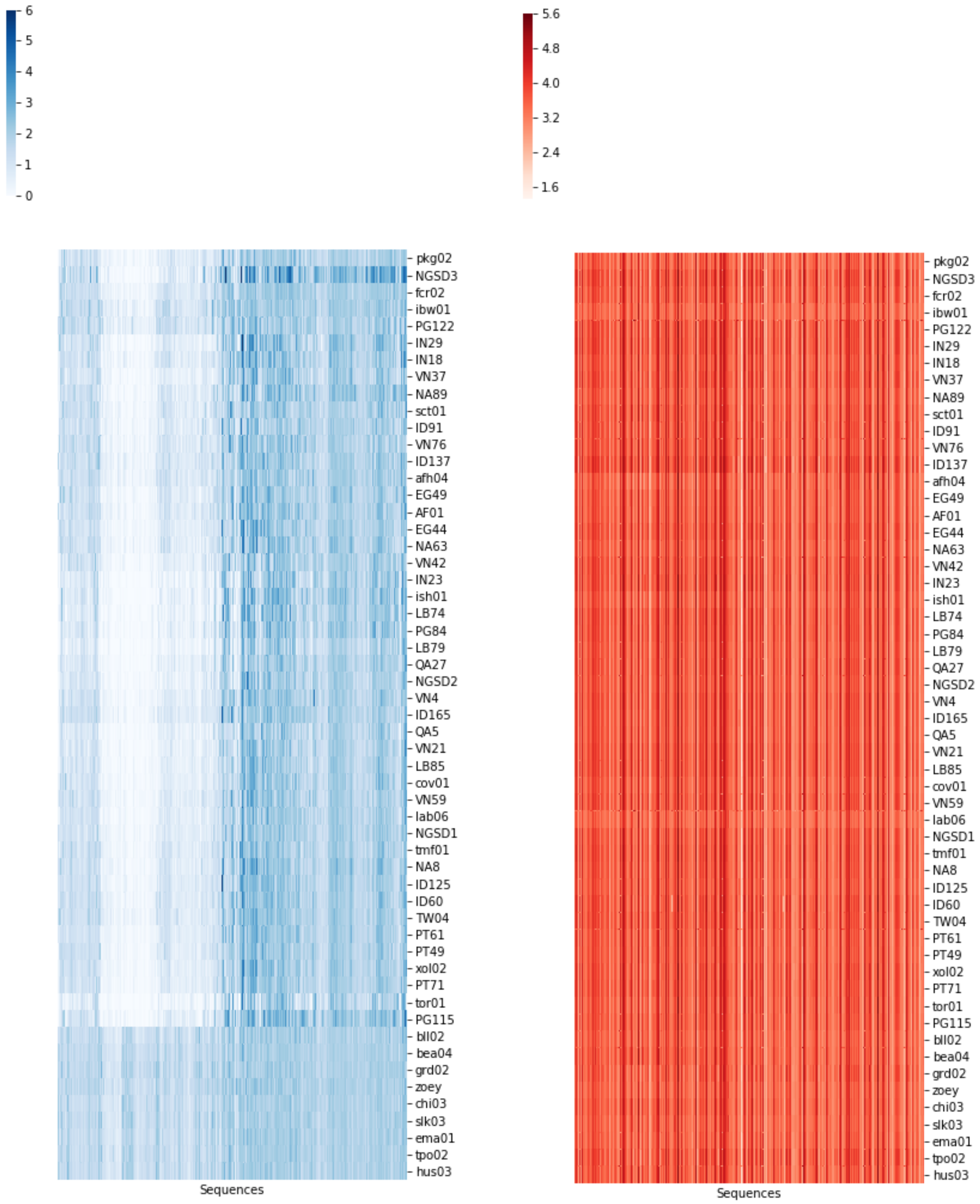


**Figure 4.6: QuicKmer-2 and single channel array CGH estimates of segment copy-number.** Each heatmap shows the estimated values for 1,753 CanFam3.1 gap segments estimated in 55 samples. (A) Estimated from QuicK-mer2 based on Illumina short-read data. The color bar in the top left indicates estimated copy-number. (B) Single channel sample intensity from array CGH. The color bar in the top left indicates $\log_{10}$ transformed intensity in the 'red' channel. The order of columns and rows is the same in both plots and was based on Figure 4.4.

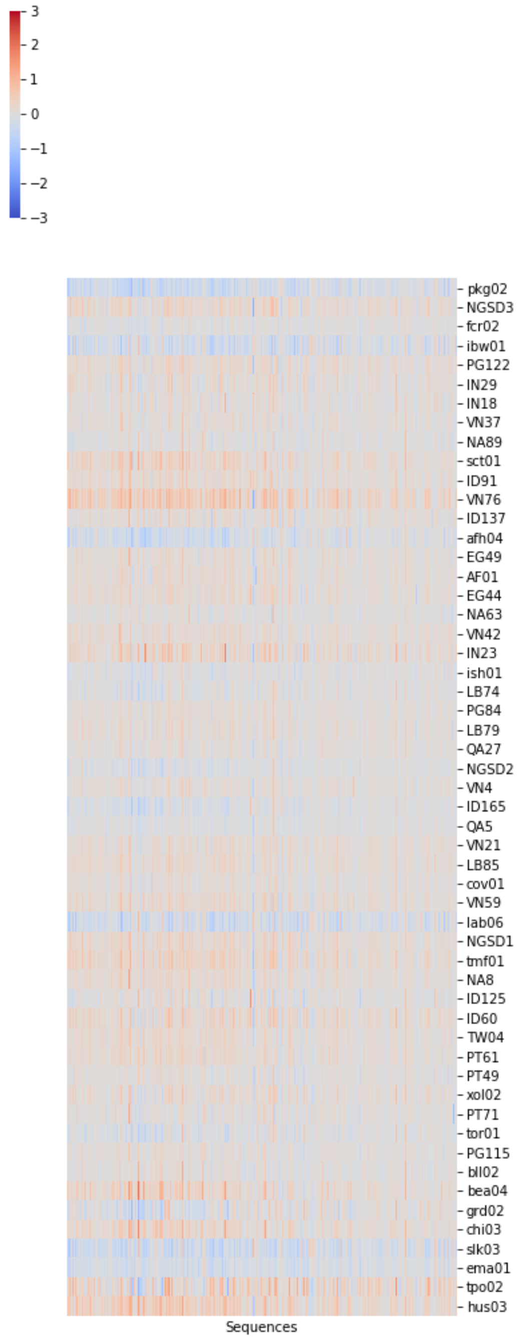


**Figure 4.7: Array-CGH relative intensity data.** The $\log_2$ ratio of intensity test *vs* reference intensity for each of 54 samples is shown for 1,735 gap segments. For each comparison, genomic DNA from Zoey was used as the reference. Row and column order are the same as in Figure 4.6. The color bar in the top left indicates the $\log_2$ ratio of (test/refence) signal.

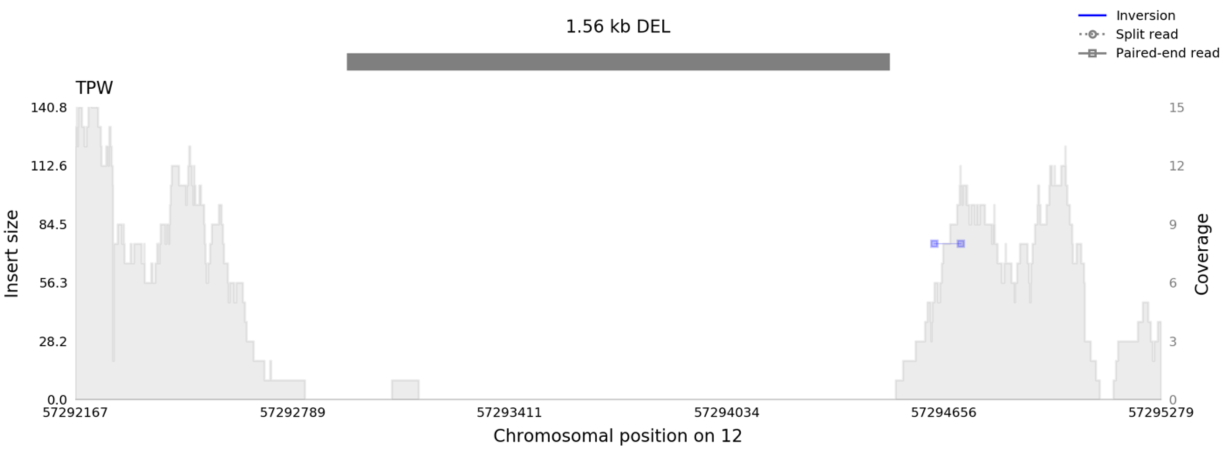


**Figure 4.8: Lack of sequence coverage without signals of deletion.** A view of reads from sample TPW (SRR1635701) aligned to the Zoey genome assembly is shown. The region corresponding to a gap is indicated as a grey bar as a candidate deletion. Although there is lack of read coverage, there are no corresponding split reads or aberrant read-pairs spanning across the interval. The plot was created using samplot (51).

*CanFam3.1 gaps are enriched for sequences with extreme GC content*

Together, the above observations indicate that although segments corresponding to gaps in the CanFam3.1 assembly are present in genomic DNA isolated from individual canines, reads derived from the regions tend to be missing from Illumina sequencing data sets without displaying read signatures typically associated with deletions. These data suggest the regions are not represented due to some technical bias in the Illumina sequencing data. Illumina sequencing is known to have a bias against sequences with extreme GC content, a bias that has been traced to multiple steps in the library preparation, flow cell cluster generation, and sequencing processes (52). Examining the gap sequences revealed an enrichment for high GC content relative to randomly permuted intervals of the same size (a median value of 67.3% GC when compared to an expected value of 39.6% GC). Examination of the distribution of GC content across segments indicates the presence of two classes of sequence: those with a GC content matching the genome wide average and a subset with increased GC content (Figure 4.9).

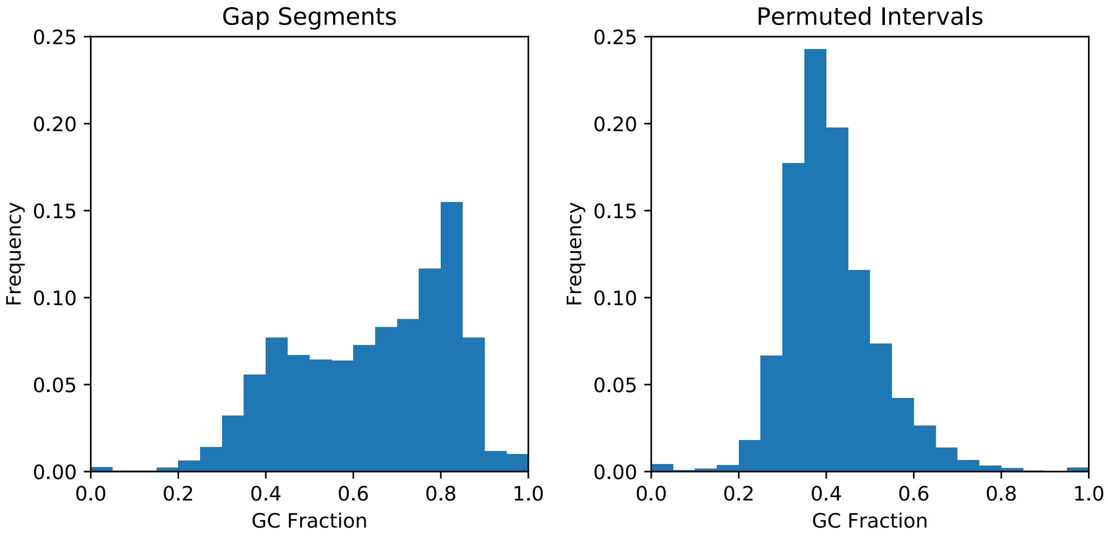


**Figure 4.9: GC content of sequences which are gaps in the CanFam3.1 assembly.** A histogram of the GC content is shown for 12,806 segments which correspond to gaps in the CanFam3.1 assembly (left). For comparison, the GC fraction was determined for a collection of randomly sampled intervals of the same size (right).

To investigate the above result further, for each segment we randomly sampled 1,000 locations of the same length in the Zoey genome and determined how many of the randomly chosen locations have a GC content as extreme as the original gap segment. This analysis yielded a permutation-based p-value indicating how extreme the GC content of each gap segment is relative to a random expectation (Figure 4.10). Of the 12,806 segments, 5,553 have a GC content greater than that found in 99% of permutations. These intervals have a median GC content of 80.9%, a median length of 531 bp, and encompass a total of 4.026 Mbp.

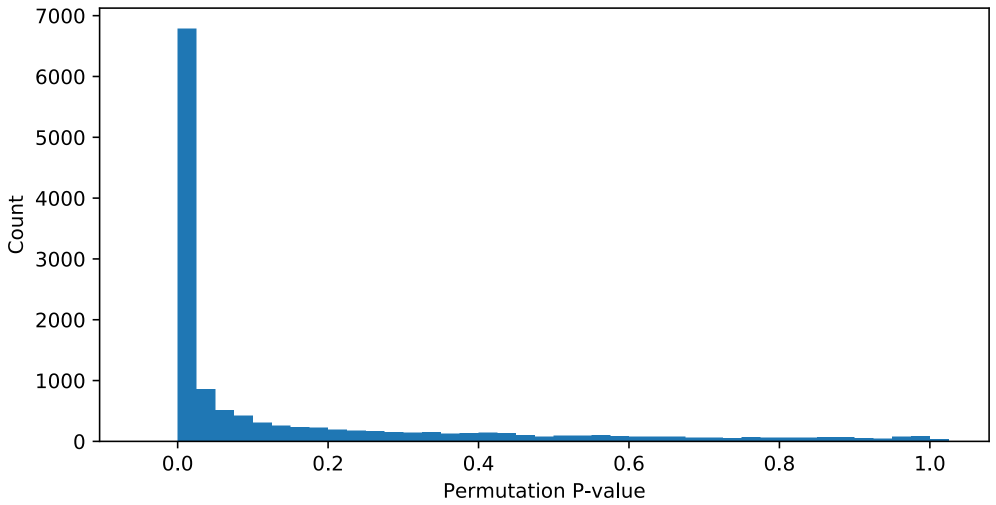


**Figure 4.10: Permutation-based p-value for GC content of each CanFam3.1 gap segment.** Results are based on 1,000 randomly permutated genome locations.

*Extremely GC-rich segments are located near transcription start sites and recombination hotspots*

Previous examinations of the CanFam3.1 assembly have indicated that the dog genome contains more CpG islands than other mammals (53, 54). Regions of high GC content in the dog genome have previously been associated with the 5' end of gene models (26). Regions of high GC content are also associated with recombination hotspots in dogs, a species that lacks a functional *PRDM9* gene, which specifies the location of recombination events in other vertebrates (55, 56). In dogs, recombination rates are substantially higher in regions with a high GC content near gene promoters (56). We determined the distance of each gap segment to the transcription start site of the nearest protein coding gene, based on the Zoey gene annotation. We then compared these distances with those obtained by randomly permutating the location of the gap segments and assigned each a permutation-based p-value based on how many randomly permutated segments of the same length were as close to a transcript start site. In total, 16.8% (2,151) of the gap segments overlap with a transcription start site. As a class, the 5,553 segments with an extreme GC content were located much closer to a transcription start site than expected by chance, with a median distance of 290 bp, compared with a median distance of 16,846 bp for the 7,253 gap segments not classified as having an extreme GC content. Permutation comparisons indicate that 21.5% (1,195) of the extreme GC intervals have a permutation-based p-value less than 0.01 and 61.2% (3,397) have a permutation p-value less than 0.5 (Figure 4.11). We performed a similar analysis based on distance to recombination hotspots identified from an LD-based genetic map (56). Due to limitations of the genetic map, regions on the X chromosome were excluded from the analysis. We found that 11.4% (1,402/12,304) of the extreme GC-rich regions are located within 1 kbp of a hotspot, compared to only 2.9% of permuted intervals that are within 1 kbp of a hotspot. As for the analysis of distance to transcription start sites, this enrichment was driven by the subset of gap segments with extreme GC content (Figure 4.12).

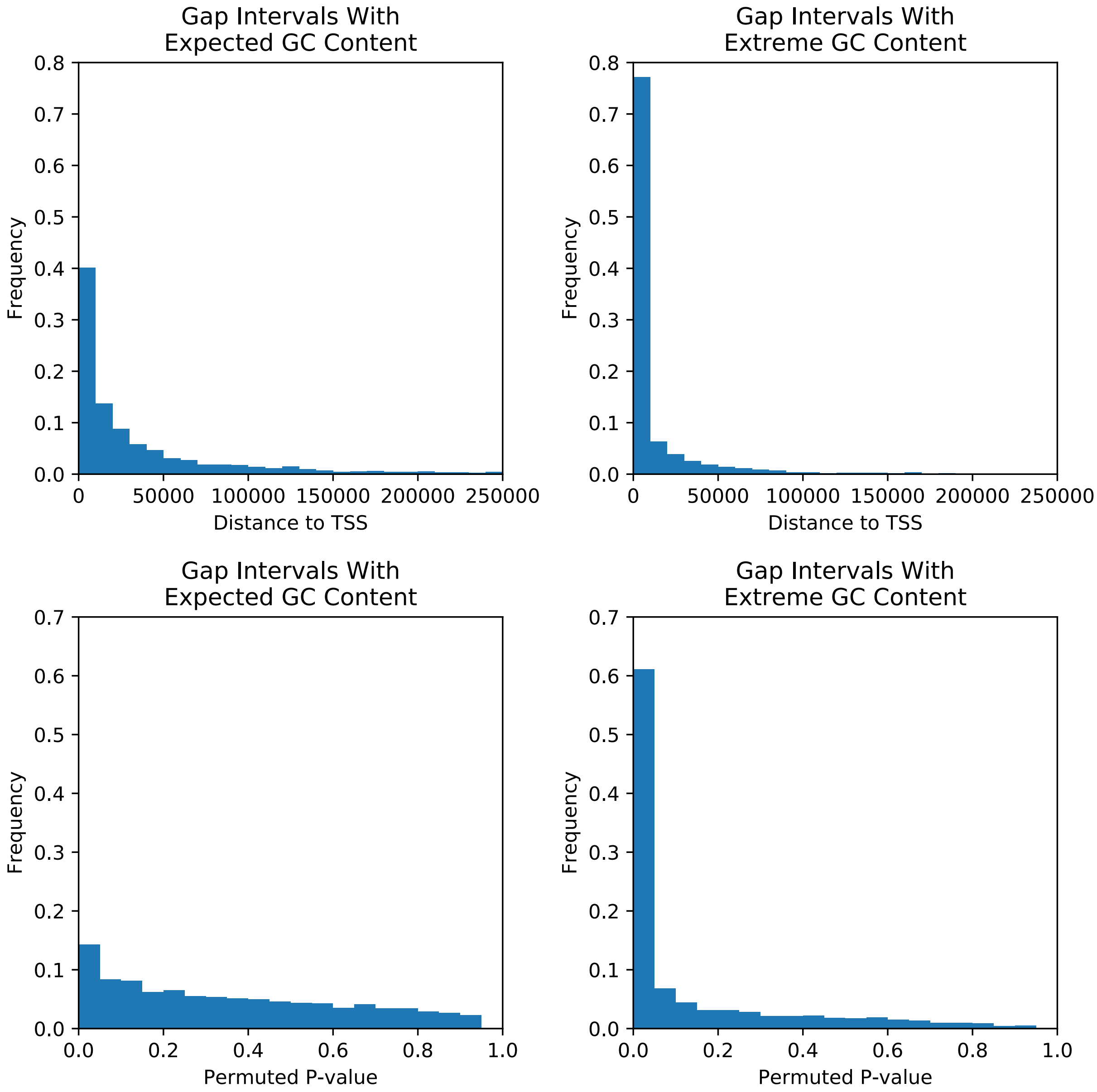


**Figure 4.11: Gap sequences with extreme GC content are located near transcription start sites.** We determined the distance from each gap segment and the transcription start site of the nearest protein-coding gene model based on the Zoey annotation. The top row shows histograms of distance to the nearest transcript start site (TSS). The bottom row shows the distribution of p-values determined by 1,000 permutations of the location of each gap interval. The first column shows results for the 7,253 intervals with a GC content that is similar to the genome wide expectation. The second column shows results for the 5,553 segments with an extreme GC content based on permutation.

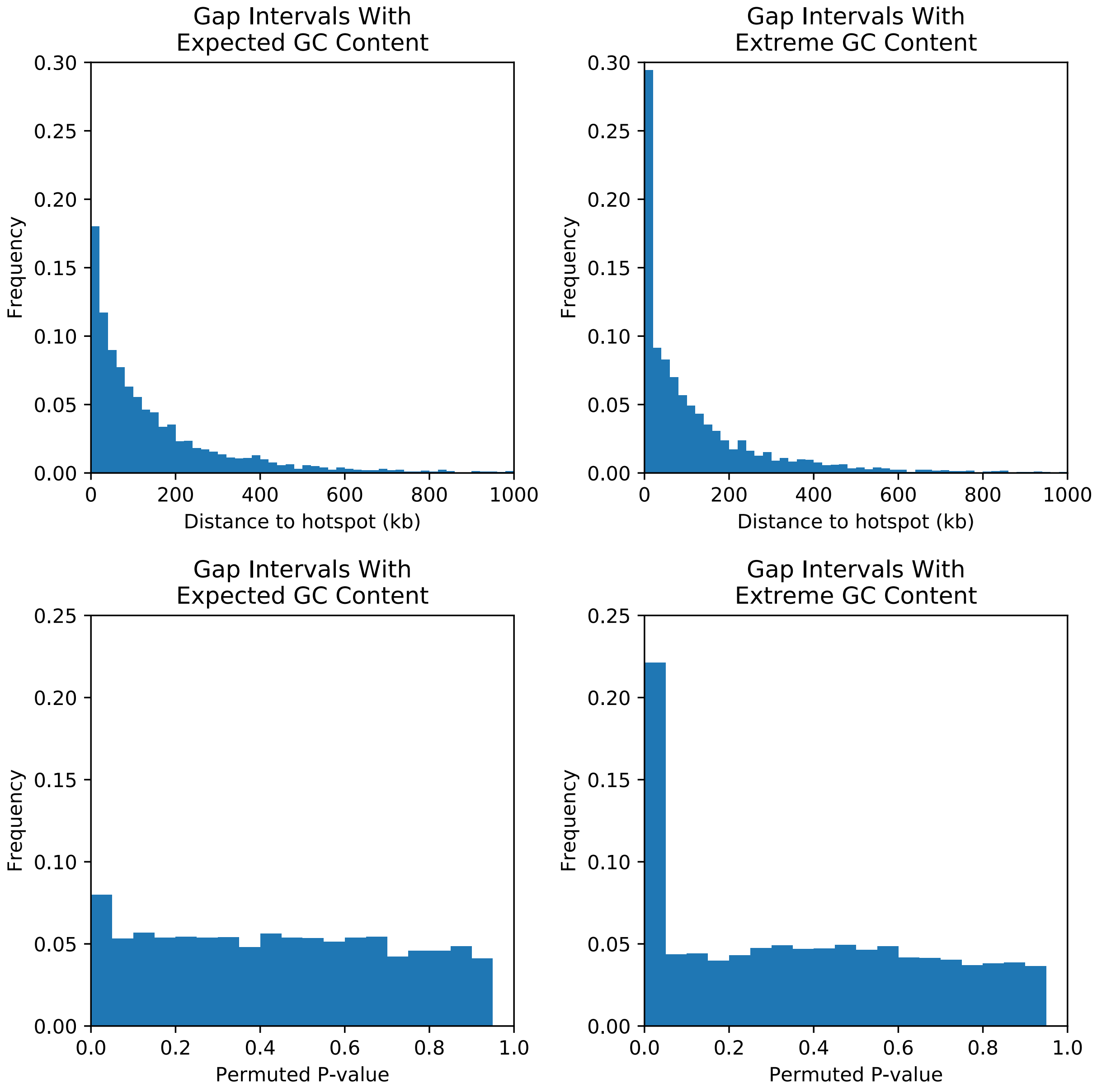


**Figure 4.12: Gap sequences with extreme GC content are located near recombination hotspots.** We determined the distance between each gap segment and recombination hotspots (from ref (56) ). The top row shows histograms of distance to the nearest hotspot. The bottom row shows the distribution of p-values determined by 1,000 permutations of the location of each gap interval. The first column shows results for the intervals with a GC content that is similar to the genome wide expectation. The second column shows results for the segments with an extreme GC content. This analysis was limited to segments located on the autosomes.

### 5. Analysis of Large Insertions and Deletions

*Filtering and breakpoint refinement*

The Zoey assembly, as well as the 6,857 secondary contigs, were aligned to CanFam3.1 using minimap2 (version 2.9-r720) with the -asm5 option (43). The output from the alignment was parsed using the paftools.js program released as part of minimap2 to identify candidate variants relative to the CanFam3.1 assembly. Given the presence of small insertion/deletion errors in PacBio reads, we focused our initial analysis on variants predicted to be at least 10 bp in size. We then refined the coordinates of the detected variant by performing target alignments of the flanking and variant sequences for each candidate. Flanking regions within 500 bp of each variant were extracted and aligned using the program AGE (57). In addition to base pair-level resolution of structural variants, AGE identifies segments of 100% sequence identity present at the breakpoints that match the variant allele. Redundant candidate calls from minimap2, where AGE assigned the same breakpoints, were removed from the analysis. Consistent with previous size cutoffs for structural variation, we focused our analysis on breakpoint-resolved structural variants predicted to be at least 50 bp in length. The candidate variants were subjected to additional filers to remove regions that may be problematic in the assembly (Table 5.1; Figure 5.1). Variants with more than 10% of their length intersecting with assembly gaps or with segmental duplications defined by self-alignment or defined by read depth in either the CanFam3.1 or Zoey assemblies were removed (Table 5.2, A-D, available as separate files in the online appendix). Additionally, we identified a small number of variants that appeared to be more complex than simple insertion-deletion events due to the presence of at least 10 bp on the ‘deletion’ allele that was not found on the ‘insertion’ allele (Table 5.3, A-D, available as separate files in the online appendix).

| | Primary Deletions | Primary Insertions | Secondary Deletions | Secondary Insertions |
|---|---|---|---|---|
| **minimap2 candidates** | 1,057,366 | 490,315 | 465,697 | 104,549 |
| **>=10 bp** | 74,320 | 62,929 | 9,248 | 9,032 |
| **AGE resolved breakpoints; >=50 bp** | 20,249 | 17,543 | 3,097 | 4,990 |
| **Less than < 10% overlap with segmental duplication** | 19,637 | 16,857 | 2,948 | 3,634 |
| **Involves unplaced sequences** | 6 | 4 | - | - |
| **>10% overlap with assembly gap** | 2,444 | 929 | 206 | 67 |
| **Complex variants/ block substitutions** | 353 | 303 | 77 | 74 |
| **Final simple insertion/deletions** | 16,834 | 15,621 | 2,665 | 3,493 |

**Table 5.1: Summary of structural variant identification and filtration.** Each row reflects a different stage of the filtration strategy. The columns depict the counts of insertions and deletions identified in the Zoey primary assembly and in the secondary Zoey contigs.

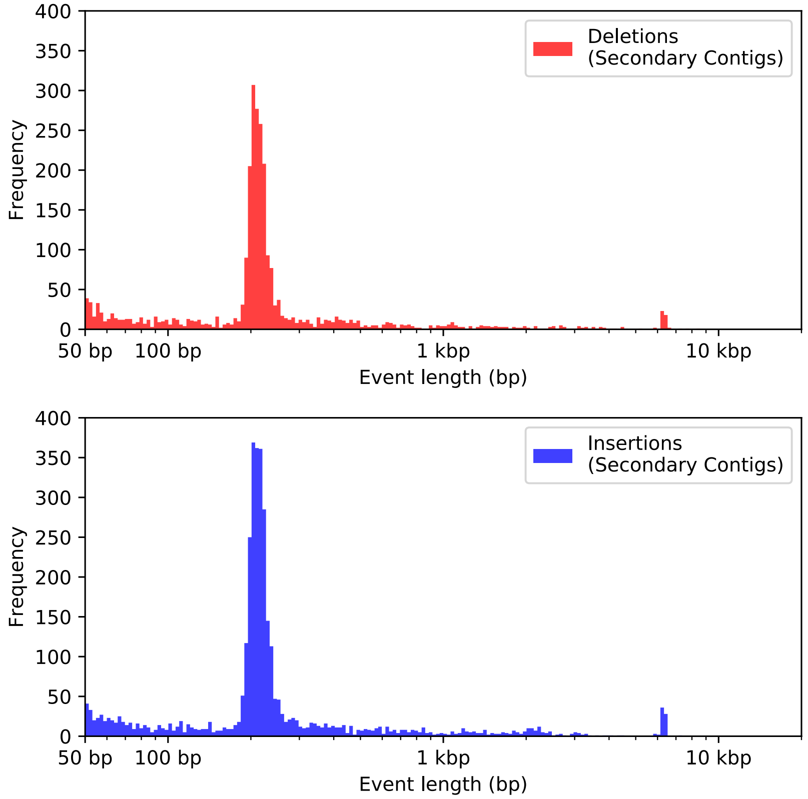


**Figure 5.1: Size distributions of insertion-deletion variants identified on the secondary contigs.** The secondary contigs were aligned to the CanFam3.1 reference genome and processed as described above. The plot shows histograms of variant size plotted on a logarithmic scale. The bins in the histogram are of equal size in the log scale.

*Dimorphic mobile element sequences*

We focused our subsequent analysis on the 38,613 simple insertion-deletion structural variants identified through our assembly comparison. We noted clear peaks in the size distribution that correspond to the expected size of canine SINE and LINE-1 elements, suggesting that much of the apparent insertion deletion differences could be due to retrotransposon insertions that differ between the Great Dane and Boxer genomes used for these assemblies. To investigate this possibility further, we performed RepeatMasker analysis of insertions and deletions in the 150-250 bp (for SINEs) and 1-7 kbp (for LINE-1s) size range. Because mechanisms other than retrotransposition can result in the differential presence/absence of specific retroelements between genomes, we required that at least 80% of inserted sequence be annotated as the indicate repeat type. We additionally focused on carnivore-specific SINEs (‘SINEC’ elements)(58). This analysis resulted in the identification of 16,221 dimorphic SINECs and 1,121 dimorphic LINEs (Tables 5.4, 5.5, 5.6, and 5.7).

The AGE aligner identifies segments of identical sequence that are found at variant breakpoints. These sequences approximate the length of target-site duplications (TSD) that generally are created during SINE and LINE retrotransposition. This initial analysis only provides an approximation of the TSD size, as the AGE aligner requires segments to have 100% sequence identity flanking the insertion as well as across the corresponding breakpoint on the deletion or empty site allele. Thus, a single base pair change in either the 5' or 3' TSD or at the corresponding segment on the non-insertion 'empty site' allele, will result in a failure to identify a candidate TSD. The TSD distribution is very similar for dimorphic LINE-1 and SINEC events, with each having a median size of 15 bp (Figure 5.2).

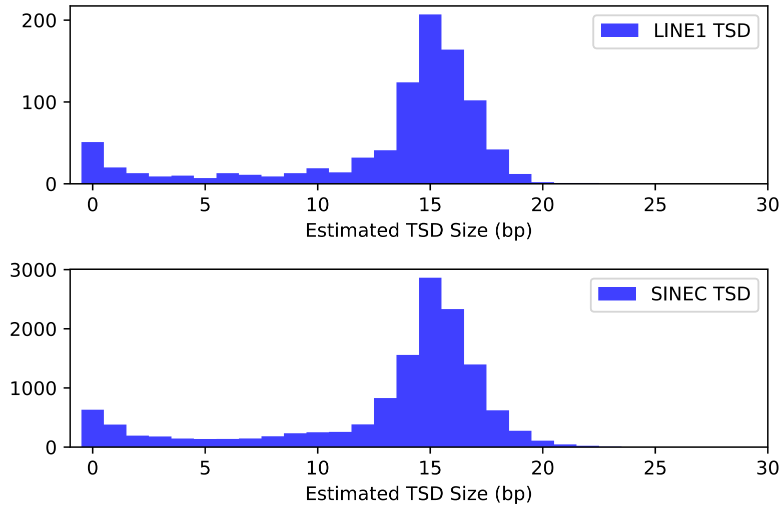


**Figure 5.2: Estimated target site duplication length for dimorphic SINEC and LINE-1 sequences.** Target site duplications (TSDs) were identified using the AGE aligner of 'insertion' and 'deletion' alleles. Segments are required to have 100% identity across all three breakpoint positions (flanking the insertion and at the empty site junction on the 'deletion' allele).

| RepeatMasker Annotation | Primary Deletions | Primary Insertions | Secondary Deletions | Secondary Insertions |
|---|---|---|---|---|
| SINEC1A_CF | 9 | 8 | 1 | 1 |
| SINEC1D_CF | 0 | 0 | 0 | 1 |
| SINEC1B1_CF | 0 | 1 | 0 | 0 |
| SINEC2A1_CF | 4823 | 3881 | 824 | 1048 |
| SINEC2A2_CF | 0 | 1 | 0 | 0 |
| SINEC_Cf | 2231 | 1940 | 396 | 487 |
| SINEC_Cf2 | 129 | 124 | 22 | 28 |
| SINEC_Cf3 | 79 | 75 | 10 | 20 |
| SINEC_a1 | 5 | 7 | 1 | 2 |
| SINEC_a2 | 16 | 28 | 2 | 5 |
| SINEC_b1 | 4 | 3 | 1 | 1 |
| SINEC_b2 | 1 | 3 | 2 | 0 |
| SINEC_c2 | 1 | 0 | 0 | 0 |
| **Total** | **7298** | **6071** | **1259** | **1593** |

**Table 5.4: RepeatMasker annotation of dimorphic Carnivore SINECs.** We only counted insertion-deletion variants within a size range of 150-250 bp where at least 80% of the sequence length matches a SINEC element.

| RepeatMasker Annotation | Primary Deletions | Primary Insertions | Secondary Deletions | Secondary Insertions |
|---|---|---|---|---|
| L1_Canid_ | 1 | 0 | 0 | 0 |
| L1_Canis1 | 53 | 101 | 11 | 13 |
| L1_Canis2 | 6 | 9 | 0 | 0 |
| L1_Carn5 | 0 | 1 | 1 | 0 |
| L1_Carn7 | 1 | 0 | 0 | 0 |
| L1_Carni | 0 | 0 | 1 | 0 |
| L1_Cf | 124 | 202 | 36 | 62 |
| L1M3 | 1 | 1 | 0 | 0 |
| L1M4a2 | 0 | 0 | 1 | 0 |
| L1M5 | 0 | 1 | 0 | 0 |
| L1MA10 | 0 | 1 | 0 | 0 |
| L1MB5 | 1 | 0 | 0 | 0 |
| L1MD | 0 | 2 | 0 | 0 |
| L1MDa | 0 | 1 | 0 | 0 |
| L1MEc | 152 | 262 | 25 | 51 |
| **Total** | **339** | **581** | **75** | **126** |

**Table 5.5: RepeatMasker annotation of dimorphic LINEs**. We only counted insertion-deletion variants with a size range of 1kb-7kb where at least 80% of the sequence length matches an annotated as a LINE.

*LINE-1 3' transduction analysis*

The polyadenylation signal encoded by LINE-1 elements is often bypassed by RNA polymerase II, resulting in the incorporation of genomic sequences located downstream of the element into the resultant LINE-1 mRNA transcript (59-61). As a result of retrotransposition, these flanking sequences are inserted into new genomic loci, resulting in an accompanying 3' transduction 'tag' sequence. The pattern of shared 3'-transduced sequences can be analyzed to identify relationships among LINE-1 elements in the genome (62) and identify the descendants of active progenitor elements (63-65). We searched the regions flanking the 3' end of each dimorphic LINE-1 to identify potential transductions. The 5' portion of the transduced sequences often contains A-rich sequence with a low-complexity. Thus, we defined the extent of 3'-transduction based on alignment of the candidate transduced sequence to the canFam3.1 assembly. We identified 18 events with a clearly defined 3'-transduction. Each of the 18 transduced sequences aligned elsewhere in the canine genome, and 17/18 aligned to a region that was not adjacent to a LINE-1 sequence in the reference. We surmise that a polymorphic, active LINE-1 likely exists at each of these respective genomic locations in the dog population and that these polymorphic LINE-1s gave rise to the observed 3'- transductions. Thus, the presence of the 17 "parentless" 3'-transductions suggest that additional, active LINE-1 copies, which were not identified in our analyses, are segregating among dogs (Figure 5.3). Moreover, we identified two LINE-1 lineages wherein the same 3'- transduced sequence was found in multiple elements suggesting the presence of an active canine LINE-1 that has spawned multiple new insertions. A summary of the transduction analysis is provided in Table 5.8, which is available as a separate file in the online appendix.

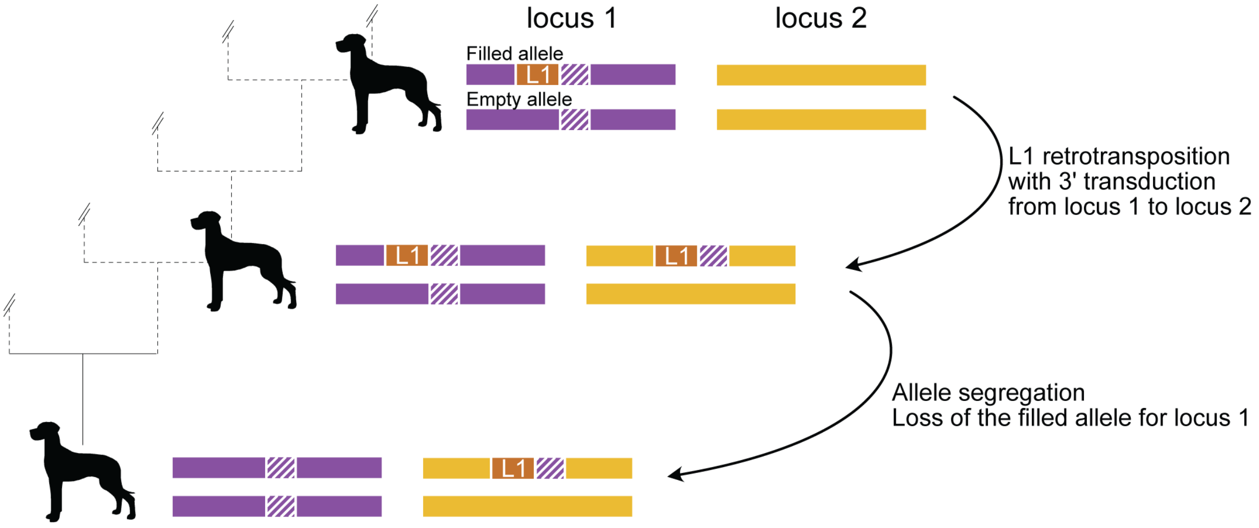


**Figure 5.3: Schematic of allele segregation leading to the appearance of "parentless" 3'-transductions.** The schematic depicts a hypothetical example. The top of the figure depicts a polymorphic active LINE-1 (brown box) at a specific location in the genome (locus 1, purple). At some point (middle of the figure), this LINE-1 is expressed and generates a new copy of itself at locus 2 (orange), which includes a 3' transduction (dashed purple box). The LINE-1 insertion at locus 2 may be full-length or contain a 5' truncation. Because of independent segregation, some individuals may inherit the LINE-1 insertion containing the 3' transduction at locus 2 without inheriting the LINE-1 sequence at locus 1 (bottom of the figure). Several examples found in the Zoey genome follow this pattern; the surmised parent LINE-1 at locus 1 is absent from the Boxer-derived CanFam3.1 reference genome. These data suggest the existence of additional, active LINE-1 sequences that are yet to be characterized.

*Annotation of transduced sequences*

Below, we present an annotated view of the sequence of each LINE-1 variant contains a 3'-transduction. The Site IDs correspond to the information in Table 5.8. In each case, sequence matching the canine L1_Cf element is colored in red (the internal portions of large insertions are omitted as indicated by … to save space), the transduced sequence that matches elsewhere in the CanFam3.1 genome is colored in blue, and the target site duplications that flank each element are annotated in **bold underline**. Each sequence was additionally compared to the corresponding empty site sequence (not shown).

**Trans_1**

>CanFam chr3:56250820-56257252
AATACAATGGATGAACTGAATAGCAAAGTAGGTCTCACTGAAGAATGAATTAGTACGCAG
AACAGGTTAGGACACTGTCTC**AAAGGCATTAGCA**GGGGAGGAGCAAGATGGCGGAAGAGT
AGGGTCTCCAAATCACCTGTCTCCACCAAACTACCTAGAAAACCTTCAAATTATCCTGAA
AATCTATGAATTCGGCCTGAGATTTAAAGAGAGACCAGCTGGAATGCTACAGTGAGAAGA
GTTCGCGCATCTATCCAGGTAGGAAGACGGGGGAAAAAGAAATAAAGGAACAAAGGCCTC
CAAGGGGGAGGGGCCCGCGAGGAGCCGGGCTGAGGCCGGGGCGAGTGTCCCCAGGACAGG
AGAGCCCCGTCCCGGAGACGCAGGAGCTGCACCGACCTTCCCGGGCGGAAAGGGGCTCGC

AGGGGGTTGGAGCAGGAACCAGGAGGGCGGGGATGCCCTCGGGCTCCCGGGGACACTAAC
AGACACCTGCGCCCGGGAGAGTGCGCCGAGCTCCCTAAGGGCTGCAGCGCGCACGGCGGG
ACCCGGCGGGACCCGGAGCAGCTGAAGGGGCTCGGGCAGCGGCTCCGCGGAGGGGGCTGC
GCGGCCCCGGGAGCAGCTCGGAGGGGCTCGGGCAGAGGAAGAGGCTCCGTGCGGAGGGGG
CTGCGCGGCCCGGGAGCGCGAATCCAACAGCGCAGGCTCCGGAGCACAGGGCGCCGGGAC
…
GTTGGGGATTTACCCCAAAGATACAAATGCAATGAAACGCCGGGACACCTGCACCCCGAT
GTTTCTAGCAGCAATGGCCACGATAGCCAAACTGTGGAAGGAGCCTCGGTGTCCATCGAA
AGATGAATGGATAAAGAAGATGTGGTTTATGTATACAATGGAATATTACTCAGCTATTAG
AAATGACAAATACCCACCATTTGCTTCAACGTGGATGGAACTGGAGGGTATTATGCTGAG
TGAAGTAAGTCAGTCGGAGAAGGACAAACATTATATGTTCTCATTCATTTGGGGAATATA
AATAATAGTGAAAGGGAATATAAGGGAAGGGAGAAGAAATGGGTGGGAAATATCAGAAAG
GGAGACAGAACGTAAAGACTGCTAACTCTGGGAAACGAACTAGGGGTGGTAGAAGGGGAG
GAGGGCGGGGGGTGGGAGTGAATGGGTGACGGGCACTGGGTGTTATTCTGTATGTTAGTA
AATTGAACACCAATAAAAAAAATAAAAATAAAAATAAAAAAAAAAAATAAAAAAAATAAA
ACTCCAGCACCTAGCAAATTGAACACCAATAAAAATAAATTTATAAATAAAAAAAAAAAA
AAA**AAAGGCATTAGCA**AAGGGTAGAACAACAGAAAAAGCAGGAGAAAAGTTGACAGATGG
AGGATACTGTAGA

**Trans_2**
>CanFam chr23:9561915-9568581
CCTTGTCCTTCATTTTTTTGATAATCAGAAATTGTACATTTTAATGTAGTAAATTTTTTT
TATAAGTGGATCTTATTTTTTT**TAATTTTTTTT**TTTATATTATATAGTGTTTATTTTAAA
TTATATTTTGAACTTCAATTAATTAGACAAGATTATAGTGAAGTATGTTAATGTTAAATA
TATAAATTCTTTCAAAGGATTTTAAACTTTGTTTCCCATACATGAAATCATCTACATTTA
AATTTAGCGTCAGAATCTTTTCCTTATTTCCTATTGGACTTCTTTTTTTTTTTCTTTCTT
TTTTTTTTTTTTTCTTTTTTTTTTTTTTTATTGGGGTTCAATTTACTAACATACAGAATAA
CACCCAGTGCCCGTCACCCATTCACTCCCACCCCCCGCCCTCCTCCCCTTCTACCACCCC
TAGTTCGTTTCCCAGAGTTAGCAGTCTTTATGTTCTGTCTCCCTTTCTGATATTTCCCAC
ACATTTCTTCTCCCTTCCCTTATTTTCCCTTTCACTATTATTTATATTCCCCAAATGAAT
GAGAACATATAATGTTTGTCCTTCTCCGACTGACTTACTTCACTCAGCATAATACCCTCC
AGTTCCATCCACGTTGAAGCAAATGGTGGGTATTTGTCATTTCTAATAGCTGAGTAATAT
TCCATTGTATACATAAACCACATCTTCTTTATCCATTCATCTTTCGTTGGACACCGAGGC
TCCTTCCACAGTTTGGCTATCGTGGCCATTGCTGCTAGAAACATCGGGGTGCAGGTGTCC
CGGCGTTTCATTGCATTTGTATCTTTGGGGTAAATCCCCAACAGTGCAATTGCTGGGTCG
…
TCAGCCCGGCTCCTCCTGGGCCCCTCCCCCTTGGAGGCCTTTGTTTCTTTATTTCTTTTT
CCCCGTCTTCCTACCTTGATAGATGCGCGAACTCTTCTCACTGTAGCATTCCAGGTGGTC
TCTCTTTAAATCTCAGGCCGAATTCATAGATTTTCAGGATAATTTGAAGGTTTTCTAGGT
AGTTTGGTGGAGACAGGTGATTTGGAGACCCTACTCTTCTGCCATCTTGCTCCTCTC**TAA**
**TTTTTTTT**ATTTATTTATGATAGTCACAGAGAGAGAGAGAGAGAGAGAGAGAGAGAGAGA
GAGAGAG

**Trans_3**
>CanFam chr16:8006641-8008015
TTTGAGTTTCCCATT**TAAACCATCGAGCCCA**TCCACATGCAGAAGAATGAAACTAGACCA
CTCTCTTTCACCATACACAAAGATAAACTCAAAATGGATGAAAGATCTAAATGTGAGACA
AGATTCCATCAAAATCCTAGAGAAGAACACAGGCAACACCCTTTTTGAACTCGGCCATAG
TAACTTCTTGCAAGATACATCCACGAAGGCAAAAGAAACAAAAGCAAAAAATGAACTATTG
GGACTTCATCAAGATAAGAAGCTTTTGCACAGCAAAGGATACAGTCAACAAAACTCAAAG
ACAACCTACAGAATGGGAGAAGATATTTGCAAATGACGTATCAGATAAAGGCTAGTTTC

CAAGATCTATAAAGAACTTATTAAACTCAACACCAAAGAAACAAACAATCCAATCATGAA
ATGGGCAAAAGACATGAACAGAAATCTCACAGAGGAAGACATAGACATGGCCAACATGCA
TATGAGAAAATGCTCTGCATCACTTGCCATCAGGGAAATACAAATCAAAACTACAATGAG
ATACCACCTCACACCAGTGAGAATGGGGAAAATTAACAAGGCAGGAAACAACAAATGTTG
GAGAGGATGCGGAGAAAAGGGAACCCTCTTACACTGTTGGTGGGAATGTGAACTGGTGCA
GCCACTCTGGAAAACTGTGTGGAGGTTCCTCAAACAGTTAAAAATAGACCTGCCCTACGA
CCCAGCAATTGCACTGTTGGGGATTTACCCCAAAGATACAAATGCAATGAAACGCCGGGA
CACCTGCACCCCGATGTTTCTAGCAGCAATGGCCACGATAGCCAAACTGTGGAAGGAGCC
TCGGTTTCCAACGAAAGATGAATGGATAAAGAAGATGTGGTTTATGTATACAATGGAATA
TTACTCAGCTATTAGAAATGACAAATACCCACCATTTGCTTCAACGTGGATGGAACTGGA
GGGTATTATGCTGAGTGAAGTAAGTCAGTCGGAGAAGGACAAACATTATATGTTCTCATT
CATTTGGGGAATATAAATAATAGTGAAAGGGAAAATAAGGGAAGGGAGAAGAAATGTGTG
GGAAATATCAGAAAGGGAGACAGAACGTAAAGACTACTAACTCTGGGAAACGAACTAGGG
GTGGTAGAAGGGGAGGAGGGCGGGGGGTGGGAGTGAATGGGTGACGGGCACTGGGTGTTA
TTCTGTATGTTAGTAAATTGAACACCAATAAAATATAAAAAAAAAAGAAAGAAAGAAAGA
AAGACATACAGAAGACTATTAAGGGCTTGCAAAAAAAAAGTTAAGATCAAAAAAATAAAA
AATAAAAAAA**TAAACCATCGAGCCCA**AATAAATAAATAAATAAATAAACCATCGA

**Trans_4a**

Note that Trans_4a and Trans4_b contain the same transduced sequence.

>CanFam chr25 rc 13010319-13016958 Reversed:
ATGGGAACGTGCAGGGGCGCTGCTGGTGGAAACCCAGACCCCTCGGTGGCGCTGCTGCAA
AATAAACAACACAGACAGGGCCAAGTGGCCAGCAGCACTGTCTGGGCGCCCATG**GAGAAC
CAGGAGAAGGG**GAGGAGCAAGATGGCGGAGGAGTAGGGTCTCCAAATCACCTGTCTCCAC
CAAACTACCTAGAAAACCCTCAAATTATCCTGAAAATCTATGAATTCGGCCTGAGATTTA
AAGAGAGACCAGCTGGAATGCTACAGTGAGAAGGGTTCGCGCTTCTATCAAGGTAGGAAG
ACGGGGAAAAAGAAGTAAAGAAACAAAGGCCTCCAAGGGGGAGGGGCCCCGCGAGGAGCC
GGGCTGAGGCCGGGGCGAGTGTCCCCAGGACAGGAGAGCCCCGTCCCGGAGGAGCAGGAG
CTGCACCGACCTTCCCGGGCGGAAAGGGGCTCGCAGGGAGGTGGAGCAGGACCCAGGAGG
GCGGGGATGCCCTCGGGCTCCCGGGGACAGTAACAGCAACTGCGCGCCCAGGAGAGTGCG
…
CTCTGGGAAACGAACTAGGGGTGGTAGAAGGGGAGGAGGGCGGGGGGTGGGAGTGAATGG
GTGACGGGCACTGGGTGTTATTCTGTATGTTAGTAAATTGAACACCAATAAAAAAAAATA
ATTAAAAAAAAAAAAATAAAGATGAGGAGGTTGAGGTCCTCAGATGAAAAAAAAAAAAAA
AAAAAAAAAAA**GAGAACCAGGAGAAGGG**TCTGGAGGGATCCCCGGGCCTGGCCTGTTGCT
GGAGGAGGTGACCCCTCGCCCCAGAGGAAGCTCTGTGCACACTCTGGAGGAGCATCCTTC
CCAGCAGCTAGGAGGCCCTAGTGAGGGGCGTGCAGAGGCG

**Trans_4b**

>CanFam chrX:28704914-28711396
CACGCCCTGGCTGAAGGTGGCACTAAACCTCTGAGCCACTGGGGCTGCCCTGTGCACGCG
TTTATGTGTTA**CAAATTTTTTTATG**GGGGGGGAGGAGCAAGATGGCGGAAGAGCAGGGT
CTCCAAATCACCTGTCTCCACCAAACTACCTAGAAAACCCTCAAATTATCCTGAAAATCT
ATGAATTCGGCCTGAGATTTAAAGAGAGACCAGCTGGAATGCTACAGTGAGAAGGGTTCG
CGCTTCTATCAAGGTAGGAAGACGGGGAAAAAGAAGTAAAGAAACAAAGGCCTCCAAGGG
GGAGGGGCCCCGCGAGGAGCCGGGCTGAGGCCGGGGCGAGTGTCCCCAGGACAGGAGAGC
CCCGTCCCGGAGACGCAGGAGCTGCACCGACCTTCCCGGGCGGAAAGGGGCTCGCGGGGA
GGTGGAGCAGGACCCAGGAGGGCGGGGATGCCCTCGGGCTCCCGGGGACAGTAACAGCAA
CTGCGCGCCCAGGAGAGTGCGCCGAGCTCCCTAAGGGCTGCAGCGCGCACGGCGGGACCC
GGCGGGACCCGGAGCAGCCGAAGGGGCTCGGGCGGCGGCTCCGCGGAGGGGGCTGCGCGG
…

ATGGATAAAGAAGATGTGGTTTATGTATACAATGGAATATTACTCACCTATGAAAAATGA
CACATACCCACCATTTGCTTCAACGTGGATGGAACTGGACGGTATTATGCTGAGTGAAGT
AAGTCAGTCGGAGAAGGACAAACATTATATGTTCTCATTCATTTGGGGAATATAAATAAT
AGTGAAAGGGAAAATAAGGGAAGGGAGAAGAAATGTGTGGGAAATATCAGAAAGGGAGAC
AGAACGTAAAGACTGCTAACTCTGGGAAACGAACTAGGGGTGGTAGAAGGGGAGGAGGGC
GGGGGGTGGGAGTGAATGGGTGACGGGCACTGGGTGTTATTCTGTATGTTAGTAAATTGA
ACACCAATAAAAAAAATAATTAAAAAAAAAAAAAATAAAGATGAGGAGGTTGAGGTCCTC
AGATGAAAAAAAAAAAAAAAAAAAAAAAAAAAAAAC**CAAATTTTTTTTATG**ATTTTGTTTATT
CAT

**Trans_5**
>CanFam chr32:8886457-8889416
GATTTATAAAGCTAATGT**GATGCTTCCAAGGTTT**TTTTTTTTTTTTTTTTTTTTTTAACA
TATAGCATTATATATTAGTTTCATGTGTACAATATAATGATTTAACAATTCTATGCATTG
CTTGGAGCTCATGATGATAATTGTACTCTTAATCCTCTTCACCTATTTCACCCGTTCCCC
ACTCACTTCCCCTCTGGTATCTGTCTGTACTCTAGAGTATGTTTTTTTTTCCTCTTTGTC
TTTTTTTTTTTTTTTTAATTTATTTTTTATTGGGGTTCAATTTACTAACATACAGAATAA
CCCCCAGTGCCCGTCACCCATTCACTCCCACCCCCCGCCCTCCTCCCCTTTTACCACCCC
TATTTCATTTCCCAGAGTTAGCAGTCTTTACGTTCTGTCTCCCTTTCTGATATTTCCCAC
…
GGTATGATTTCTTCTTTAAACGTTTGATAAAATTCCCCTGGGAAGCCATCTGGCCCTGGA
CTCTTGTGTCTTGGGAGGTTTTTGATGACTGCTTCAATTTCCTCCCTGGTTATTGG**GATGC
TTCCAAGGTTT**AATCAGAG

**Trans_6**
>Zoey chr1:26791625-26798188
CTCCAAGTCCTGGGATGGAGCCCTGAATGGAGCTCTCTGTTTCTCCCTCTCCCTCTGCCC
A**CCGCTCTGCCTTCTT**TTTTTTTTTTTTTTCATTACGAAAAAAAGATTTTATTTTTTAAA
GAATCTCTACACTCAACTTGGGGCTCGAACTTACAACCCTGAGACCAAGTCTCATGCTCT
ACTAAGCCAGCCAGGTGCCCCAATAAATATTCAAATTATAATTTGAGAAGTTTTCAGTTT
CATTCGGCCTTCTTTTTTTTTTTTTTTTTTTTTTTTTATTGGTGTTCAATTTACTAACAT
ACAGAATAACACCCAGTGCCCGTCACCCATTCACTCCCACCCCCGCCCTCCTCCCCTTCT
ACCACCCCTAGTTCGTTTCCCAGAGTTAGCAGTCTTTACGTTCTGTCTCCCTTTCTGATA
TTTCCCACCCATTTCTTCTCCCTTCCCTTATTTTCCCTTTCACTATTATTTATATTCCCC
AAATGAATGAGAACATATAATGTTTGTCCTTCTCCGACTGACTTACTTCACTCAGCATAA
…
GGTCCCGCCGGGTCCCGCCGTGCGCGCTGCAGCCCTTAGGGAGCTCGGCGCACTCTCCTG
GGCGCGCAGTTGCTGTTACTGTCCCGGGAGCCCGAGGGCATCCTCGCCCCCTGGGTCCTG
CTCCACCTCCCACGAGCCCCTTTCCCCGGGAAGGTCGGTGCAGCTCCTGCTCCTCCGGGA
CGGGGCTCTCCTGTCCTGGGGACACTCGCCCCGGCCTCAGCCCGGCTCCTCGCGAGGCCC
TCCCCCTTGGAGGCCTTTGTTTCTTTATTTCTTTTTCCCCGTCTTCCTACCTTGATAGAA
GCGCGAACTCTCCTCACTGTTGCATTCCAGCTGGTCTCTCTTTAAATCTCAGGCCGAATT
CATAGATTTTCAGGATAATTTGAAGGTTTTCTAGGTAGTTTGGTGGAGACAGGTGATTTG
GGGACCCTGCTCTTCCGCCATCTTGCTCCTCCCC**CCGCTCTGCCTTCTT**GTGCTTGCGCT
CTCTCTCTGTTAAATAATTAAAAA

**Trans_7**
>Zoey chr2:44763614-44770272
GAATGGGAAAAGGTAATTGCAAATGTTATATTCGATAAGGGGTTAATATCCAAAAATTCA
T**AAAGAACTTACACAACTC**GGGGGGAGGAGCAAGATGGCGGAAGAGTAGGGTCTCCAAAT
CACCTGTCTCCACCAAACTACCTAGAAAACCTTCAAATTATCCTGAAAATCTATAAATTC

GGCCTGAGATTTAAAGAGAGACCAGCTGGAATGCAACAGTGAGAAGAGTTCGCGCTTCTA
TCAAGGTAGGAAGACGGGGAAAAAGAAATAAAGAAACAAAGGCCTCCAAGGGGGAGGGGC
CCCGCGAGGAGCCGGGCTGAGGCCGGGGCGAGTGTCCCCAGGACAGGAGAGCCCCGTCCC
GGAGGAGCAGGAGCTGCACCGACCTTCCCGGGCGGAAAGGGGCTCGCGGGGAGCTGGAGC
…
GAAATGACAAATACCCACCATTTGCTTCAACGTGGATGGAACTGGAGGGTATTATGCTGA
GTGAAGTAAGTCAGTTGGAGAAGGACAAACATTATATGTTCTCATTCATTTGGGGAATAT
AAATAATAGTGAAAGGGAATATAAGGGAAGGGAGAAGAAATGTGTGGGAAATATCAGAAA
GGGAGACAGAACGTAAAGACTGCTAACTCTGGGAAACGAACTAGGGGTGGTAGAAGGGGA
GGAGGGCGGGGGTGGGAGTGAATGGGTGACGGCACTGGGTGTTATTCTGTATGTTAGTA
AATTGAACACCAATAAAAATAAATTAAAAAAAAAAAAAAGAAAATGAAAAGAAGTAATAT
ATGCAGCTGGCCATTCCCAATTAAGCAACAGCTCCTTGGTTAGCAGTAGTCTGAACCTGC
AGCATTTGGGTTGTTTTTCTGCTGAGCTATTTAAGATGTTCATTAGTTCAAAAAGAAATC
AGCAAGCAAGCTGCTAATAAAGGTAGTAAATTTGCTTCTAAAAAAAAAAAAAAAAAAAA**A**
**AAGAACTTACACAACTC**AACACCAAAAATAAATAAATAAATAATCCAGTTTAAAAATGT

**Trans_8**
>Zoey chr5:34110431-34112681
AGCAAGAAAAAGAAATAAAAGTCATCCGGATTGAAATGACGAAGTAAAACCACAATGATT
T**AAAGAAAAAGACTATTA**CAAACTCTCCCTCTTCGCCGATGACATGATACTCTACATAGA
AAACCCAAAAGTCTCCACCTCAAGATTGCTAGAACTCATACAACAATTCGGTAGCGTGGC
AGGATACAAAATCAATGCCCAGAAGTCAGTGGCATTTCTATACACTAACAATGAGACTGA
AGAAAGAGAAATTAAGGAGTCAATCCCATTTACAATTGCACCCAAAAGCATAAGATACCT
…
CTCATTCATTTGGGGAATATAAATAATAGTGAAAGGGAATAGAAGGGAAGGGAGAAGAAA
TGTGTGGGAAATATCAGAAAGGGAGACAGAACGTAAAGACTGCTAACTCTGGGAAACGAA
CTAGGGGTGGTAGAAGGGGAGGAGGGCGGGGGTGGGAGTGAATGGGTGACGGCACTGG
GTGTTATTCTGTATGTTAGTAAATTGAACACCAATAAAAAATAAATTAAAAAAAAAAAAA
AAAAAAAAAAAGAAAAGATGATCCAGAATGAAAACTATGAAAACTATGTTTGAGGCTTAG
GGGCCCAGGATTCACATTATATCAGGTGGTTAGAGAGCTCCAGGAGAGATATTGATGAGG
AAAAAATAAAATTGTTACAATTCCTTAAAAAAAAAAAAAAA**AAAGAAAAAGACTATTA**GA
GTGGATCAAAAAAAAAAGAGTCAACTACATG

**Trans_9**
>Zoey chr8:45413459-45417490
CTGGGATCTTTGCCTTACCCTTCTGCTATTTCATGGTTTCTTTGGTTCCTACTCTACTAT
C**TCTGGTTGATTTTT**TTTTTTTTTTTTTTTTTTATCACATTTGTTTTATTAACACATCT
TATAAATTATAATCTTACTTTCTCCAACATCTCTTTTGAGTTCTATCTATATGTCATAAG
CTACTTGCTTATTCATACCTATACTTAACATATATTATGAGATTTGTCTCTCTCTGATAA
ACTACTACCTGTTCACTATCTTGTTTAACATTCTTACAAGCTTCATTACTCCTGTTTGTT
CTTCTTTTATAACAGAATTGAATATTTTTGAATAATAAAGCAAACACACATTCCAACAAC
TTCCTCCATAATTGAAGGACAGGAACTTGTTATAGAGGTAAAAGAGACCTCTGTCACTTT
CATTATTCACTTTAGATGACTTTATACATTTTTTTTTTTTTTTGGTTTATTTTTCTTTT
TTTTTTTTTTCTTTTTTTTTTTTTTATTTTTTTTTTTTTATTGGTGTTCAATTTACTAACA
TACAGAATAACACCCAGTGCCCGTCACCCATTCACTCCCACCCCCCGCCCTCCTCCCCTT
CTACCACCCCTAGTTCGTTTCCCAGAGTTAGCAGTCTTTATGTTCTGTCTCCCTTTCTGA
TATTTCCCACACATTTCTTCTCCCTTCCCTTATTTTCCCTTTCACTATTATTTATATTCC
…
TTGAGTTCTTTCCAGTTTTTGAATGGATGCTTGTATTGCGATGTATTTCCCCCTTAGGAC
AGCTTTCGCTGCATCCCAAAGATTTGAACGGTTGTATCTTCATTCTCATTAGTTTCCAT
GAATCTTTTTAATTCTTCCTTAATTTCCTGGTTGACCCTTTTATCTTTTAGCAGGATGGT

CCTTAACCTCCATGTGTTTGAGGTCCTTCCAAACTTCTTGTTGTGATTTAGTTCTAATTT
CAAGGCATTATGGTCCGAGAATATGCAGGGGACAATCCCAATCTTTTGGTATCGGTTCAG
ACCCGATTTGTGACCCAATATGTGGTCTATTCTGGAGAAAGTTCCATGTGCG**TCTGGTTG**
**ATTTTT**GATACTGCACTATCATAGTGCCAAGGGCTACTATATATCCATAATTTCTTGACA
TCACATAATGTG

**Trans_10**
>Zoey chr11:16365277-16369597
TGCAAAACCACATCACTGGTATATTACCAAT**GCTATATATATTTTA**TTTTATTTTTTTT
TGGTTCATAATTCCAGTTTTAATATAGGAGTTAAAAGATAAAAGCACAAAAACAATTATA
AATCTATATTAGTGGGTATACAATATATATTTGCTGTGGGCTTTTCATAGATGGCTTTTA
AGATGTCGAGGAATGTTCCCTCTATCCCTACACTCTGAAGAGTTTTGATCAGGAATGGAT
GCTGTATTTTGTCAAATGCTTTCTCTGCATCCAATGAGAGGATCCTATGGTTCTTGGTTT
TTCTCTTGCTGATATGATGAATCACATTGATTGTTTTACGGGTGTTGAACCAGCCTTGTG
TCCCAGGGATAAATCCTACTTGGTCATGGTGAATAATTTTCTTAATGTACTGTTGGATCC
TATTGGCCAGTATCTTGTTGAGAATTTTTGCATCCATGTTCATCAGGGATATTGGTCTGT
AATTCTCCTTTTTGGTGGGGTCTTTGTCTGGTTTTGGAATTAAGGTGATGCTGGCCTCAT
…
GGGAGGCCGGATCCTGGGCTGTGTCCCGGCGCCCTGTGCTCCGGAGCCTGCGCTGGTGGA
TTCGCGCTCCCGGGCCGCGCAGCCCCCTCCGCGGAGCCGCCCCCGAGCCCCTGAGCTGCT
CCAGGAACCGCGCAGCCCCCTCCGCACGGAGCCTCTTCCTCTGCCCGAGCCCCCCCGAGC
TGCTCCCGGGGCCGCGCAGCCCCCTCCGCGGAGCCGCCGCCCGAGCCCCCCGAGCTGCTC
CGGGTCCCGCCGTGCGCGCTGCAGCCCTTAGGGAGCTCGGCGCACTCTCCTGGGCGCGCA
GTTGCTGTTACTGTCCCCGGGAGCCCGAGGGCATCCCCGCCCTCCTGGGTCCTGCTCCAC
CTCCCCGCGAGCCCCTTTCCCCCGGGAAGGTCGGTGCAGCTCCTGCTCCTCCGGGACGGG
GCTCTCCTGTCCTGGGGACACTCGCCCCGGCCTCAGCCGGGCTCCTCGCGGGCCCCTCCC
CCTTGGAGGCCTTTGTTTCTTTACTTCTTTTTCCCCGTCTTCCTACCTTGATAGAAGCGC
GAACTCTCCTCACTGTTGCATTCCAGGTGTTCTCTCTTTAATTCTCAGGCCGAATTCATA
GATTTTCAGGATGATTGGAAGGTTTTCTAGGTAATTTGGTGGAGACAGGTGATTTGGGGA
CCCTACTCTTCCGCCATCTTGCTCCTCCCCC**GCTATATATATTTTA**AAAAGCGAAGGAAA
T

**Trans_11**
>Zoey chr12:35127147-35133370
TATTTTTCTGAGTACTGTACGTTTTAGTTATATTCATAGTG**CCTATGTGACTATTTT**TTT
TTTTTTTTTTTTTTTTGATTCAGGTATAGTGTTTTTGAGCGCATCTCTTTGTATGGACT
TTTCAGTCATTGCAGTTAAGCTTACTTAAATTATCCACCTACTGGTGTCAACATTTTTGT
CATTTTATTTTATTTTATTTATTTATTTTTTTTTATTGGTGTTCAATTTACTAACATACA
GAATAACACCCAGTGCCCGTCACCCATTCACTCCCACCCCCCACCCTCCTCCCCTTCTAC
CACCCCTAGTTCGTTTCCCAGAGTTAGCAGTCTTTACGTTCTGTCTCCCTTTCTGATATT
TCCCACACATTTCTTCCCCCTTCCCTTCTTTTCCCTTTCACTATTATTTATATTCCCCAA
ATGAATGAGAACATATAATGTTTGTCCTTCTCCGACTGACTTACTTCACTCAGCATAATA
CCCTCCAGTTCCATCCACGTTGAAGCAAATGGTGGGTATTTGTCATTTCTAATAGCTGAG
…
CCCCCCGAGCTGCTCCGGGTCCCGCCGAGCGCTGCAGCCCTTAGGGAGCTCGGCGCACTC
TCCTGGGCGCGCAGTTGCTGTTACTGTCCCGGGGAGCCGGAGGGCATCCCCGCCCTCCTG
GGTCCTGCTCCAACTCCCCGCGAGCCCCTTTCCGCCCGGGAAGGTTGGTGCAGCTCCTGC
GTCTCCGGGGCGGGGCTCTCCTGTCCTGGGGACACTCGCCCCGGCCTCAGCCCGGCTCCT
CGCC**CCTATGTGACTATTTT**AATTACTGGTTTAATGAAAGCTAA

**Trans_12**
>Zoey chr13:41174201-41180947
TGGAAGAGCCCAGATGCACATCAACAGATGAATGGATAAAG**AAGATGTGA**GGGGGAGGAG
CAAGACGGCGGAAGAGTGGGGTCTCCAAATCACCTGTCCCAACCAAATTACCTACAAAAC
CTTCAAATTACCCTGAAAATCTATGAATTCGGCCTGAGAATTAAAGAGAGAACACCTGGA
ATGCTACAGTGAGAAGAGTTGGCGCTTCTATCAAGGTAGGAAGACGGAAAAAAGAAATAA
AAAAACAAAGGCCTCCAAGGGGGAGGGGCCCCGCGAGGAGCCGGGCTGAGGCCGGGGCGA
GTGTCCCCAGGACAGGAGAGCCCCGTCCCGGAGACGCAGGAGCTGCACCGACCTTCCCGG
GGGAAAGGGGCTCGCGGGGAGTTGGAGCAGGACCCAGGAGGGCGGGGATACCCTCGGGCT
CCCTGGGACAGTAACAGAGCAACTGCGCGCCCAGGAGAGTGCGCCGAGCTCCCTAAGGGC
TGCAGCGCGCACGGCGGGACCGGCGGGACCCGGAGCAGCTCGGAGGGGCTCGGGCGGCGG
CTCCGCGGAGGGGGCTGCGCGGCCCCGGGAGCAGCTCGGAGGGGCTCGGGCAGAGGAAGA
GGCTCCGTGCAGAGGGGCCTGCGCGGTTCCAGGAGCAGCTCGGAGGGGCTCGGGCAGAGG
…
ACCCCAAAGATACAGATGCAATGAAACGCCGGGACACCTGCACCCCGATGTTTATAGCAG
CAATGGCCACGATAGCCAAATTGTGGAAGGAGCCTCGGTGTCCAACGAAAGATGAATGGA
TAAAGAAGATGTGGTTTATGTATACAATGGAATATTACTCAGCTATTAGAAATGACAAAT
ACCCACCATTTGCTTCAACGTGGATGGAACTGGAGGGTATTATGCTGAGTGAAGTAAGTC
AGTCGGAGAAGGACAAACATTATATGTTCTCATTCATTTGGGGAATATAAATAATAGTGA
AAGGGAATATAAGGGAAGGGGGAAGAAATGTGTGGGAAATATCAGAAAGGGAGACAGAAC
GTAAAGACTGCTAACTCTGGGAAACGAACTAGGGGTGGTGGAAGGGGAGGAGGGCGGGGG
GTGGGAGTGAATGGGTGACGGGCACTGGGTGTTATTCTGTATGTTAGTAAATTGAACACC
AATTAAAAAAAAAAAATTAAAAATAAAAAAAAATAAAAAAAAAAAAAGAGAGTAGAATGCT
AAGTGAAGTAAGCCAGTCAGAGAAAAACAAATACACATGAGTGATTTCACTCATAAGTCG
AATTTAAGAAACAAAACAAGCCAAAGAAAAGAGAGAGACAAACCAAGAAACAGACTCTCA
ACTATAGTGAACAAACTGATGGTTACCAGAGGGGAAGTGGGTGGGGGAAGGGGGATAATG
GGAATGGGGATTAAAGAGTATACTTATCATGATGAAAAAAAATAGTAATAATAAAGAGAA
ATAATGAAAAAGAAAAAAAAAAAAAAA**AAGATGTGA**ATAATAGAATATTACTCAGCCATA
AAAAAGAATATGATATATTACCAATTG

**Trans_13**
>Zoey chr14:30252401-30259294
GCTGGAGATAAGGCACAATGACATGTTAACATCTTCTATAT**AACAATTTAGCTAGA**GGGG
GAGGAGCAAGATGGCGGAAGAGTAGGGTCTCCAAATCACCTGTCTCCACCAAATTACCTA
GAAAACCTTCAAATTATCCTGAAAATCTATGAATTCGGCCTGAGATTTAAAGAGAGACCA
GCTGGAATGCAACAGTGAGAAGAGTTCGCGCTTCTATCAAGGTAGGAAGACGGGGAAAAA
GAAATAAAGGAACAAAGGCCTCCGAGGGGGAGGGGCCCCGCGAGGAGCCGGGCTGAGGCC
GGGGCGAGTGTCCCCAGGACAGGAGAGCCCCGTCCCGGAGACGCAGGAGCTGCACCGACC
TTCCCGGGCGGAAAGGGGCTCGCAGGGAGTTGGAGCAGGACCCAGAAGGAAGGGGATGCC
CTCGGGCTCCCCGGGTCAGTAACAGCAACTGCGCGCCCAGGAGAGTGCGCCGAGCTCCCT
AAGGGCTGCAGCGCGCACGGCGGGACCCGGCGGGACCCGGAGCAGCTGAAGGGGCTCGGG
CGGCGGCTCCGCGGAGGGGGCTGCGCGGCCCCGGGAGCAGCTCGGAGGGGCTCGGGCAGA
…
TGGGGGTGGTGGAAGGGGAGGAGGGCGGGGGGTGGGAGTGAATGGGTGACGGGCACTGGG
TGTTATTCTGTATGTTAGTAAATTGAACACCAATAAAAAAAAAAAAAAAAAAAAAAAAAA
AAAAAAAAAAAGAATGGCGCCTTTGGATCTTCTGTCCAAATCCGGCTTATTACAATGTCT
TAAGATTATAATTCATCAGGGCAACATATTTACTAGGTGAAAACTGACATGCTATTTGAG
AAATAGCTGTCATCTGCATCATCATGCACAAATTCTTTAAGTGTGCCTAAAGATCCAATC
TTATCATCTTCCCATCACTTGAACTTAACTAATAACATCAATCATTTTAAATATGTGCGG
GGGGATCCCTGGGTGGCTCAGTGGTTTAGTGCCTGCCTTCGGCCCAGGGCATGATCCTGG
AGTCCCGGGATCGAGTTCCCGCATCAGGTTCCCTGCATGGAGCCTGCTTCTCCCTCTGCC

TGTCTCTCTCTCGGTCTCTCATGAATAAATAAATAAAATATTTTGTAAATAAAAAAAAAA
AAAAAAAAAAAAAAAAAAAAAAAAAAAAAAAAAAAA**AACAATTTAGCTAGA**AACAAGTGGAA
TCATTTTCCTATCAAAAAATTCATTGAGATATATAAGACCTGTAGCCATAAACA

**Trans_14**
>Zoey chr15:23988119-23994782
TTCTAAGTTTTTTTCAAAAACTAGAGAATCTGTTATCATTAAAATTAAACTACTTTTGTA
TTTGACTTAAA**AAATAAAAAGTT**GGGGGGAGGAGCAAGATGGCGGAAGAGTAGGGTCTCC
AAATCACCTGTCTCCACCAAACTACCTAGAAAACCTTCAAATTATCCTGAAAATCTATGA
ATTCGGCCTGAGATTTAAAGAGAGACCAGCTGGAATGCAACAGTGAGAAGAGTTCGCGCT
TCTATCAAGGTAGGAAGACGGGGAAAAAGAAATAAAGAAACAAAGGCCTCCAAGGGGGAG
GGGCCCGCGAGGAGCCGGGCTGAGGCCGGGGCGAGTGTCCCCAGGACAGGAGAGCCCCGT
CCCGGAGGAGCAGGAGCTGCACCGACCTTCCCGGGGGAAAGGGGCTCGCGGGGAGCTGGA
GCAGGACCCAGGAGGGCGGGGATGCCCTCGGGCTCCCGGGGACACTAACAGCAACTGCGC
…
GCCACGATAGCCAAACTGTGGAAGGAGCCTCGGTGTCCAACGAAAGATGAATGGATAAAG
AAGATGTGGTTTATGTATACAATGGAATATTACTCAGCTATTAGAAATGACAAATACCCA
CCATTTGCTTCAACGTGGATGGAACTGGAGGGTATTATGCTGAGTGAAGTAAGTCAGTCA
GAGAAGGACAAACATTATATGTTCTCATTCATTTGGGGAATATAAATAATAGTGAAAGGG
AAAATAAGGGAAGGGAGAAGAAATGTGTGGGAAATATCAGAAAGGGAGACAGAACATAAA
GACTGCTAACTCTGGGAAACGAACTAGGGGTGGTAGAAGGGGAGGAGGGCGGGGGGTGGG
AGTGAATGGGTGACGGGCACTGGGGGTTATTCTGTATGTTAGTAAATTGAACACCAATAA
AAAAATAAATTAAAAAAAAAAATAAATGACATAATGTAAAAAAATAAAAATAAAAAAAAA
AAAATAAAGTCAACTGATTATGGGCATTAAGGAGGGCACGTGATATGATGAGCACTGTGT
GTTATATGTTGGCAAATTGAATTTAAGTTTAAAAAAACAATAAAATAAAATAATAAAAAT
AAACAACTAGAAAAAATAAAATAAAAAAAAAAAAATAAATAAAT**AAATAAAAAGTT**TAAA
GACACTTAAGGAGGTCACATGATATCATGAGACTGGGTGTTATATACAACAGATGAATAA
CTGA

**Trans_15**
>Zoey chr18:50946915-50953546
GTCAGGCTCCCTGCATGGAGCATGCCTCTCTCTCTACCTGTGTCTCTGCCTCTCTCTCTG
TATCTCTCATT**AATAAATAAATAAAATCT**GGGGGGAGGAGCAAGATGGCGGAAGAGTAGG
GTCTCCAAATCACCTGTCTCCACCAAACTACCTAGAAAACCTTCAAATTATCCTGAAAAT
CTATGAATTCGGCCTGAGATTTAAAGAGAGACCAGCTGGAATGCTACAGTGAGAAGAGTT
CGCGCATCTATCAAGGTAGGAAGACGGGGAAAAAGAAGTAAAGAAACAAAGGCCTCCAAG
GGGGAGGGGCCCGCGAGGAGCCGGGCTGAGGCCGGGGCGAGTGTCCCCAGGACAGGAGAG
CCCCGTCCCGGAGGAGCAGGAGCTGCACCGACCTTCCCGGGCGGAAAGGGGCTCGCGGGG
AGTTGGAGCAGGACCCAGGAGGGCGGGGATGCCCTCGGGCTCCCGGGGACAGTAACAGCA
ACTGCGCGCCCAGGAGAGTGCGCCGAGCTCCCTAAGGGCTGCAGCGCGCACGGCTTGACC
…
TGGCCACGATAGCCAAACTGTGGAAGGAGCCTCGGTGTCCAACGAAAGATGAATGGATAA
AGAAGATGTGGTTTATGTATACAATGGAATATTACTCAGCTATTAGAAATGACAAATACC
CACCATTTGCTTCAACGTGGATGGAACTGGAGGGTATTATGCTGAGTGAAGTAAGTCAGT
CGGAGAAGGACAAACATTATATGTTCTCATTCATTTGGGGAATATAAATAATAGTGAAAG
GGAAAATAAGGGAAGGGAGAAGAAATGTGTGGGAAATATCAGAAAGGGAGACAGAACGTA
AAGACTGCTAACTCTGGGAAACGAACTAGGGGTGGTAGAAGGGGAGGAGGGCGGGGGGTG
GGAGTGAATGGGTGACGGGCACTGGGTATTATTCTGTATGTTAGTAAATTGAACACCAAT
ACATCATGTCCAGGTGAAAAAAAAAAAAGAGTTCATCTTCTTGAAAATTATAAATAATTT
TCTTTAGTATAATCTTAAAAATTTGTAGCAGAAGTAATAAACACTTAGAATACTTTTTAA
AAATTCAGGGAATATAAGCTTAATAAAATAATAAATCTAAAGAAGAATAAGAGGATAACT

ATAAAAAAAAAATAAATAAATA**AATAAATAAATAAAATCT**TTTAAAAAATTGATCTTACA
TGAAAATTATTTTCAATGTCAAATTTTATGAC

**Trans_16a**

Trans_16a, Trans_16b, and Trans_16c share the same transduced sequence. Trans_16b, and Trans_16c are related by duplication in the Zoey assembly. Because this segment is not spanned by a single contig, it is not clear if this is a real segmental duplication or an assembly artifact.

```
>Zoey chr19:20170857-20175383
GGATGAGCATAGGAAGAGCTGTTGACATCTTTATGTTAGAAGACAATCCAAAATGATTGA
TAACATTATGAGAGTTCAAACTTTTTTTTTTTTTTTTTTTCTTTGTAATCTTTGTGATC
TTTGGTTACACTAAGGTTTCTTAAATAGTATACAAAAAGCATAAAATACAAATGAAACTT
TGTCTGATTGCTGAGGCTAGGACTTCCAGTACTATGTTGAACAGCAGTGGTGAGAGTGGA
CATCCCTGTCTTGTTCCTGATCTTAGGGGAAAGGCTCCCAGTGCTTCCCCATTGAGAATG
ATATTTGCTGTGGGCTTTTCATAGATGGCTTTTAAGATGTCGAGGAATGTTCCCTCTATC
CCTACACTCTGAAGAGTTTTGATCAGGAATGGATGCTGTATTTTGTCAAATGCTTTCTCT
GCATCCAATGAGAGGATCATATGGTTCTTGGTTTTTCTCTTGCTGATATGATGAATCACA
TTGATTGTTTTACGGGTGTTGAACCAGCCTTGTGTCCCAGGGATAAATCCTACTTGGTCA
TGGTGAATAATTTTTTTAATGTACTGTTGGATCCTATTGGCCAGTATCTTGTTGAGAATC
…
CAGCCCCTCCGCGGAGCCGCCGCCCGAGCCCCTTCAGCTGCTCCGGGTCCCGCCGGGTCC
CGCCGTGCGCGCTGCAGCCCTTAGGGAGCTCGGCGCACTCTCCTGGGCGCGCAGTTGCTG
TTACTGTCCCCGGAGCCCGAGGGCATCCCCGCCCTCCTGGGTCCTGCTCCAACTCCCCGC
GAGCCCCTTTCCCCCGGGAAGGTCGGTGCAGCTCCTGCTCCTCCGGGACGGGGCTCTCCT
GTCCTGGGACACTCGCCCCGGCCTCAGCCCGGCTCCTCGCGGGCCCCTCCCCCTTGGAGG
CCTTTGTTCCTTTATTTCTTTTTCCCCGTCTTCCTACCTTGATAGATGCGCGAACTCTTC
TCACTGTAGCATTCCAGCTGGTCTCTCTTTAAATCTCAGGCCGAATTCATAGATTTTCAG
GATAATTGAAGGTTTTCTAGGTAGTTTGGTGGAGACAGGTGATTTAGGGACCCTACTCTT
CCGCCATCTTGCTCCTCGAGTTCAAACTTTTGATCCCATTACACTTCATTCCCTTGCAGA
GCCACCTGACTCAGAATGGTCTGTACA
```

**Trans_16b**

```
>Zoey chr3:72514051-72518795
CCATCTTGTGTTTTCTTTCTTTCACTCATATAAATAAAGTCTTTATTAAAAAAGTCTAAT
TTGAGATTCCTTTTTTTTTTTTTTTTTTTTTTAGGAGAAATCTTTGTAATCTTTGTGATC
TTTGGTTACACTAAGGTTTCTTAAATAGTATACAAAAAGCATAAAATACAAATGAAACTT
TGTCTGATTGCTGAGGCTAGGACTTCCAGTACTATGTTGAACAGCAGTGGTGAGAGTGGA
CATCCCTGTCTTGTTCCTGATCTTAGGGGAAAGGCTCCCAGTGCTTCCCCATTGAGAATG
ATATTTGCGGTGGGCTTTTCATAGATGGCTTTTAAGATGTCGAGGAATGTTCCCTCTATC
CCTACACTCTGAAGAGTTTTGATCAGGAATGGATGCTGTATTTTGTCAAATGCTTTCTCT
GCATCCAATGAGAGGATCATATGGTTCTTGGTTTTTCTCTTGCTGATATGATGAATCACA
TTGATTGTTTTACGGGTGTTGAACCAGCCTTGTGTCCCAGGGATAAATCCTACTTGGTCA
TGGTGAATAATTTTCTTAATGTACTGTTGGATCCTATTGGCCAGTATCTTGTTGAGAATT
TTTGCATCCATGTTCATCAGGGATATTGGTCTGTAATTCTCCTTTTTGGCGGGGTCTTTG
TCTGGCTTTGGAATTAAGGTGATGCTGGCTTCATAGAACGAATTTGGAAGTACTCCATCT
CTTTCTATCTTTCCAAACAGCTTTAGGAGAATAGGTATGATTTCTTCTTTAAACGTTTGA
TAGAATTCCCCTGGGAAGCCATCTGGCCCTGGACTCTTGTGTCTTGGGAGGTTTTTGATG
ACTGCTTCAATTTCCTCCCTGGTTATTGGCCTGTTCAGGTTTTCTATTTCTTCCTGTTCC
…
ACCTCCCTGGAGCCCCTTTCCCCCGGGAAGGTCGGTGCAGCTCCTGCTCCTCCGGGACGG
GGCTCTCCTGTCCTGGGGACACTCGCCCCGGCCTCAGCCCGGCTCCTCGCGGGCCCCTCC
```

CCCTTGGAGGCCTTTGTTCCTTTATTTCTTTTTCCCCCGTCTTCCTACCTTGATAGATGC
GCGAACTCTTCTCACTGTAGCATTCCAGCTGGTCTCTCTTTAAATCTCAGGCCGAATTCA
TAGATTTTCAGGATAATTTGAAGGTTTTCTAGGTAGTTTGGTGGAGACAGGTGATTTAGG
GACCCTACTCTTCCGCCATCTTGCTCCTCCCCCCC**CTAATTTGAGATTCCT**AATGAAAAG
TCCCA

**Trans_16c**
The end of the insertion (5' of LINE-1) is disrupted by alteration of the apparent duplication in the Zoey assembly, however the 3' end contains the same transduced sequence as well as sequence matching the target site duplication found in Trans_16b.

>Zoey chr2:55188056-55192598
TTTCACTCATATAAATAAAGTCTTTATTAAAAAAGT**CTAATTTGAGATTCCT**TTTTTTTT
TTTTTTTTTTTTAGGAGAAATCTTTGTAATCTTTGTGATCTTTGGTTACACTAAGGTTTC
TTAAATAGTATACAAAAAGCATAAAATACAAATGAAACTTTGTCTGATTGCTGAGGCTAG
GACTTCCAGTACTATGTTGAACAGCAGTGGTGAGAGTGGACATCCCTGTCTTGTTCCTGA
TCTTAGGGGAAAGGCTCCCAGTGCTTCCCCATTGAGAATGATATTTGCGTGGGCTTTTCA
TAGATGGCTTTTAAGATGTCGAGGAATGTTCCCTCTATCCCTACACTCTGAAGAGTTTGA
TCAGGAATGGATGCTGTATTTTGTCAAATGCTTTCTCTGCATCCAATGAGAGGATCATAT
GGTTCTTGGTTTTTCTCTTGCTGATATGATGAATCACATTGATTGTTTTACGGGTGTTGA
…
CCGGGAAGGTCGGTGCAGCTCCTGCGTCTCCGGGACGGGGCTCTCCTGTCCTGGGGACAC
TCGCCCCGGCCTCAGCCCGGCTCCTCGCGGGGCCCCTCCCCCTTGGAGGCCTTTGTTCCT
TTATTTCTTTTTCCCCGTCTTCCTACCTTGATAGATGCGCGAACTCTTCTCACTGTAGCA
TTCCAGCTGGTCTCTCTTTAAATCTCAGGCCGAATTCATAGATTTTCAGGATAATTTGAA
GGTTTTCTAGGTAGTTTGGTGGAGACAGGTGATTTGGGGACCCTACTCTTCCGCCATCTT
GCTCCTCCCCCCCGATAACACCATTTTATTTCAACACTGTGCTGGAGGTTGTAGCCAGTG
TGGTCAGAAAAAAAATGAATAAAAAACATTCACAATGGAAAGA

**Trans_17**
>Zoey chr23:20368309-20375156
CACAAATATAGAAATTGGCAATAGATTACATTTTTCTCTGCAATAGGCAGAATCCTTCAT
T**GAAAAATAGAGTGAATC**GGGGGGAGGAGCAAGATGGCGGAAGAGTAGGGTCCCCAAATC
ACCTGTCTCCACCAAACTACCTAGAAAACCTTCAAATTATCCTGAAAATCTATGAATTCG
GCCTGAGATTTAAAGAGAGACCAGCTGGAATGCTACAGTGAGAAGAGTTCGCGCATCTAT
CAAGGTAGGAAGACGGGGAAAAAGAAATAAAGGAACAAAGGCCTCCAAGGGGGAGGGGCC
CCGCGAGGAGCCGGGCTGAGGCCGGGGCGAGTGTCCCCAGGACAGGAGAGCCCCGTCCCG
…
CAAACTGTGGAAGGAGCCTTGGTGTCCAACGAAAGATGAATGGATAAAGAAGATGTGGTT
TATGTATACAATGGAATATTACTCAGCTATTAGAAATGACAAATACCCACCATTTGCTTC
AACGTGGATGGAACTGGAGGGTATTATGCTGAGTGAAGTAAGTCAGTCGGAGAAGGACAA
ACATTATATGTTCTCATTCATTTGGGGAATATAAATAATAGTGAAAGGGAAAATAAGGGA
AGGGAGAAGAAATGGGTGGGAAATATCAGAAAGGGAGACAGAACGTAAAGACTGCTAACT
CTGGGAAACTAGGGGTGGTAGAAGGGGAGGAGGGCGGGGGGTGGGAGTGAATGGGTGACG
GGCACTGGGTGTTATTCTGTATGTTAGTAAATTGAACACCAATAAAAAAAAAATAAGAAA
TTTGGAAGGCAAAAAAAAAAAATAAAATAAAAAAAAATAAAAAAAAATAATAAAATAAAA
TAAAAAAAGAATAAAAATAAAAATAAAATTAAGTAACCTAGGGAAAACCTTTTCTACACT
GTTTGATCTATAGGAAAGTATCAATAAGTAACAACTATAATTATTTCTGTTAATAATATT
TGATTTTTAGAAAGCATTTAAAATGAAAAACATAGATCAGCAATTATTTCTAGTCTCTGG
AAGGGGTTGATGAGGTTAAGAAATTATTCTGTATAAATTGTAGATCTGGACTATACACTG
AGTCTTGTGGATCTTAATTAGAATAGCTTTTAATTCCTCAATTATGAATATGAATAATAA

AATTATATATAATACGGATAATAAAAAATAAAAAAATAAAAAAAAAAAGAAAAATAGAGT
GAATCAATGTTATATTCTCTTGGAACTTGTTTAATCATGTATAATATTAAGGCGAAGTCA
AACCTGAC

**Trans_18**
>Zoey chr27:10823103-10827419
TTACACTCATATAAAGCTAGCCAGTGGTCCTGTAATTGATC**AGAAGACTTTTCTT**TTTTT
TTTTTTTTTTTTTTTTTTTTATTTTGTAATATCTATATTTTGAACACTGACATCATTAT
ACTATGTGAAATTTTGCTAGTAATACAATACATGGTCCTTGTAAATATATATGTTTCCTT
TATAAGATTTAAATTGGAGTAGCATGATAAGAAGTCATGAAATGAAAGTACTTAAAATCA
AAGTAATGTTTATCACATTGATATCATTACCTGACTTAGAAACTGTGTTGTTTGGCATAT
GTGAAATTTTGTTTCAAAACTAAACTTTTTTTTTTTTTTTTTTGTAAGGAAAGAAAAAAC
TTATATTTATACAAAAATATTGGTGTTCAATTTACTAACATACAGAATAATACCCAGTGC
CCGTCACCCATTCACTCCCACCCCCGCCCTCCTCCCCTTCTACCACCCCTAGTTCGTTTC
CCAGAGTTAGCAGTCTTTACGTTCTGTCTCCCTTTCTGATATTTCCCACACATTTCTTCT
CCCTTCCCTTATTTTCCCTTTCACTATTATTTATATTCCCCAAATGAATGAGAACATATA
ATGTTTGTCCTTCTCCGACTGACTTACTTCACTCAGCATAATACCCTCCAGTTCCATCCA
…
TGGATGTAAAGTTCTGTAGATATCTGTGAAATCCATCTGGTCTAGTGTATCATTTAAAGC
TCTCGTTTCTTTGGAGATGTTTTGCTTAGAAGACCTATCGAGTATAGAAAGAGCTAGATT
GAAGTCACCAAGTATAAGTGTATTATTATCTAAGTATTTCTTCACTTTGGTTAATAATTG
ATTTATATATTTGGCAGCTCCCACATTCGGAGCATATATATTGAGGATTGTTAAGTCCTC
TTGTTGAATAGATCCTTTAAGTATGATATAGTGTCCCTCTTCATCTCTCACTACAGTCTT
TGGGGTAAATTTTAGTTTATCTGATATAAGGATGGCTACCCCTGCTTTCTTTTGAGGACC
ATTCGAATGGTAAATGGTTCTCCAACCTTTTATTTTCAGGCTGTAGGTGTCCTTCTG**AGA**
**AGACTTTTCTT**AATTGCTACCTAGAGTGAAGGGCAAGAAAGGCAATTTTCCTTGCAG

*Comparison of full-length L1_Cf elements in Zoey and CanFam3.1*

The L1_Cf consensus sequence is 6327 bp in length and contains homopolymer runs and segments of GC-rich sequence. These features include a stretch of seven cytidine residues in the ORF1p coding sequence. Such homopolymeric sequences are prone to replication errors and could be difficult to resolve, particularly because indels are the major source of errors encountered in PacBio sequencing data (66). Moreover, these types of errors are rarely corrected with Illumina short-read sequencing data due to the absence of reads with unique alignments to the internal portions of LINE-1 sequences. As such, we believe the number of L1_Cf elements containing intact open-reading frames is underestimated in the Zoey Pac Bio assembly. To compare the relative content of the Zoey and CanFam3.1 assemblies, we tabulated the number of L1_Cf elements that are full-length or nearly full-length.

As illustrated in Table 5.5, dimorphic canine LINEs are assigned to disparate families by RepeatMasker. This assignment is based solely on alignment percent identity, without regard to the presence of specific markers which are diagnostic to specific LINE sub-families. Thus, we searched the Zoey and CanFam3.1 assemblies for matches to the L1_Cf consensus sequence using RepeatMasker with the custom library (-lib) setting. As shown in Table 5.9, the Zoey assembly contains a larger number of long L1_Cf sequences exhibiting low sequence divergence when compared to the CanFam3.1 assembly. This number is further increased when we analyze the Zoey secondary contigs, which likely reflect the presence of dimorphic LINE-1s in the diploid Zoey genome. Additional data is required to determine the extent to which these differences reflect different levels of completeness for the two assemblies or real differences in L1_Cf content among dog breeds.

| Criteria | CanFam3.1 | Zoey Primary Assembly | Zoey Secondary Contigs | Zoey Total |
|---|---|---|---|---|
| >= 5 kbp | 3,091 | 3,195 | 354 | 3,549 |
| >= 5 kbp, <= 5% divergence | 2,031 | 2,263 | 308 | 2,571 |
| >= 5 kbp, <= 2% divergence | 922 | 1,124 | 218 | 1,342 |
| >= 6.290 kbp | 1,124 | 1,352 | 204 | 1,556 |
| >= 6.290 kbp, <= 2% divergence | 600 | 837 | 169 | 1,006 |
| 6.327 kbp, <= 2% divergence | 113 | 156 | 31 | 187 |

**Table 5.9: Summary of L1_Cf elements found in genome assemblies.** The counts of sequences that satisfy the indicated criteria are given for the CanFam3.1 reference genome, the Zoey genome assembly, the Zoey secondary contigs, and the total found in Zoey.

### 6. Mobile Element Analysis

*Identification of candidate intact LINE-1 insertions*

Prior to finalizing the assembly, we used a previously described pipeline (14) to identify candidate intact LNE-1 insertions that are present in Zoey and absent from the CanFam3.1 assembly. First, Illumina short-read data was aligned to the CanFam3.1 assembly and candidate LINE-1 insertions were identified using the program Retroseq (67). We excluded candidates within 500 bp of another LINE-1 sequence annotated in the reference genome. Assembly of the insertion supporting reads for each putative site was performed as described previously (14). Given the size of a full-length LINE-1 insertion relative to the length of the Illumina sequencing reads, this process is expected to result in two contigs representing the 5' and 3' insertion junction sequences, respectively. The assembled contigs were analyzed by RepeatMasker to identify LINE-1-containing assemblies. Candidate full-length calls were identified as having separate contigs that respectively contained: (i) assembled 5′ and 3′ L1_Cf derived edges, (ii) ≥30 bp of both LINE-derived and mappable genomic sequence at either breakpoint, and (iii) the presence of matching target site duplications flanking the assembled 5' and 3' junctions. Candidate sites with contigs satisfying these criteria were aligned to the CanFam3.1 reference to identify the chromosomal positions of each insertion and compared to the map of clones identified in sequenced fosmid pools.

*LINE-1 subcloning strategy*

Individual fosmid clones containing putative full-length LINE-1 calls in Zoey were isolated and assembled using long-read PacBio sequencing. Sequences were obtained from three fosmids that contained full-length LINE-1s (Figure 6.1). Sequence analysis revealed that fosmid clone 104_5, located on chr1, had intact copies of both ORF1 and ORF2 without obvious amino acid substitutions that would adversely affect their respective functions.

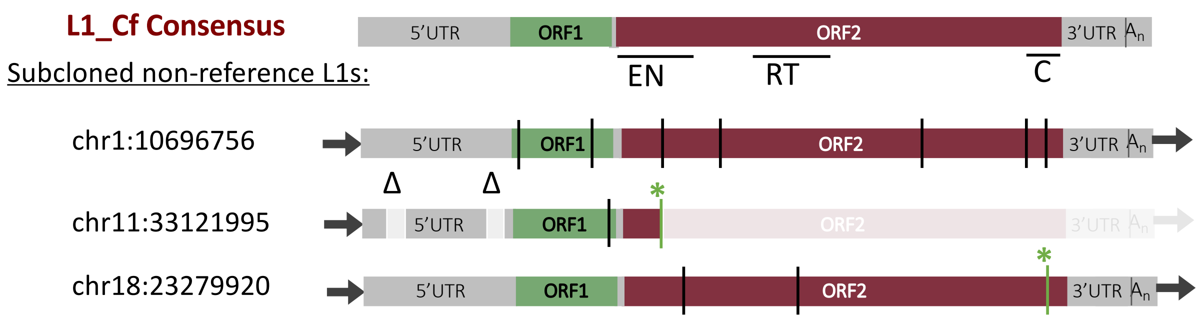


**Figure 6.1: Schematics of full-length LINE-1s.** We initially isolated three different full-length LINE-1s from fosmid clones. The structure of each LINE-1 was compared to the L1_Cf consensus sequence. The cartoon in the top row indicates the structure of the L1_Cf consensus. The colored rectangles denote the 5' LINE-1 untranslated region (UTR), ORF1, ORF2, and 3' UTR segments, respectively. The approximate positions of the endonuclease (EN), reverse transcriptase (RT), and cysteine-rich carboxyl terminal domain (C) within ORF2 are indicated by horizontal lines. The sequence structure of three non-reference LINE-1s is shown below. Black vertical lines indicate nonsynonymous sequence changes, vertical green lines indicate sequence changes that result in stop codons (green asterisk), and triangles indicate deletions. The LINE-1 insertion on chr1 is predicted to contain intact open reading frames for both ORF1 and ORF2.

We next tested whether the full-length canine LINE-1 sequence were capable of retrotransposition using an established cultured cell-based assay (8, 68). Briefly, the candidate full-length canine LINE-1 sequence was subcloned into an episomal mammalian expression vector (pCEP4) equipped with a retrotransposition indicator cassette (*mneoI*). The strategy used to create the canine LINE-1 expression constructs required several steps, took advantage of unique restriction sites located within the 5'-UTR (*Eco*RI) and 3'-UTR (*Dra*III) of the L1_Cf consensus sequence (RepBase release 21.06), and is summarized below (Figure 6.2).

Step 1: To create the pCR-L1Cf-Empty plasmid (Figure 6.2), we modified the pCR Blunt vector (Invitrogen). PCR was used to introduce a sequence that contains: (i) a *Not*I restriction site at its 5' end; (ii) the 101 first nucleotides of the L1_Cf-5'UTR consensus sequence (which includes the unique *Eco*RI site); (iii) a stretch of four adenosine residues; (iv) 18 nucleotides of the L1_Cf-3'UTR consensus sequence (which begins at the *Dra*III site); and (v) a *Psi*I restriction site at its 3' end. Positive clones initially were identified using restriction mapping; Sanger DNA sequencing confirmed the integrity of the sequence.

Step 2: To create pCR-L1Cf-104-5 (as well as other canine LINE-1 vectors), we digested fosmid DNA from Zoey containing the candidate full-length canine LINE-1 sequence with *Eco*RI and *Dra*III. The liberated ~6263 bp restriction fragment was subcloned into the *Eco*RI and *Dra*III cleaved pCR-L1Cf-Empty plasmid. Positive clones initially were identified using restriction mapping; Sanger DNA sequencing confirmed the integrity of the sequence.

Step 3: To create the canine LINE-1 episomal expression vectors, the candidate full-length canine LINE-1 containing vectors (*e.g.*, pCR-L1Cf-104-5) were digested with *Not*I and *Psi*I. The liberated ~6.26 kbp DNA fragment was used to replace an active human LINE-1 sequence present in the pJM101/L1.3 expression plasmid (69, 70). Briefly, pJM101/L1.3 plasmid DNA was cleaved with *Not*I and *Bst*Z17I and the band corresponding to the ~12.3 kbp pCEP4 plasmid DNA backbone containing the *mneoI* retrotransposition indicator was gel purified. Ligating the ~6.26 kbp *Not*I and *Psi*I restriction fragment containing the full-length canine LINE-1 and the ~12.3 kbp pCEP4 plasmid DNA containing the *mneoI* retrotransposition indicator cassette created the canine LINE-1 expression plasmids shown in Figure 6.2. Positive clones initially were identified using restriction mapping; Sanger DNA sequencing confirmed the integrity of the sequence.

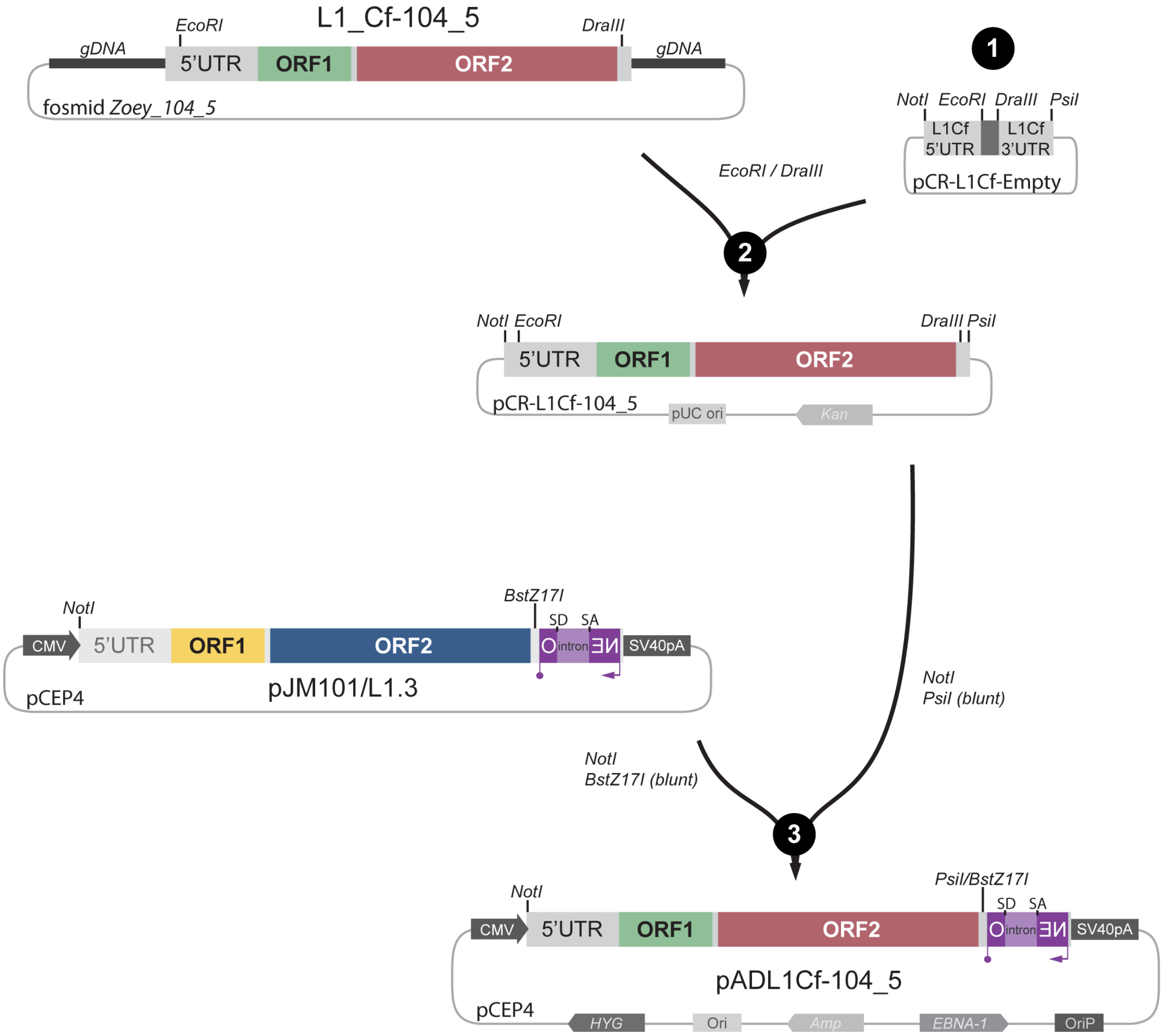


**Figure 6.2: LINE-1 subcloning strategy.** Schematic representation of the steps required to create canine LINE-1 expression plasmids containing a retrotransposition indicator cassette. Details are noted in the above text.

*SINEC_Cf subcloning strategy*

To assay the capability of the canine LINE-1 to act as a driver for SINEC retrotransposition *in trans*, the consensus sequence of a tRNA-derived SINEC_Cf element was equipped with a *neo*$^{tet}$ retrotransposition indicator cassette that can monitor the detection of RNA polymerase III (pol III) transcripts. To accomplish this task, we modified the Alu *neo*$^{tet}$ vector (10), which contains a 7SL RNA PolIII enhancer sequence followed by an active copy of a human Alu element equipped with a "backward" copy of a neomycin resistance gene whose expression is driven by an SV40 promoter. The *neo* gene is disrupted by a self-splicing tetrahymena group I intron that is in the same transcriptional orientation as the Alu element. The strategy used to create the canine SINEC_Cf is summarized below and is described in Figure 6.3.

Notably, this strategy can be used to readily test individual SINEC-Cf elements for their ability to retrotranspose in future studies.

Step 1a: PCR was used to generate a DNA fragment containing: (i) a *Sal*I restriction site near its 5' end; (ii) the 7SL RNA polymerase III enhancer followed by a *Not*I restriction site; (iii) an engineered *Age*I restriction site upstream of the *neo*$^{tet}$ indicator cassette; and (iv) a fragment of the *neo*$^{tet}$ indicator cassette that includes a *NgoM*IV restriction site.

Step 1b: To create pADSINE-Empty, the abovementioned PCR product was digested with *Sal*I and *NgoM*IV and the cleaved product was used to replace the *Sal*I to *NgoM*IV site in the Alu *neo*$^{tet}$ vector (10). Positive clones initially were identified using restriction mapping; Sanger DNA sequencing confirmed the integrity of the sequence.

Step 2a: Overlapping oligonucleotides (SINE_Cf_FULLs and SINE_Cf_FULLas) were used in PCR reactions to generate a SINE_Cf consensus sequence (obtained from Repbase) (9) that was flanked by unique *Not*I and *Age*I restriction sites.

Step 2b: To create the pADSINEC_Cf-RB expression vectors, the abovementioned PCR product was cleaved with *Not*I and *Age*I and the resultant fragment was introduced into the *Not*I and *Age*I pADSINE-Empty vector. Positive clones initially were identified using restriction mapping; Sanger DNA sequencing confirmed the integrity of the sequence.

The primer sequence used to reconstruct the SINEC_Cf consensus sequence are listed below.

*SINEC_Cf_FULLs*
5'-
acc**gcggccgc**GGGATCCCTGGGTGGCGCAGCGGTTTGGCGCCTGCCTTTGGCCCAGGGCGCGA
TCCTGGAGACCCGGGATCGAATCCCACGTCGGGCTC-3'

*SINEC_Cf_FULLas*
5'-
ggg**accggt**ATTTATTTATGATAGTCACACAGAGAGAGAGAGAGAGGCAGAGACACAGGCAG
AGGGAGAAGCAGGCTCCATGCACCGGGAGCCCGACG-3'

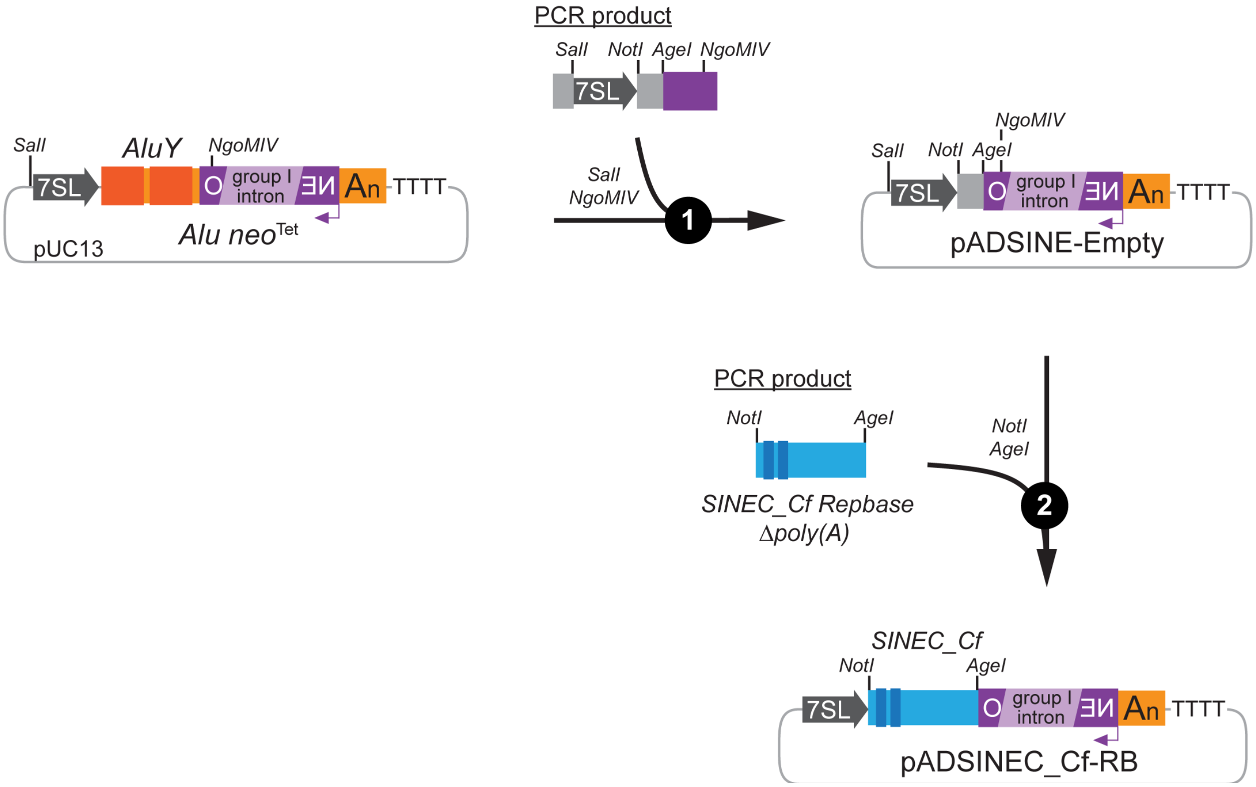


**Figure 6.3: SINEC_Cf consensus subcloning strategy.** Schematic representation of the steps required to create canine SINEC_Cf expression plasmids containing a retrotransposition indicator cassette. Details are noted in the above text.

*Cultured cell retrotransposition assay*

Cis-based LINE-1 retrotransposition assays

HeLa-HA cells were maintained as described previously (68). Briefly, HeLa-HA cells were cultured in complete media consisting of MEM ([-] L-Glutamine) tissue culture medium supplemented with 10% fetal bovine serum, 20U/mL Penicillin/Streptomycin, and 1x GlutaMax (GIBCO) at 37°C in 5.0% $CO_2$. Cells grown to ~80% confluency were washed in 1x phosphate buffered saline solution (PBS) and harvested in 0.25% Tryspin-EDTA (GIBCO). The cells were resuspended by repeated pipetting in complete media. Approximately 2 x $10^3$, 2 x $10^4$, or 2 x $10^5$ cells were plated in each well of a 6-well tissue culture plate and cultured for 24 hours. The candidate full-length canine LINE-1 constructs (*e.g.*, ADL1Cf-104_5), containing the *mneo*I cassette, as well as the human LINE-1 expression constructs JM101/L1.3 (L1Hs) and JM105/L1.3 (L1Hs RT-), which were used as positive and negative controls, respectively, then were transfected into the HeLa-HA cells. Briefly, 3 μg of each construct was incubated for 20 minutes in the presence of 291μL pre-warmed Opti-MEM medium (GIBCO) and 9μL Fugene 6 transfection reagent (Promega). Three wells of each cell concentration then were transfected with 100μL of each transfection mixture. The media was replaced with fresh, pre-warmed complete media 24 hours post-transfection. After 72 hours, G418 selection began by growing the cells in complete media supplemented with 400μg/mL G418 (GIBCO). The G418-containing media was replaced daily during the duration of the experiment (typically 12 to 14 days). The tissue culture plates then were washed using 1x PBS, fixed using a 2% formaldehyde plus 0.2% glutaraldehyde solution prepared in 1x PBS, and stained with an aqueous 0.1% crystal violet solution to visualize the resultant cell foci. In these experiments, HeLa-HA foci were suitable for counting at the 2 x $10^4$ cell dilution. We then reported the canine LINE-1 retrotransposition efficiencies relative to the positive (L1Hs) and negative (L1Hs RT-) controls (68).

*Trans*-based SINEC retrotransposition assay

HeLa-HA cells were plated at an approximate concentration of 4 x $10^5$ cells per well of a 6-well tissue culture plate and were cultured for 24 hours prior to co-transfection (11). Co-transfections were performed with equal amounts of the ADL1Cf-104_5Δ*neo* construct (L1_Cf), a derivative of ADL1Cf-104_5 that lacks the *mneoI* reporter, and the SINEC_Cf-*neo*$^{tet}$ construct ADSINEC_Cf-RB. Separate co-transfections of JM101/L1.3 Δ*neo* (L1Hs) or JM105/L1.3Δ*neo* (L1Hs RT- (70, 71)) with equal amounts of *Alu neo*$^{tet}$ were used as positive and negative controls, respectively. Co-transfections were performed in triplicate. SINEC_Cf and L1CfΔ*neo* plasmids (1.5 μg each) were mixed and incubated for 20 minutes in 291 μL pre-warmed Opti-MEM medium (GIBCO) and 9 μL of the Fugene HD transfection reagent (Promega); 100 μL of the resultant mixture was transferred to each of three wells in the 6-well tissue culture plates. The media was replaced with fresh, pre-warmed complete media 24 hours post-transfection. After 72 hours, G418 selection began by growing the cells in complete media supplemented with 400μg/mL G418. After 12-14 days post-transfection cells were washed, fixed, and stained as described above.

### 7. Genome Browser TrackHub Resources

*Accessing the TrackHub*
A UCSC TrackHub hosting the Zoey assembly, as well as relevant annotations of both the Zoey assembly and CanFam3.1, is hosted at: https://github.com/KiddLab/zoey_genome_hub. The TrackHub can be loaded into the UCSC track hub (72) by pasting the following link into the 'My Hubs' tab at: http://genome.ucsc.edu/:
https://raw.githubusercontent.com/KiddLab/zoey_genome_hub/master/zoey-genome.hub.txt .

*Summary of available data*
Annotation tracks for CanFam3.1

*Deletions, Insertions, Inversions, Complex_SV*: These tracks give the coordinates of structural variants identified between the CanFam3.1 and Zoey assemblies.

*Secondary_del, Secondary_ins*: These tracks give the coordinates of structural variants identified between the CanFam3.1 assembly and the alternative haplotypes assembled from Zoey.

*CHORI-82, CHORI-82 discordant, CHORI-82 large discordant*: These tracks give a clone map of end-sequence pairs from the CH82 BAC library aligned to the genome using bwa mem. Tracks are shown separately for concordant clones, clones with an aberrant mapping, and clones with an aberrant mapping that spans more than 5 Mbp. For the discordant clones, clones that appear to be too large (indicting a deletion relative to the genome reference) are colored in red, clones that appear too small (indicating an insertion relative to the reference) are colored in blue, clones with an orientation suggestive of an inversion are colored in green, clones with an everted orientation suggestive of a new tandem duplication are colored in orange, and clones whose end sequences map to different chromosomes are shown in purple.

*Fosmid, fosmid_discordant, fosmid_large_discordant*: Same as above but for the Tasha fosmid library downloaded from the NCBI Trace Archie as described in Section 1.

*Gaps_filled*: This track gives the location of gaps in the CanFam3.1 assembly which are confidently filled in by sequence in the Zoey genome. The name of each element reflects the corresponding coordinates in the Zoey genome.

*SegDup*: This track gives the coordinates of segmental duplications identified by genome self-alignment using SEDEF (42). The track is color coded based on the sequence identity of the duplicated segments:

90-92% very light gray
92-94% light gray
94-96% medium gray
96-98% dark gray
98-99% yellow
>99% orange

*ZoeyFASTCN*: This track reports copy-number estimated on the CanFam3.1 assembly based on Illumina short-read sequencing data from Zoey that was processed using the fastCN pipeline (12).

*Zoeylift*: This track summarizes the liftOver comparison between the canFam3.1 and Zoey assembly. Segments that liftOver to the same chromosome are colored in red, segments that align to the same chromosome but in an inverted orientation are colored in green, segments that correspond to different chromosomes are colored in purple.

Annotation tracks for Zoey

*Ensembl cDNA BLAT*: This track shows Ensembl gene models aligned to the Zoey genome using BLAT.

*RefSeq Mammal*: This track shows RefSeq mammalian gene models aligned to the Zoey genome using BLAT

*Unfiltered genes, filtered genes*: These tracks depict the total and final filtered set of predicted protein-coding gene models described in Section 2 of this note.

*Filtered expression*: This track shows the expression level estimated for each of the filtered genes across tissues. See Section 2 of this note for additional information.

*lncRNA predictions*: This track shows long-noncoding RNAs predicted in the Zoey genome.

*Assembly gaps*: This track indicates gaps in the Zoey assembly

*Complex_SV, deletions, insertions*: As indicated, these tracks show the filtered structural differences between the Zoey and CanFam.1 assemblies.

*Inversions:* This track shows the filtered set of candidate inversions

*Secondary_deletions, secondary_insertions*: These tracks show the position of insertions and deletions between the Zoey primary assembly and the Zoey alternative contigs.

*TRF*: This track shows tandem repeats identified by tandem repeat finder (40).

*CanFam Gap*: This track shows locations of gaps in CanFam3.1 that have been precisely covered in the Zoey assembly

*CanFam_lift*: This track shows the liftOver alignment between the Zoey and CanFam3.1 assemblies. The color code is as described for the corresponding track above.

*CH82_concordant, CH82_discordant, CH82_largediscordant*: These tracks depict mapping of CHORI-82 BAC clones to the Zoey genome. Coloring is as described for the corresponding track above.

*Contigs_map*: This track shows the positions of original assembled contigs aligned to the assembly using minimap2 (43).

*GC_content*: This track depicts the GC content of the assembly in sliding windows.

*Scaffmap_merge*: This track shows the locations of the scaffolds of the final Zoey assembly as mapped using minimap2 (43).

*SegDup*: This track shows the location of segmental duplications in the Zoey assembly based on genome self-alignment. The color is as described for the corresponding track above.

*Tasha fastCN, Zoey fastCN*: These tracks show copy-number estimated from read data (Illumina data for Zoey, Sanger data form WGS plasmid library for Tasha) aligned against the Zoey assembly.

*WindowMasker_and_DUST*: This track shows the parts of the assembly that are masked by the WindowMasker and DUST pipelines (41).

*RepeatMasker*: This track shows common repeats sequences in the genome annotated using RepeatMasker (39).

Other data
Additional data is available on the Zoey Genome gitHub page at: https://github.com/KiddLab/zoey_genome_hub/tree/master/supporting-data. This includes the chain files required to use liftOver to convert between the coordinates of CanFam3.1 and the Zoey genome as well as the assembled gene sequence of the final and filtered gene sets created using the gene annotation process described in Section 2.

Example browser views, depicting a region that is inverted between the CanFam3.1 and Zoey assemblies, are shown in Figures 7.1 and 7.2.

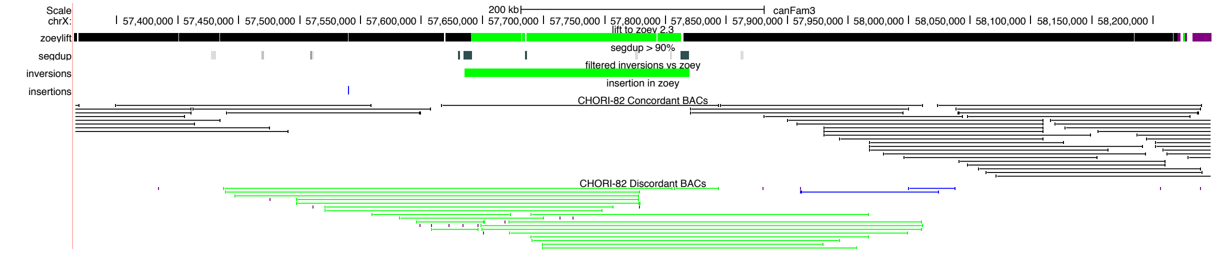


**Figure 7.1:** Annotated tracks for the CanFam3.1 assembly. Shown is an example region on the X chromosome. The top track illustrates alignment of segments of the CanFam3.1 assembly against the Zoey genome. The green region in the middle shows several segments that align in an inverted orientation. This region is flanked by sequence which is duplicated in the CanFam3.1 assembly (SegDup track). This region meets the criteria to call a candidate inversion, as shown by the green bar in the inversion track. Interestingly, concordant clone support from the CH-82 BAC library is lower in this region (black bars), and there is a cluster of BACs from this library (in green) that show a pattern consistent with an inversion relative to the assembly.

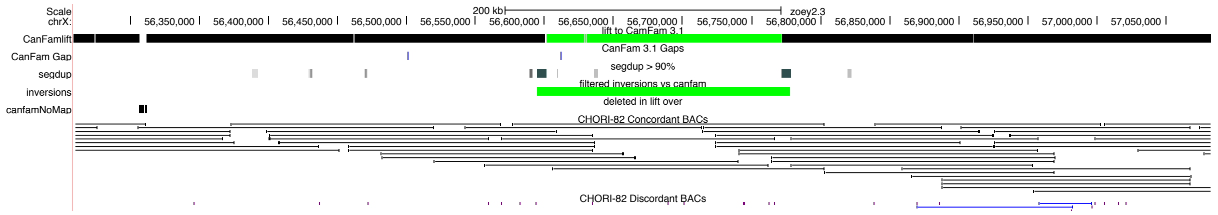


**Figure 7.2: Annotated tracks for the Zoey assembly.** The region corresponds to the same locus shown in Figure 7.1 Note that in the Zoey assembly the CH-82 Tasha BAC library shows concordant mapping across the predicted inversion (green bar).
